# Comprehensive interaction profiling and machine learning prediction of bacteriophage infectivity across clinically diverse *Pseudomonas aeruginosa*

**DOI:** 10.64898/2026.05.19.726084

**Authors:** Denish Piya, Avery J.C. Noonan, Hemaa Selvakumar, Mohamad Alayouni, Sarshad Koderi Valappil, Flavien Maucourt, Isabella Murray, Madeline Svab, Collis Bousliman, Matthew Heidenblut, Britney Oriheula, Alexey Kazakov, Hans K. Carlson, Yiyan Yao, Emerson Smith, Simon Roux, Adam M. Deutschbauer, Jamie L. Inman, Adam P. Arkin, Vivek K. Mutalik

**Affiliations:** Environmental Genomics and Systems Biology, Lawrence Berkeley National Laboratory, Berkeley, CA, USA; Molecular and Cell Biology Department, University of California Berkeley, Berkeley, CA, USA; Plant and Microbial Biology Department, University of California Berkeley, Berkeley, CA, USA; Department of Bioengineering, University of California Berkeley, Berkeley, CA, USA; Biological Systems and Engineering, Lawrence Berkeley National Laboratory, Berkeley, CA, USA; DOE Joint Genome Institute, Lawrence Berkeley National Laboratory, Berkeley, CA, USA

## Abstract

The rise of antibiotic-resistant bacterial infections has driven renewed interest in bacteriophage therapy, where viruses that specifically kill bacteria are used as targeted antimicrobials. *Pseudomonas aeruginosa*, a WHO critical-priority pathogen that causes severe infections in hospitalized and immunocompromised patients, presents a major challenge for phage therapy because of its extraordinary genetic diversity. Phages effective against one bacterial strain often fail against others, and existing cross-resistance-profiling approaches require iterative empirical testing of each new patient isolate. To establish a genome-based framework for rapid phage-isolate matching, we assembled a collection of 95 genomically diverse *P. aeruginosa* phages representing 20 genera and tested each against 99 genetically diverse clinical isolates, generating 9,405 infection outcome measurements. Bacterial O-antigen serotype emerged as the dominant determinant of strain susceptibility, while defense systems, anti-defense systems, and prophage burden contributed smaller strain-specific effects. The full curated multivariate model explained 47% of strain-susceptibility variance. Machine-learning models integrating these features and pangenome-derived gene clusters reached a per-strain AUROC of 0.86. In an *in vivo* proof-of-concept test against a single held-out strain, the ML-designed cocktail produced a ∼12-fold greater median CFU reduction than the expert-designed cocktail (q = 0.045), with both cocktails substantially reducing burden relative to the untreated control (∼113-fold for ML, ∼9-fold for CG; both q < 10□³). SHAP analysis of the model identified bacterial surface-architecture genes (LPS biosynthesis, outer membrane proteins, type IV pili) as the dominant predictors, with defense-system content modulating which specific phages succeed against a strain rather than uniformly damping susceptibility. Together, these results establish a genome-based framework for predicting phage susceptibility in genetically diverse clinical isolates.

## Introduction

Antimicrobial-resistant *Pseudomonas* aeruginosa is a WHO critical-priority pathogen responsible for severe nosocomial infections worldwide, with multidrug resistance rates continuing to rise across geographic regions^1,2,3,4–6^. Its combination of intrinsic resistance mechanisms and capacity for horizontal gene transfer renders conventional antibiotic development increasingly ineffective against this organism^7^. Bacteriophages offer a complementary therapeutic strategy. Their strain-specific targeting, ability to replicate at infection sites, and potential to be engineered for enhanced activity make them promising candidates for treating antibiotic-resistant *P. aeruginosa* infections^8–10^. However, deploying phage therapeutically requires systematic knowledge of phage-host interactions at a scale that ad hoc isolation-and-testing workflows cannot deliver.

Phage therapy faces a structural challenge that antibiotics do not. Phages exhibit strain-level specificity determined by bacterial surface architecture^11,12^. Different surface components, including lipopolysaccharide, flagella, pili, and outer membrane porins, serve as receptors for phage attachment, and these structures can vary considerably even among strains of the same species ^13,14^. Whole-genome sequencing has revealed that *P. aeruginosa* exhibits remarkable genomic diversity, with individual strains differing by hundreds to thousands of genes despite belonging to the same species^15–20^. For most dsDNA phages, infection initiation depends on compatible interactions between bacterial surface receptors and phage receptor-binding proteins located in tail fibers or tail spikes^21,22^. Additional factors, including chromosomal antiphage defense systems and prophage-encoded immunity modules, further shape outcomes but are difficult to predict from genomic features alone^21,23,24^. Consequently, a phage infecting one isolate may fail to infect another despite close genomic relatedness, complicating the development of broadly effective phage therapeutics.

Bacteria can evolve resistance to phages through modification, loss, or altered regulation of their surface receptors. To circumvent this resistance mechanism, phage cocktails include multiple phages that target independent receptors, reducing the likelihood that a single bacterial mutation will confer cross-resistance^25–30^. Systematic selection of phages through cross-resistance profiling has yielded cocktails utilizing non-redundant receptors that are active against more than 96% of clinical *P. aeruginosa* isolates^31^; receptor-guided rational design has produced a five-phage cocktail with 76% infectivity across diverse MDR clinical isolates that showed efficacy in a murine wound infection model^32^. These approaches achieve high coverage but require empirical characterization of each phage against reference strains or mutant panels, a per-isolate empirical workflow that does not scale to clinical timeframes. Machine learning offers a complementary path. Models trained on large-scale, genomically diverse phage-host interaction datasets can predict the susceptibility of novel *P. aeruginosa* isolates to characterized phage banks from genome sequence alone, reducing the per-isolate empirical testing required by existing cross-resistance and receptor-guided workflows. Once trained, the same model can be reused for many patient isolates, enabling rapid phage selection in clinical settings.

Despite decades of *P. aeruginosa* phage research, systematic characterization of phage-host interactions across representative strain collections has been limited^28,33^. Most published studies have used reference strains such as PAO1 and PA14 or small convenience samples of locally available clinical isolates, and have typically tested either many phages against a few strains or many strains against a few phages^34–38^. Without a curated strain collection representing the diversity of clinical isolates across different sequence types, accessory genome content, geographic origins, and infection sources, our understanding of phage host-range determinants risks being biased toward laboratory-adapted lineages^39,40^. This data gap directly constrains predictive modeling, as models trained on narrow strain collections risk learning correlations specific to those lineages rather than principles governing phage-host compatibility broadly^39,55,58^.

Here, we assembled 99 genomically diverse clinical *P. aeruginosa* strains and 95 phages representing 20 genera, profiled all 9,405 pairwise interactions, dissected the genomic determinants of compatibility using multivariate analyses, and trained machine-learning models that predict susceptibility for held-out strains from genome sequence alone. We then used these models to design a three-phage cocktail and compared its *in vivo* efficacy against an expert-designed cocktail in a murine wound infection model targeting a strain that was withheld from feature selection and model training. The integrated approach establishes a scalable framework for matching phages with genetically diverse clinical *P. aeruginosa* isolates.

## Results

### Assembly of genetically diverse bacterial strains and phage collections for large-scale interaction profiling

To develop robust machine learning models and a comprehensive view of phage-host interaction networks in *P. aeruginosa*, we assembled matched collections of bacterial strains and phages that together capture the genomic diversity relevant to clinical phage therapy. For the bacterial panel, we used 96 strains from the 100-strain MRSN (Multidrug Resistant Organism Repository and Surveillance Network) diversity panel, which was curated from 3,785 clinical *P. aeruginosa* isolates collected globally^15^. In addition, we supplemented the collection with three laboratory reference strains (PAO1, PA14, and PUPa3) for a total of 99 strains (*Fig. 2A*). This panel spans 91 sequence types, including epidemic clones ST-111, ST-235, ST-244, and ST-253, and exhibits substantial variation in phage-relevant host systems and structures, including prophage content (0 to 4 complete prophages per strain) (*Fig. 2A / Dataset S1*), antiphage defense systems (2 to 22 systems per strain) (*Supp. Fig. 1*), O-antigen serotypes (13 distinct types) (*Supp. Fig. 2*), and antibiotic resistance profiles, reflecting the breadth of genomic diversity present in clinical populations (*Supp. Fig. 3*).

**Fig. 1.**
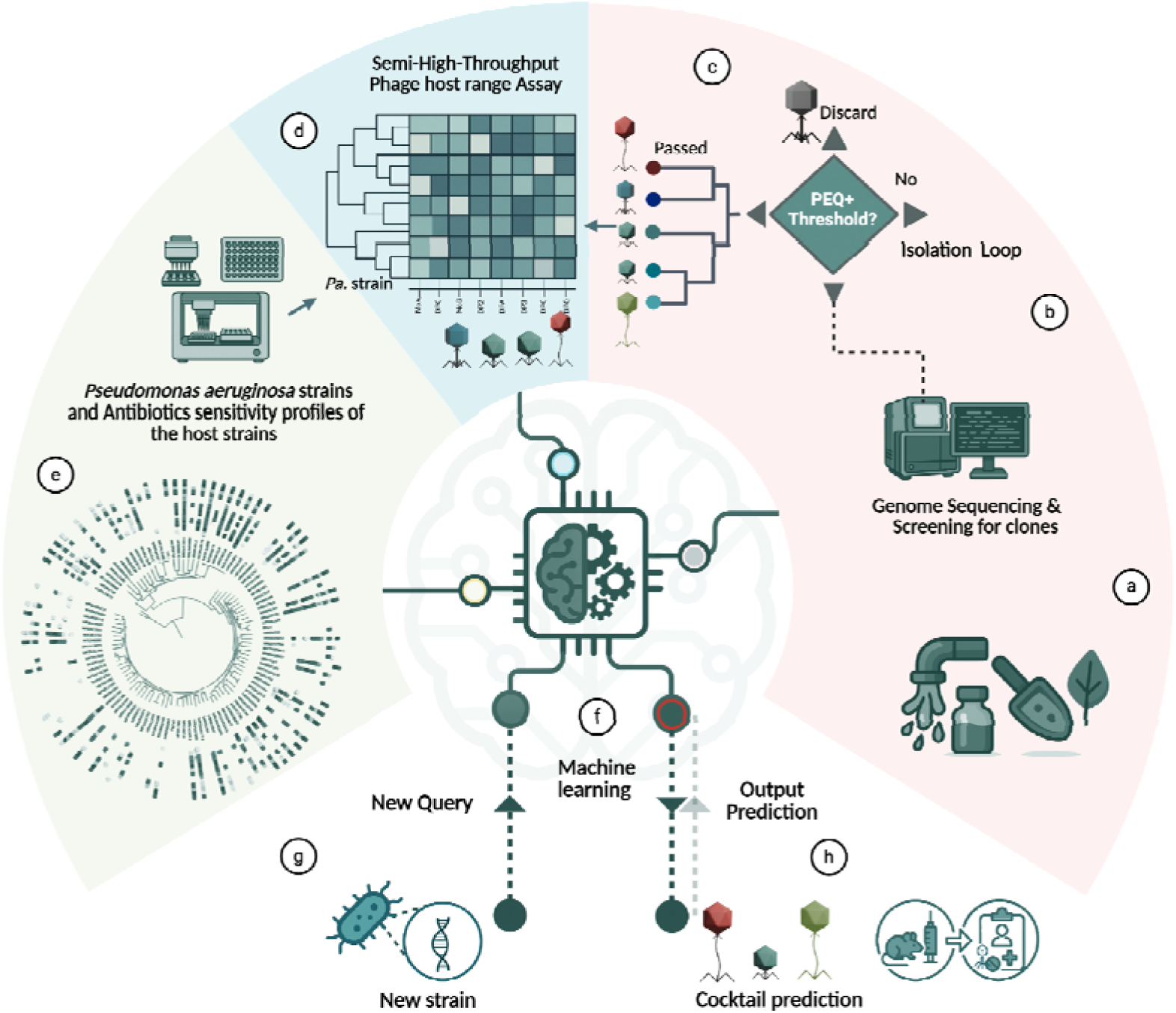
Experiment schematics. (a) Phages used in this study were either isolated in the lab or obtained from collaborators. Phages were enriched from diverse environmental samples on strain(s) of choice. (b) Phage genomes were sequenced using Illumina platform and the assembled contigs were annotated. (c) Phage genomes were compared using PhaMMseqs and PhamClust and were retained for the assay if the proteomic equivalent quotient (PEQ) met a determined threshold. Isolation and selection loop were repeated until we had the desired number of genetically diverse phages in our collection. (d,e) During each iteration, a new host strain was used for phage enrichment and isolation to increase the genomi diversity of isolated phages. Phages selected for the assay were aliquotted and spotted onto different hosts using a semi high-throughput phage host-range assay. The phenotype was scored manually as 0 (no positive interaction), 1 (hazy clearing of the lawn), or 2 (complete clearing of the lawn or individual plaques). (f) The data was used to train a machine learning model. (g,h) For an incoming new host genome, phage cocktails were formulated based on machine learning predictions, and their efficacy was tested in a murine model.

**Fig. 2.**
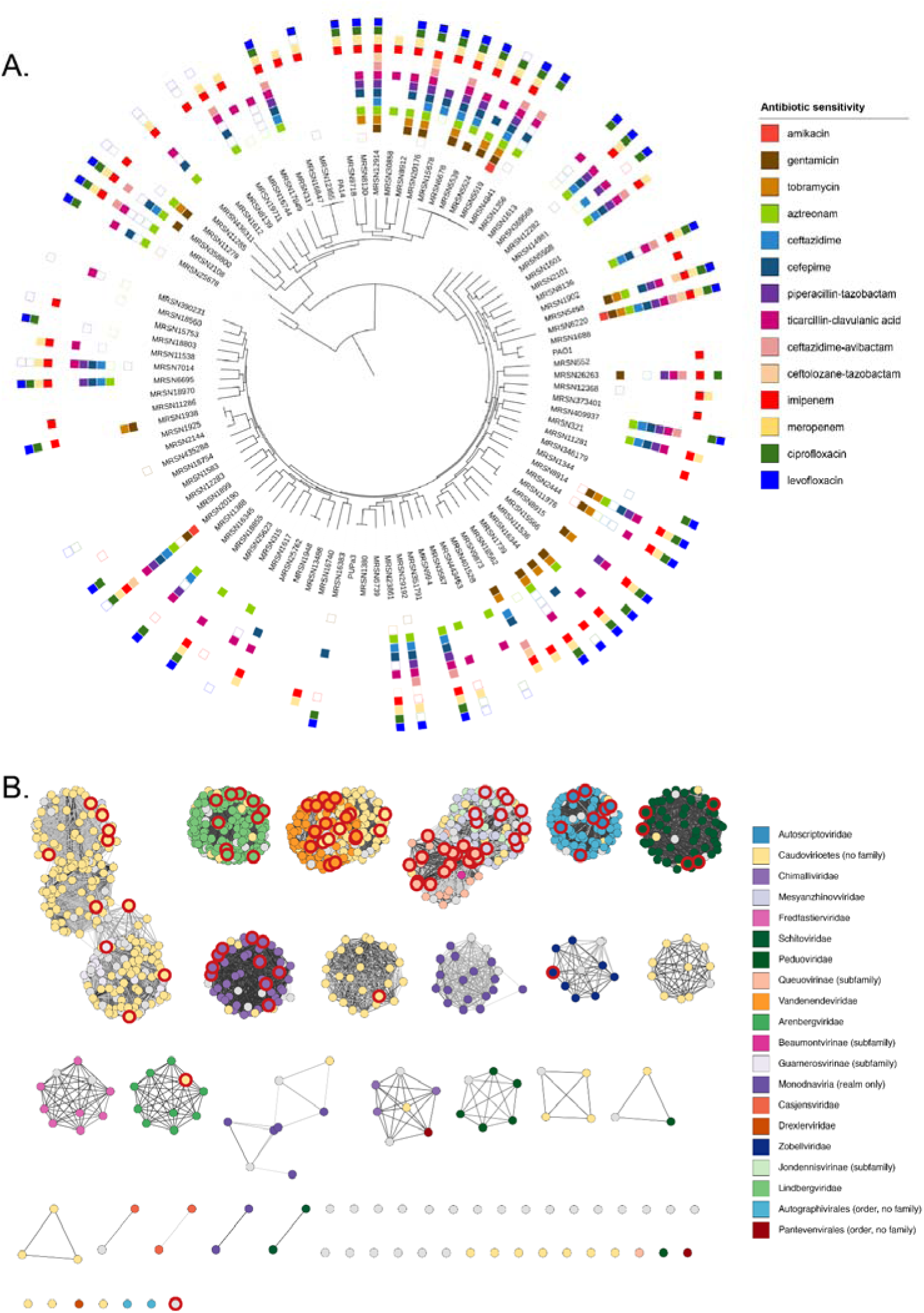
Description of *P. aeruginosa* strains and phages used in the assay. A) Neighbor-joining (rapidNJ) phylogenetic tree of 99 *P. aeruginosa* strains inferred from multiple alignment of the persistent pangenome portrays the genomic diversity of this collection. The phylogenetic tree was visualized in iTOL ^43^. Antibiotics sensitivity profiles of the strains, obtained from earlier publication ^15^ are indicated as following: filled squares (resistant), empty squares (intermediate) and sensitive (no squares). Antibiotics sensitivity profiles for PAO1, PA14 and PUPa3 have not been marked. B) Phage proteome similarity network of 95 phages used in this study and 1135 *P. aeruginosa* phage genomes available in the NCBI database (as of October 2025) visualized in Cytoscape. Phages from this study are marked with red node borders. Edges represent PEQ values above 0.15. Node colors and shapes represent different taxonomic groups as shown in the legend

**Fig 3.**
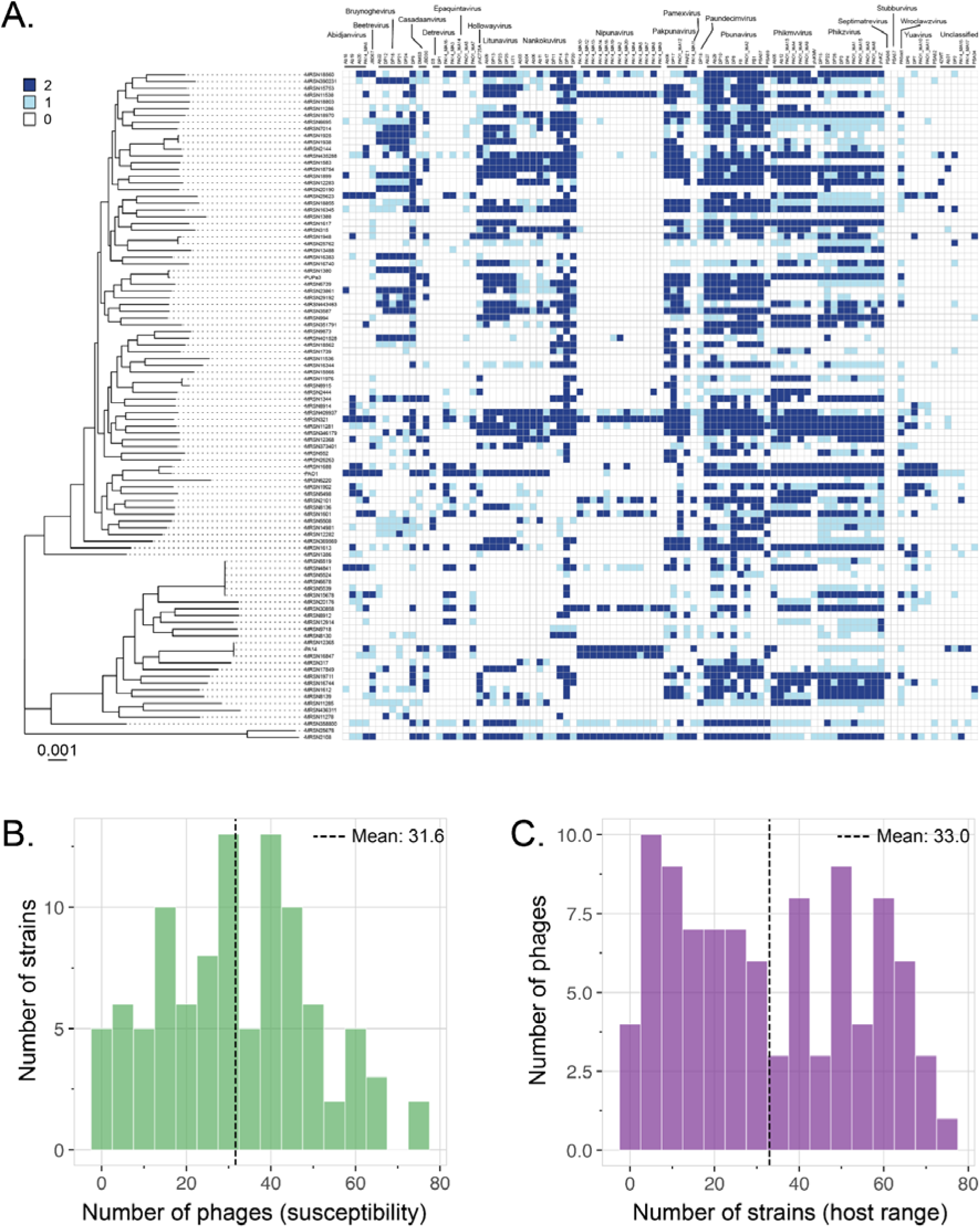
Susceptibility of a panel of genetically diverse *P. aeruginosa* phages on genetically diverse *P. aeruginosa* strains. (A) A manually curated panel of genetically diverse 95 *P. aeruginosa* phages were spotted on the 100-strain diversity panel of *P. aeruginosa* strains obtained from the Multidrug-Resistant Organism Repository and Surveillance Network (MRSN), Walter Reed Army Institute of Research. Commonly used model lab strains PAO1, PA14 and PUPa3 were also tested along with th MRSN panel. The infection outcome was manually scored (0, 1 or 2) after overnight incubation at 37 LJ. The score of “0” indicates that a positive interaction was not observed (white). The score of “1” indicate that a hazy clearing of the lawn was observed (light blue), whereas the score of “2” indicates that a complete clearing of lawn or individual plaques were observed (dark blue). The heatmap shows 9405 interaction datapoints between 95 phages and 99 strains; The color of the boxes indicate the interaction scores as described in the legend on the left side of the figure. The genomic diversity of the strains are represented by the maximum-likelihood phylogenetic tree (left of the figure) inferred from multiple alignment of the persistent pangenome. Phages used in the assay are listed on the top of the panel and are grouped based on genera; The names of phages belonging to the same genus are underlined and the genus name is indicated on the top. (B) The histogram shows the number of strains susceptible to a given number of phages; the mean susceptibility is 31.6 phages per strain, with a broad distribution indicating substantial variation from highly resistant to broadly susceptible strains. (C) The histogram shows the number of phages capable of infecting a given number of bacterial strains; the mean host range is 33.0 strains per phage, with a peak at 5-15 strains suggesting many phages are relatively specialized, while a secondary elevation at 45-60 strains reflects a subset of broadly infective phages.

For the phage collection, we assembled 95 genomically diverse *P. aeruginosa* phages through multiple isolation strategies (Methods). To ensure maximal diversity, we applied iterative curation using PhamClust, which computes proteomic equivalence quotient (PEQ) scores representing the product of shared gene fraction and average amino acid identity between phage pairs^41^. Phages were selected for inclusion only if their PEQ scores against all existing collection members fell below 0.95, corresponding approximately to the ICTV species boundary (>95% nucleotide identity)^42^ (*Supp. Fig. 4*) The resulting collection spans 20 recognized genera (*Fig. 2B*), representing one of the most genomically diverse *P. aeruginosa* phage panels assembled for systematic interaction profiling to date. Genus *Nipunavirus* is most abundantly represented (13 phages), while seven genera (*Beetrevirus*, *Hollowayvirus*, *Pamexvirus*, *Paundecimvirus*, *Septimatrevirus*, *Stubburvirus*, *Wroclawvirus*) contribute single representatives, and six phages remain taxonomically unclassified.

**Fig. 4.**
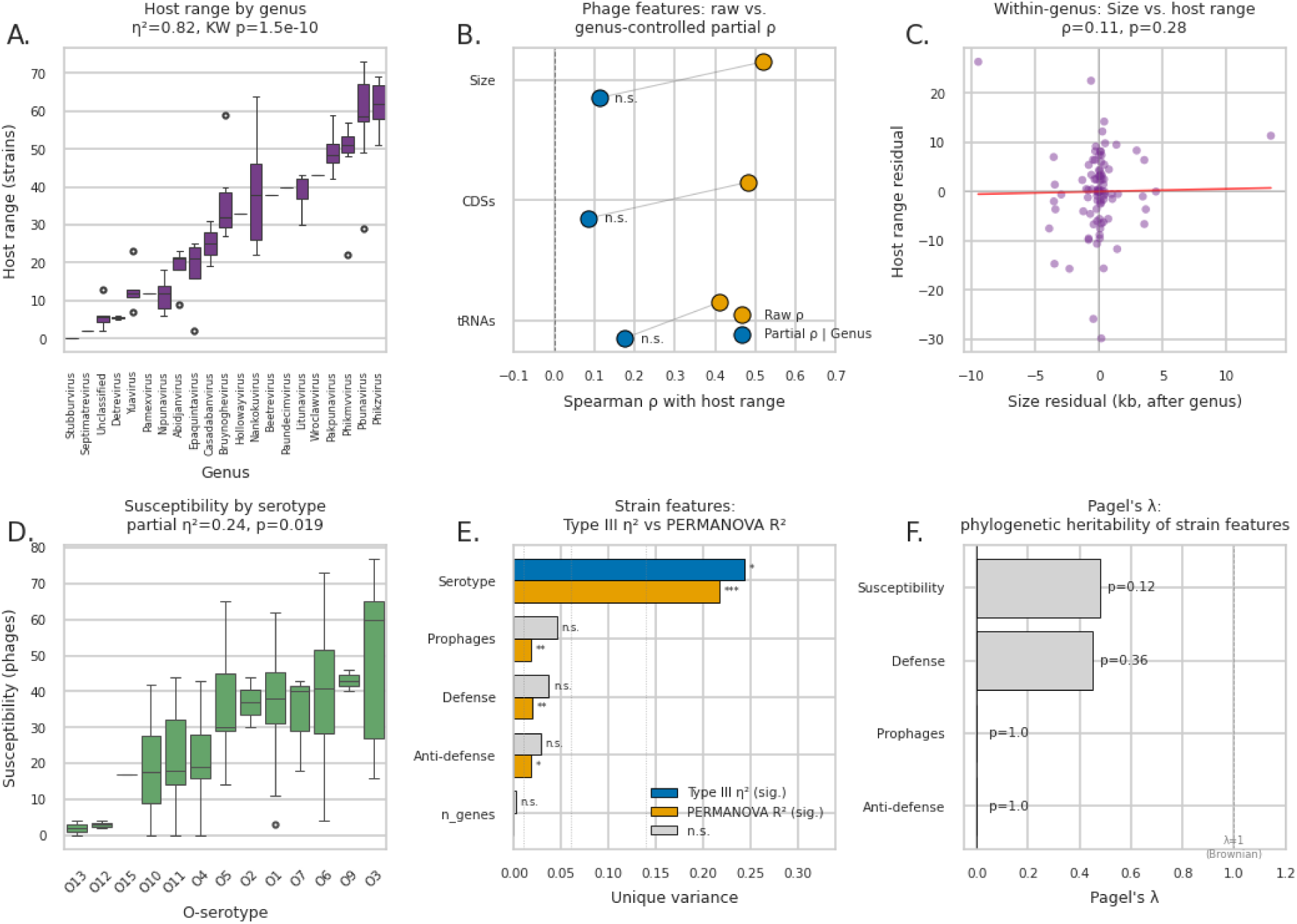
Multivariate-aware analysis of phage- and strain-level genomic features. Top row: phage host range (n = 95 phages). A. Host range by phage genus (14 genera with ≥ 2 representatives shown). Host range differed significantly among genera (Kruskal–Wallis H = 73.92, η² = 0.823, ε² = 0.850, p = 1.5×10⁻¹⁰). B. Forest plot of raw (orange) vs. genus-controlled partial (blue) Spearman ρ for genome size, CDS count, and tRNA gene count vs. host range. All three univariate correlations (ρ = 0.41–0.52, p < 10⁻⁴) become non-significant after controlling for genus (partial ρ = 0.09–0.17, p > 0.08); n.s. = not significant. C. Within-genus residual scatter of genome size vs. host range, after removing genus-mean effects on both axes (Spearman ρ = 0.11, p = 0.28; red line, OLS fit). The genus-controlled signal is essentially flat. Bottom row: strain susceptibility (n = 99 strains for univariate boxplot in D and Type III ANOVA in E; n = 95 infectable strains for PERMANOVA in E and Pagel’s λ in F). D. Susceptibility by O-antigen serotype across 13 types (Kruskal–Wallis H = 31.60, η² = 0.228, ε² = 0.322, p = 1.6×10⁻³). E. Grouped bar chart comparing partial η² from Type III ANOVA on per-strain susceptibility count (left bars) and partial R² from phylogeny-controlled PERMANOVA on Jaccard infection-pattern distances (right bars), for each strain-level predictor. Bars are colored when significant (blue = Type III, orange = PERMANOVA) and gray when not significant; significance markers indicate * p < 0.05, ** p < 0.01, *** p < 0.01. Cohen’s effect-size reference lines for small/medium/large shown as dotted verticals. n_genes was tested in the Type III ANOVA only. Both analyses agree that serotype is the dominant predictor; defense, anti-defense, and prophage burden show the magnitude/pattern asymmetry described in the text (significant for infection pattern but borderline or not significant for susceptibility magnitude). F. Pagel’s λ for strain-level features on the marker-gene phylogeny. Anti-defense count and prophage burden show no phylogenetic signal (λ ≈ 0, LR p = 1.0); defense count and overall susceptibility show moderate phylogenetic signal (λ = 0.45 and 0.48, respectively) that does not reach LR-test significance (p = 0.36 and 0.12). Bars are colored when LR p < 0.05 and gray otherwise; vertical reference at λ = 1 indicates Brownian-motion expectation.

### Comprehensive phage-host interaction profiling reveals genus-level host range patterns with critical strain-level variation

We conducted systematic spot assays across all 95 phages and 99 strains, generating 9,405 pairwise interaction datapoints, one of the largest such dataset reported for *P. aeruginosa* (Methods). Clearance-based spotting assays have been routinely used as a proxy for phage host-range to capture a range of bactericidal or bacteriostatic interactions, including mechanical or enzymatic lysis and growth inhibition, in addition to full productive lysis^44^. However, as these phenotypes are consistent, strain-specific, and relevant in the clinical targeting of *P. aeruginosa*, specifically in diffusion-limited environments like biofilms, we posit them to represent a suitable metric for phage-host interaction characterization. Interactions were scored as 0 (no visible lysis), 1 (partial or turbid lysis), or 2 (clear lysis). Of the 9,405 pairings, 3,132 (33.3%) produced positive outcomes (score ≥1): 2,015 exhibited complete lysis (score 2; 21.4% of all pairs) and 1,117 showed partial lysis (score 1; 11.9% of all pairs). The remaining 6,273 pairs (66.7%) showed no interaction (*Fig. 3A, Supp. Fig. 5, Dataset S2*). This positive interaction rate is comparable to other large-scale *P. aeruginosa* datasets and higher than rates observed in other phage-host systems^36,45,46^.

**Fig. 5.**
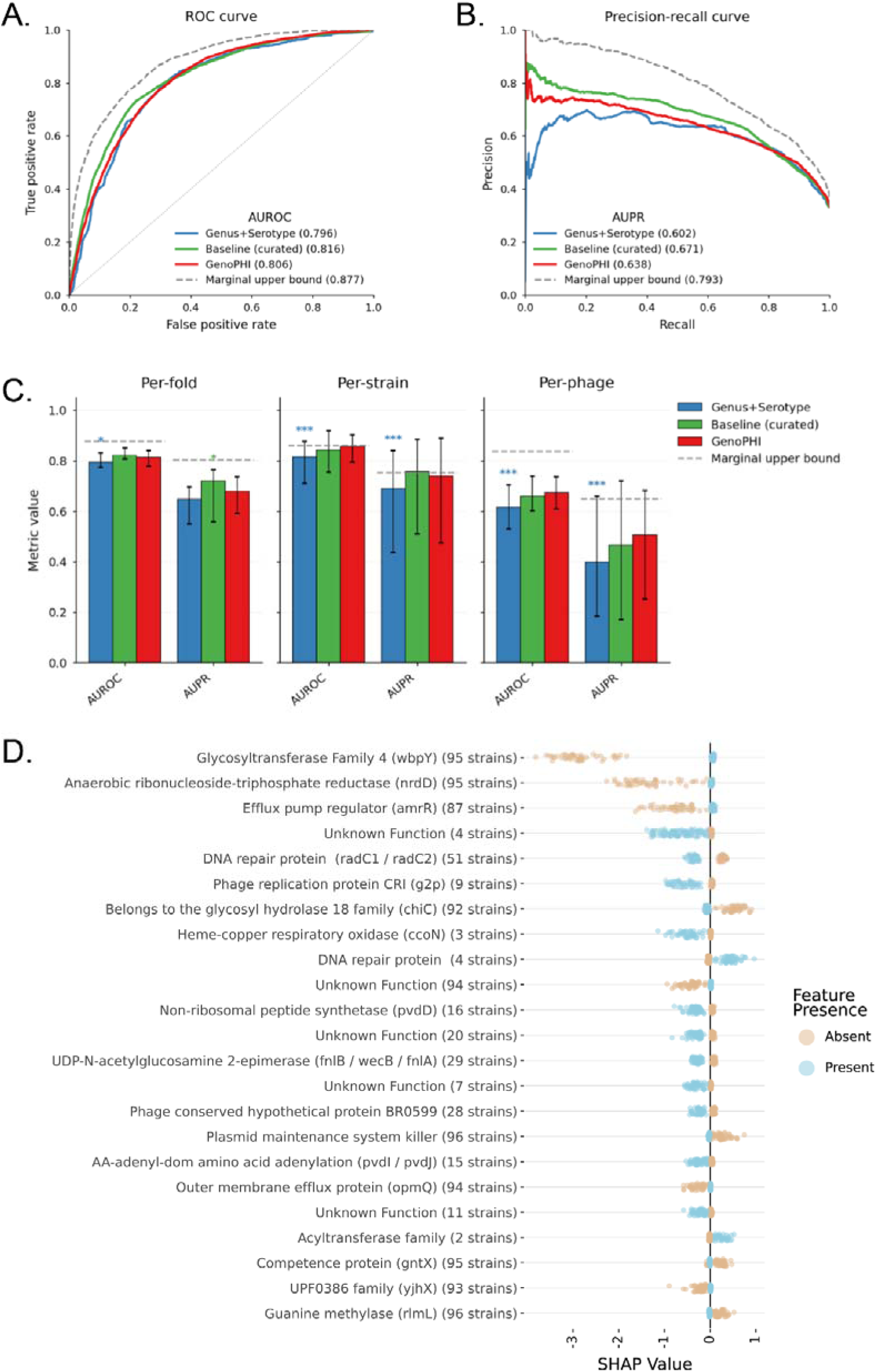
Model performance and feature importance. (**A**) ROC and (B) precision–recall curves for all three models and the marginal upper bound (gray dashed line), computed on predictions pooled acros all folds. Legend entries show AUROC and AUPR. (C) Cross-validated AUROC and AUPR for three CatBoost models (1) Genus+Serotype, (2) Baseline (curated genomic features), and (3) GenoPHI (pangenome-derived gene-cluster features) under three aggregation views: per-fold, per-strain, and per-phage. For per-fold, predictions are pooled within each of 20 strain-held-out folds. For per-strain and per-phage, predictions are pooled across all folds in which the entity appeared and one AUROC/AUPR is computed per entity. Bars show the median across folds or entities; error bars span the interquartile range. Asterisks indicate paired Wilcoxon significance against GenoPHI within each metric and view (*p < 0.05, **p < 0.01, ***p < 0.001). Gray dashed lines show the non-deployable degree-only reference (closed-form prediction from per-phage breadth × per-strain susceptibility) at the matching aggregation view. (D) SHAP values associated with predictive features indicate whether presence of a feature (blue) or absence (orange) is associated with increased likelihood of infection (SHAP value > 0) or decreased likelihood of infection (SHAP value < 0).

Individual phage host ranges varied dramatically. Ab29 (*Pbunavirus*) infected 73 strains (73.7%), DP8 (*Pbunavirus*) infected 71, and DP4 (*Phikzvirus*) infected 69, representing the broadest-range phages in the collection. At the opposite extreme, PSA57 (*Stubburvirus*) showed no positive interactions with any strain in the collection (though it was successfully propagated on an isolate not included in the panel), while PAO1_MA14 (*Epaquintavirus*), PSA56 (*Septimatrevirus*), and PA14_MA17 (unclassified) each infected only two strains (*Dataset S2*). The distribution of phage host ranges was bimodal, with peaks at 5–15 strains and 45–60 strains, consistent with distinct populations of specialist and generalist phages (*Fig. 3C*). Cross-genus pairwise Jaccard analysis of the broadest-range phage from each of 10 well-sampled genera revealed convergence on shared hosts ∼40% above marginal-rate expectation (mean Jaccard 0.36 vs. 0.25 expected under independent host sampling; convergence ratio 1.40; *Supp. Fig. 6*). Hierarchical clustering identified modest within-pool sub-structure but no qualitatively distinct broad-spectrum strategies (*Supp. Fig. 7*). This cross-genus convergence is comparable to within-genus convergence among the top-10 broadest phages overall (ratio 1.38), indicating that apparent generalism is not primarily driven by within-genus shared receptor ancestry.

At the genus level, three genera demonstrated broad host range, infecting more than half of the strain panel: *Phikzvirus* (60 ± 6 positive interactions), *Pbunavirus* (59 ± 13), and *Pakpunavirus* (50 ± 7) (*Table S1*). Phage host range varied significantly by genus (Kruskal-Wallis H = 73.92, η² = 0.823, ε² = 0.850, p = 1.5×10□¹□) (*Fig. 4A*). However, substantial within-genus variation was observed. In *Pbunavirus*, Ab29 infected 73 strains while PSA69 infected only 29. In *Phikzvirus*, host range spanned 51 strains (PAO1_MA8) to 69 (DP4). In *Pakpunavirus*, coverage ranged from 42 (PAPZ1) to 59 (DP17). Critically, the broadest individual phage (Ab29) exceeded the mean of even the broadest genus (*Phikzvirus*), demonstrating that genus classification alone cannot predict individual phage performance and that strain-level characterization remains essential for therapeutic selection.

Bacterial strain susceptibility was equally variable, ranging from zero interactions (four pan-resistant strains) to 77 positive interactions (MRSN2108), with a mean of 32 ± 18 interactions per strain (*Fig. 3B*). Network topology analysis revealed two structural features of the interaction structure. First, the network was significantly anti-nested: NODF = 60.5 was lower than the null distribution from 1000 swap-null replicates that preserved marginal degree distributions (Z = −4.0, one-sided p < 0.001 for anti-nestedness; *Supp. Fig. 8*), indicating that broad-range phages and narrow-range phages target non-overlapping host subsets rather than a hierarchical generalist-includes-specialist architecture. This anti-nestedness extends to within-lineage structure: per-genus NODF testing identified five of 10 well-sampled genera as individually anti-nested after Benjamini-Hochberg multiple-testing correction (*Pbunavirus*, *Phikmvvirus*, *Bruynoghevirus*, *Epaquintavirus*, *Litunavirus*; z = −2.87 to −12.57; *Table S2*). Second, the network showed no significant modularity (Q = 0.024, p = 0.14; *Supp. Fig. 9*). Together with the anti-nestedness result, this is consistent with niche partitioning between distinct phage-host sub-communities at multiple taxonomic scales, possibly reflecting receptor-system specialization and defense-system heterogeneity.

### Phage genus and bacterial O-antigen serotype are the dominant determinants of interaction breadth

To determine whether the observed patterns of phage-host interactions could be explained by defined genomic and phenotypic features, we examined a set of phage- and strain-level variables. On the phage side, we assessed taxonomic classification at the genus level, genome size, number of coding sequences (CDSs), and tRNA gene content (*Supp. Fig. 10*). On the strain side, we evaluated O-antigen serotype, prophage burden, and the number of annotated phage defense and anti-defense systems. These features were selected to capture the dominant biological axes implicated in *P. aeruginosa* phage compatibility by the published literature: structural and genus-level determinants of phage host range (genus, genome size, CDS count, tRNA content), and the host-side immune and surface determinants of susceptibility (O-antigen serotype, defense and anti-defense system counts, prophage burden). We treat this set as a curated, mechanistically interpretable baseline against which the pangenome-based GenoPHI representation^39^ is later compared, rather than as an exhaustive feature space.

Several of these features are biologically expected to covary. On the phage side, genome size and CDS count are nearly redundant (Spearman ρ = 0.93), and both correlate moderately with tRNA gene count (ρ = 0.56 and 0.66, respectively). On the strain side, prophage burden correlates with anti-defense count (ρ = 0.48) and total gene count (ρ = 0.84) (*Supp. Fig. 11-12*). Such collinearity can produce misleading univariate associations when correlated features are interpreted as independent predictors^47^, and motivates the multivariate analyses that follow. Variance inflation factors for the predictor matrices used in the analyses below were below 5 (Type III ANOVA max VIF = 3.5; PERMANOVA max GVIF = 1.79; *Supp. Tables S3-S4*), confirming that multicollinearity is moderate and that unique-variance contributions from each predictor can be interpreted at face value.

On the phage side, host range was overwhelmingly explained by phage genus, which accounted for ∼82–85% of the observed variance (Kruskal–Wallis H = 73.92, η² = 0.823, ε² = 0.850, p = 1.5×10□¹□). Post-hoc pairwise comparisons using Dunn’s test with Bonferroni correction revealed multiple statistically significant differences among genera, with the most pronounced contrasts between narrow-range genera (e.g., *Nipunavirus*, median host range = 12 strains) and broad-range genera including *Pbunavirus* (median 58.5) and *Phikzvirus* (median 62; both p < 0.001). Genome size, coding sequence (CDS) count, and tRNA gene count showed strong univariate Spearman correlations with host range (ρ = 0.41–0.52, all p < 10□□), but all three became non-significant after controlling for genus via partial correlation (Size partial ρ = 0.11, p = 0.28; CDSs partial ρ = 0.09, p = 0.41; tRNAs partial ρ = 0.17, p = 0.09; *Table S5*). Per-genus Spearman tests within each of the 10 well-sampled genera further showed no consistent within-genus correlation between genome content and host range (24 valid tests, none surviving Benjamini–Hochberg correction; *Table S6*). Together these analyses indicate that genome content correlates with host range only through its covariation with genus, rather than as an independent predictor at the per-phage level. In a phylogeny-controlled distance-based redundancy analysis (dbRDA)^48^ on Jaccard distances of phage infection profiles, phage genus uniquely explained 27.7% of variance after partialling out the first 10 principal coordinates of neighbor matrices (PCNM eigenvectors) derived from the PEQ phage tree (F = 2.93, p < 0.001; *Table S7*). The remaining variance attributable to genus in the unconditioned model (63.9% total; difference = 36.2 percentage points) was therefore shared with PEQ-based phylogenetic structure, but a substantial unique component remained. This residual genus contribution confirms that the genus effect is not solely a proxy for fine-grained phage genomic similarity but reflects genus-level biology, potentially including receptor utilization, structural protein homologs, or anti-defense system content that vary between but not within genera. We avoid implicating anti-defense content here, as our phylogenetic-signal analysis below finds essentially no heritable structure for anti-defense count (Pagel’s λ ≈ 0).

To disentangle the contributions of correlated strain-level features, we conducted two complementary analyses: a Type III ANOVA on per-strain susceptibility count (the magnitude of susceptibility / the total number of phages infecting each strain), and a phylogeny-controlled PERMANOVA on Jaccard distances of per-strain infection profiles (the pattern of susceptibility / which specific phages infect each strain). Different features may predict patterns without predicting magnitude when they shape specificity of compatibility rather than overall susceptibility. Both identified serotype as the dominant unique predictor (partial η² = 0.243, F = 2.20, p = 0.019; PERMANOVA R² = 0.218, F = 2.24, p < 0.001; *Table S8-S9*), consistent with O-antigen and lipopolysaccharide structure being a primary determinant of receptor accessibility and compatibility. The full count-based model explained 47% of variance in strain susceptibility (adjusted R² = 0.365, F(16,82) = 4.52, p = 2.5×10□□; *Table S10*); the pattern-based model explained ∼44% of pairwise infection-profile distance variance, with phylogenetic eigenvectors contributing only ∼5% in aggregate (PCNM5 the only individually significant axis at p = 0.025).

The two analyses diverge informatively on the secondary predictors. Defense system count, anti-defense count, and prophage burden each contributed small but significant unique variance to susceptibility pattern (PERMANOVA R² = 0.018–0.019, all p ≤ 0.011; *Table S11*) but contributed marginally or not at all to susceptibility magnitude (Type III partial η² = 0.028–0.045, p = 0.052–0.13). This magnitude/pattern asymmetry indicates that defense, anti-defense, and prophage burden shape which specific phages successfully infect a strain rather than uniformly reducing the total number of compatible phages, consistent with these features being phage-type-specific in their effect (e.g., CRISPR vs. RM systems targeting specific phage sequences) rather than acting as a general susceptibility damper.

To assess whether these strain-level features are inherited along strain phylogeny, we computed Pagel’s λ for each feature on the marker-gene tree (n = 95 strains; *Table S12*). Anti-defense system count and prophage burden showed essentially no phylogenetic signal (λ ≈ 0; LR test p = 1.0 for both), consistent with horizontal mobility of prophages and prophage-encoded anti-defense modules. Defense system count (λ = 0.45) and overall susceptibility (λ = 0.48) showed moderate phylogenetic structure but did not reach formal significance under the LR test (p = 0.36 and 0.12, respectively), indicating partial clonal structure that is not strong enough to dominate compatibility prediction at this sample size. Together with the small unique phylogenetic contribution in the multivariate model (combined PCNM R² = 0.052), these analyses indicate that phage compatibility is determined by features that are at most partially heritable along strain phylogeny, supporting per-strain genomic characterization rather than reliance on phylogenetic placement to predict susceptibility.

Taken together, these analyses separate the magnitude of susceptibility from the pattern of which phage-strain pairs interact. Phage genus and host O-antigen serotype shape both magnitude and pattern; defense, anti-defense, and prophage content shape pattern without altering magnitude, modifying which specific phages succeed rather than how many. Combined with the global and within-genus anti-nestedness of the interaction matrix, this means that phages with similar host-range sizes can target non-overlapping host sets. For predictive modeling, representations that capture only the magnitude axes will rank phages well on average but fail on the strain-specific compatibility structure that determines which phage in a cocktail covers which gap.

### Predictive modeling of phage-host interactions

To translate the interaction dataset into predictive capability, we trained a series of nested CatBoost models under a 20-fold strain-based nested cross-validation scheme that withheld 10% of strains entirely from feature selection, training, and tuning per fold, simulating encounter with a novel patient isolate. We compared three deployable models of increasing feature richness against a non-deployable degree-only reference. The deployable models were Genus+Serotype (phage genus and host O-antigen only), a curated baseline adding defense and anti-defense system counts, prophage burden, genome size, and gene count, and GenoPHI, which replaces curated features with pangenome-derived MMSeqs2 protein clusters and model-based feature selection. The degree-only reference assigns each pair the product of full-matrix phage breadth and host susceptibility. This model is not deployable for novel isolates because it requires knowledge of the test strain’s interactions. It serves to quantify the predictive performance achievable from first-order degree structure alone, establishing what can be achieved from connectivity patterns as a benchmark for model performance.

Pooled across all held-out predictions (Fig. 5A-B), the three deployable models reached AUROC 0.80–0.82 and AUPR 0.60–0.67, with the curated baseline modestly ahead (pooled AUROC 0.82, AUPR 0.67) of GenoPHI (0.81, 0.64) and Genus+Serotype (0.80, 0.60); all three trailed the marginal reference (gray dashed; AUROC 0.88, AUPR 0.79). Splitting performance by held-out entity revealed an asymmetry hidden in the pooled view (*Fig. 5C, Dataset S3, Supp. Fig. 13*): on per-strain metrics, deployable models matched the degree-only reference, recovering the per-strain ranking obtainable from degree structure alone (GenoPHI per-strain median AUROC 0.86 vs reference 0.86), while on per-phage metrics they fell well short (GenoPHI per-phage median AUROC 0.67 vs reference 0.84). This asymmetry is consistent with the receptor-dominant architecture identified in the previous section: per-strain susceptibility integrates a small set of strong, readable signals (most prominently O-antigen serotype, recoverable from LPS biosynthesis genes), whereas per-phage host range is set largely by genus-level structural biology that is not fully reducible to the curated features available to any single model. Among deployable models, GenoPHI significantly outperformed Genus+Serotype on all per-entity metrics (*Dataset S3*; all p ≤ 3×10⁻⁴, paired Wilcoxon) and was statistically indistinguishable from the curated baseline at the median (all p ≥ 0.24).

Aggregate medians, however, obscure where model differences are concentrated. We binned per-entity performance along four partially non-redundant difficulty axes including phylogenetic novelty, phenotypic divergence from nearest neighbors, marginal difficulty (failures of degree-based prediction), and full-matrix degree (*Supp. Fig. 14, Dataset S4*). The trade-off was consistent across axes. On easy phage terciles the curated baseline outperformed GenoPHI on both AUROC and AUPR (all p ≤ 0.003), while on hard phage terciles GenoPHI outperformed the curated baseline (all p ≤ 0.021), with both feature-rich models substantially outperforming Genus+Serotype across all difficult bins. The strain-side pattern was directionally similar but smaller, reaching significance only at high marginal difficulty (per-strain AUROC 0.78 vs 0.73, p = 0.005) (*Supp. Fig. 15*). The pattern reflects different inductive biases. Curated features encode a small set of well-understood biological signals that suffice for routine cases, while pangenome features raise the floor on hard cases at modest cost to easy ones.

The difficulty-dependent trade-off translates into clinically meaningful differences for hard-to-treat strains. We used each model’s confidence to select 3- or 5-phage cocktails for held-out strains under HDBSCAN-derived receptor-diversity constraints, and compared each model against two non-ML baselines: a greedy set-cover that maximizes strain coverage across training-fold data, and a published promiscuity-based method that selects the most broadly infectious phage per cluster from training-fold interaction data; all approaches are deployable from a training-fold interaction matrix (*Table S14 & S15*). Cocktail success was defined as ≥1 phage in the top-k recommendation that infects the strain. On the lowest-susceptibility quartile (Q1), where individual phage selection matters most, GenoPHI achieved the highest top-3 success rate at 65.8%, significantly exceeding Genus+Serotype (31.6%, McNemar p = 0.002) and trending above the curated baseline (47.4%, p = 0.092), greedy set-cover (47.8%, p = 0.45), and the promiscuity baseline (57.9%, p = 0.61). At top-5 in Q1, GenoPHI tied the promiscuity baseline at 78.9%, both significantly exceeding the curated baseline (55.3%, p = 0.022) and Genus+Serotype (44.7%, p = 0.002). By Q3–Q4 all approaches converged at ≥98% top-3 success, reducing cocktail design to a covering problem. Because the susceptibility profile of a new clinical isolate is unknown a priori, we selected GenoPHI for downstream in vivo validation.

SHAP analysis (*Fig. 5D, Supp. Fig. 17*) showed that bacterial surface architecture dominates predictions. Among the top 25 features, LPS and polysaccharide biosynthesis genes, including *wbpY* (the single most important feature), *chiC*, *fnlA*, *fnlB* and *wecB* are consistent with serotype-level signal at gene resolution. Outer membrane protein OmpQ, efflux regulator AmrR, and type IV pili and fimbriae-associated proteins, all candidate or established phage receptors in *P. aeruginosa*, were absent from the curated representation. Prophage-related proteins (g2p, gntX) overlap mechanistically with the curated prophage-burden feature. GenoPHI’s top features thus overlap mechanistically with the curated representation (LPS biosynthesis to serotype; phage/prophage proteins to prophage burden) while additionally surfacing candidate receptor mechanisms absent from the curated model.

### Comparison of classical genetics and machine learning approaches for phage cocktail design

To compare ML-guided and expert-driven cocktail design head-to-head, we constructed two three-phage cocktails targeting PAO1 under distinct selection strategies for resistance management: a classical-genetics cocktail (CG) maximizing genus and receptor diversity under curated biological constraints, and a GenoPHI-selected cocktail (ML) chosen by the deployable pangenome-based model. Critically, the two cocktails differ in what was known about PAO1 when they were designed. The CG cocktail was built using direct lab testing of phages on PAO1 and on PAO1 mutants, while the ML cocktail was selected with PAO1 completely excluded from model training (Fig. 1). The comparison therefore contrasts a best-case expert cocktail informed by the target strain against a blind predictive cocktail.

For the classical genetics (CG) cocktail, we first identified genera with broad host range from the interaction matrix (*Phikzvirus*, *Pbunavirus*, *Pakpunavirus*, *Phikmvvirus*). Receptor diversity was then imposed as a selection criterion to minimize the probability that single bacterial mutations could confer cross-resistance to all cocktail components, a key design principle for therapeutic phage cocktails^49,50^. Phage phiKZ (*Phikzvirus*, flagellum receptor) and phage phiKMV (*Phikmvvirus*, type IV pili receptor) were selected based on their established, characterized receptor usage^51–53^. To identify a third phage utilizing a distinct receptor, we generated PAO1 mutants independently resistant to phiKZ (PAO1^phiKZR) and phiKMV (PAO1^phiKMVR), then screened the phage panel for candidates producing clear lysis zones on lawns of both resistant mutants. Phage PAO1_MA12 (Pakpunavirus, the members of which use the O-specific antigen (OSA) as the receptor^54,55^) met this criterion, yielding a CG cocktail of phiKZ + phiKMV + PAO1_MA12 representing three genera and three confirmed receptor systems (*Table S17*).

The ML cocktail was designed by training models on all strains except PAO1 (which was held out from feature selection and training) (*Supp. Fig. 18A-D*), then applying the GenoPHI cocktail design algorithm, which jointly optimizes predicted infection probability and mechanistic diversity, to select the three optimal phages from the collection (*Supp. Fig. 18E-F*). The model thus produced cocktail predictions for PAO1 without having seen any PAO1 interaction data, simulating the clinical scenario of a newly sequenced patient isolate not represented in the training data. The algorithm selected DP4 (*Phikzvirus*), Ab03 (*Nankokuvirus*), and PAO1_MA13 (*Phikmvvirus*), all of which scored “2” (clear lysis) against PAO1 in the interaction matrix (*Table S17*). Both cocktails included representatives of *Phikzvirus* and *Phikmvvirus*, with the ML pipeline selecting phages predicted to have broader host range (DP4 and PAO1_MA13) over the empirically characterized members in each genus (phiKZ and phiKMV); the corresponding ML and CG pairs are genomically near-identical (DP4–phiKZ: 98.6% nucleotide identity over 97% query coverage; PAO1_MA13–phiKMV: 93.2% identity over 91% query coverage; *Table S17*) The cocktails diverged on the third selection: the CG cocktail included PAO1_MA12 (*Pakpunavirus*, selected to broaden receptor diversity via cross-resistance screening), while the ML cocktail included Ab03 (*Nankokuvirus*, selected by the cocktail-design algorithm and from a genus not characterized as broadly infective in our genus-level analysis and not among the receptor-characterized genera used to build the CG cocktail). The ML pipeline thus contributes in two distinct ways: within receptor-similar genera, it identifies the best-predicted member of the cluster (raising single-phage host range from 22 to 56 strains in the *Phikmvvirus* case; *Table S17*); and at the third-component selection, it diverges from the CG cocktail’s curated-receptor logic to select a phage the model itself predicts will perform well.

Both cocktails suppressed PAO1 growth in overnight liquid culture assays. While individual phages allowed bacterial regrowth (visible as OD increase) after initial suppression, neither the CG nor ML cocktail permitted detectable regrowth, consistent with multi-component cocktails presenting a higher barrier to resistance evolution (*Supp. Fig. 19*). These *in vitro* performance tests validated both cocktails for progression to *in vivo* evaluation.

### Machine learning-guided phage cocktails demonstrate therapeutic efficacy in a murine wound infection model

To provide in vivo proof-of-concept that ML-guided phage selection translates to measurable therapeutic benefit, we evaluated CG and ML cocktails in an established murine full-thickness wound infection model. BALB/c mice (10–16 weeks, male) received two 6 mm excisional wounds inoculated with PAO1 (2×10□ CFU per wound). Phage cocktails (10□ PFU/mL, 100 µL per wound) or SM buffer control were administered at 2 h, 24 h, and 48 h post-infection (*Fig. 6A*); wounds were collected and homogenized at 72 h for bacterial enumeration on selective Cetrimide agar.

**Fig 6.**
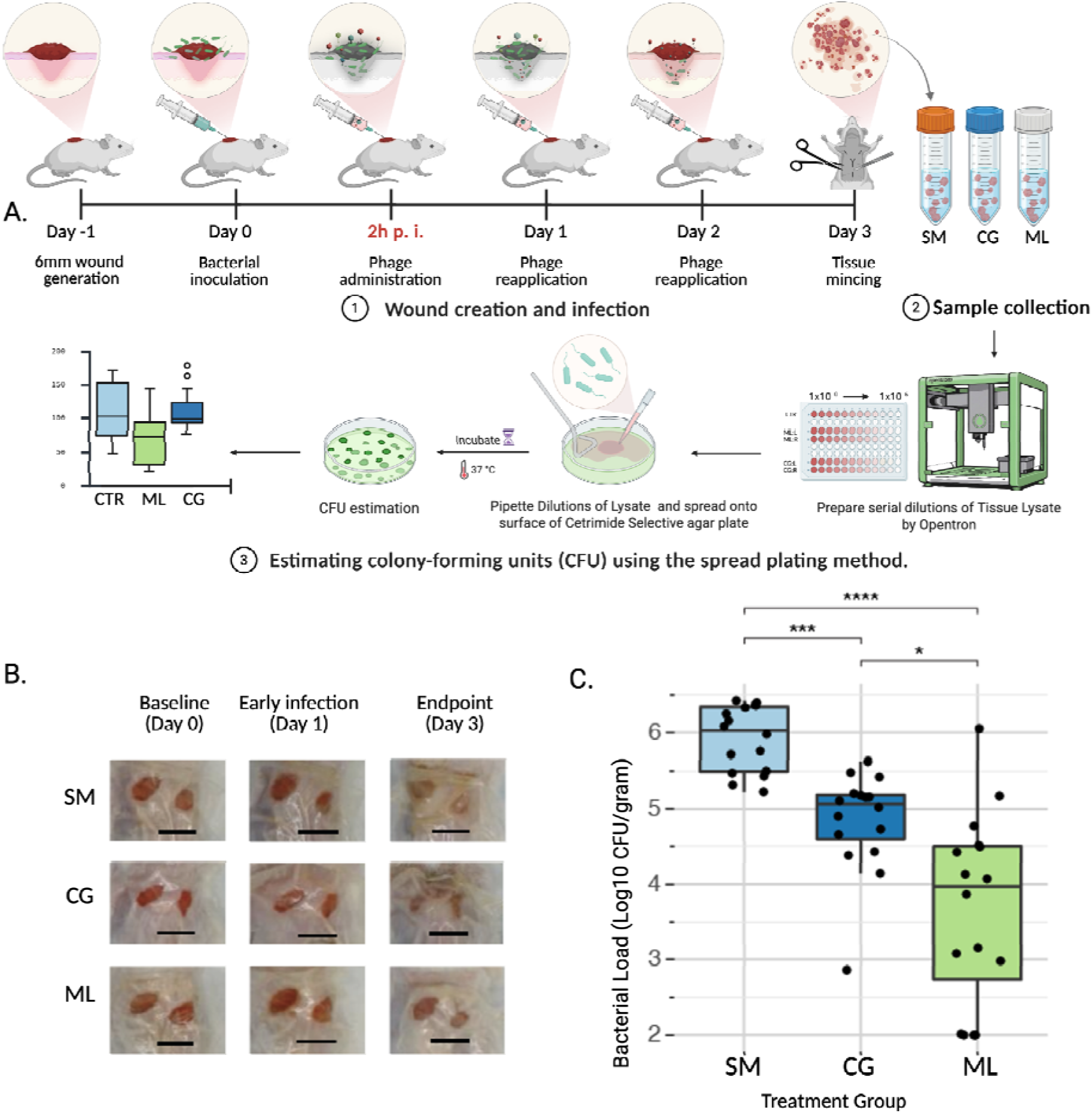
Phage cocktails reduce bacterial burden in male mouse PAO1 wound infections. (**A**) A full-thickness dorsal wound was created and inoculated with *P. aeruginosa* PAO1. Two hours post-infection, animals received the first topical treatment with either vehicle control (SM), a classical genetics–derived phage cocktail (CG), or a machine learning–predicted phage cocktail (ML). Additional treatments were administered at 24 h and 48 h post-infection. Wounds were monitored and collected on the collection day (72 h post-infection). (B) Representative wound images acquired at the indicated time points. A 10 mm scale bar is shown in each image. (C) Data are presented as logLJLJ CFU per wound (mean ± SEM). The limit of detection (LOD) was 100 CFU per wound (logLJLJ = 2). Each group had 16 wounds (8 animals). Groups were compared by Kruskal-Wallis test with Dunn’s multiple comparisons test and Holm correction. *P < 0.05, **P < 0.01, ***P < 0.001, ****P < 0.0001

The mice appeared clinically well throughout the study, with minimal weight loss and no obvious differences in overall behavior or gross appearance among groups. Wounds in the SM group appeared more frequently opaque than those in the CG and ML groups, consistent with a higher bacterial burden in SM-treated wounds (*Fig. 6B / Table S18*).

Treatment effects on bacterial burden were quantified by CFU enumeration on selective Cetrimide agar^56^. Values below the limit of detection (LOD = 100 CFU/gram) were set to the LOD for statistical analysis; this censoring affected 3/16 ML samples (no CG or SM samples), so the reported ML-vs-CG and ML-vs-SM fold reductions are conservative lower bounds on the true ML effect. By CFU enumeration, both the CG and ML cocktails produced substantial reductions in bacterial burden relative to the SM buffer control (Kruskal–Wallis P = 1.9×10□□; Dunn’s post-hoc with Holm correction: CG vs SM P = 9.8×10□□, ∼9-fold median reduction; ML vs SM P = 1.2×10□□, ∼113-fold median reduction). The ML cocktail produced an additional ∼12-fold reduction in median CFU over CG (ML vs CG P = 0.045) (Fig. 6C).

We used 16S rRNA amplicon sequencing as a community-level check on the CFU result. Interpretation is constrained at low bacterial burden in treatment conditions: after filtering samples with <1,000 total reads (n = 44 retained: ML = 15, CG = 13, SM = 16), residual non-host read depth in treatment samples was low — median 46 microbial reads in ML and 497 in CG — and *P. aeruginosa* read counts in the ML group were at or below reliable detection (median 13 reads). Read-count abundance metrics in treatment samples therefore carry substantial sampling noise; we report read-count abundances and direction of change, and do not attempt community diversity statistics (alpha/beta diversity), which are unreliable at this depth and richness (mean 3.1-3.8 ZOTUs per sample). Within these limits, 16S sequencing was consistent with CFU enumeration: both treatment groups had significantly lower total microbial load and lower absolute *P. aeruginosa* read counts than SM controls, and the ML group was lower than CG on both (e.g. *P. aeruginosa reads*: ML vs SM q = 2.1×10□□, CG vs SM q = 1.4×10□³, ML vs CG q = 1.8×10□³; *Supp. Fig. 20*; *Table S19-S20*). *P. aeruginosa* as a fraction of residual microbial reads did not differ significantly between ML and CG (q = 0.64), a comparison that is underpowered at a median of 46 microbial reads per ML sample. The absolute-burden measures, CFU enumeration and *P. aeruginosa* read counts, consistently favored the ML cocktail.

## Discussion

The clinical promise of phage therapy for multidrug-resistant bacterial infections is tempered by practical implementation challenges. Unlike antibiotics, phages exhibit strain-level specificity, which preserves the host microbiome but complicates therapeutic product development, as no single formulation can achieve universal coverage within a species. Bacterial hosts can rapidly evolve resistance, necessitating multi-phage cocktails, yet rational cocktail design has historically required resource-intensive iterative screening using plaque assays and growth curves across candidate strain-phage combinations. Machine learning offers a complementary strategy for prioritizing candidate therapeutic phages directly from genome sequence. Here, we applied this framework to *P. aeruginosa*, a WHO critical-priority pathogen responsible for high-mortality infections in vulnerable patient populations.

This study establishes the most comprehensive phage-host interaction resource assembled for *P. aeruginosa* to date and demonstrates that machine learning models trained on systematically curated large-scale interaction data can guide therapeutically active phage cocktail design with in vivo validation. The 9,405 interaction data points generated here, spanning 95 genomically diverse phages across 20 genera and 99 clinically representative strains, enabled development of predictive models that match a non-deployable degree-only reference on per-strain susceptibility prediction — recovering the per-strain ranking obtainable from degree structure alone (median AUROC 0.86 for GenoPHI vs reference 0.86). In a single-strain *in vivo* head-to-head test, an ML-designed three-phage cocktail produced a greater bacterial burden reduction in a murine wound infection model than an expert-designed cocktail formulated using classical receptor-diversity principles (∼12-fold median CFU reduction, q = 0.045), supporting the use of computational prediction as a viable strategy for therapeutic phage selection.

### Phage genus is the dominant determinant of host range

Phage genus dominated host range prediction (η² = 0.823, ε² = 0.850, p = 1.5 × 10□¹□), reflecting the conserved genomic architecture and shared receptor-binding strategies that define each genus. However, substantial within-genus variation, exemplified by Ab29 and PSA69 (both *Pbunavirus*) infecting 73 and 29 strains respectively, establishes that genus classification provides insufficient resolution for therapeutic selection and that strain-level characterization remains essential.

Genome size, CDS count, and tRNA content correlated with host range in univariate analyses, consistent with ^57–59^ observations linking large genomes and tRNA content to broader infection capacity. Within *P. aeruginosa*, however, these associations largely disappeared after controlling for genus, indicating that genome-content effects primarily covary with genus identity rather than acting as independent predictors of host range. Together, these results suggest that prioritization by genome size or tRNA content functions largely as a proxy for genus selection rather than an independent heuristic for broad host range.

### Surface receptor architecture dominates infection outcomes

The primacy of bacterial surface determinants in governing phage susceptibility emerged as a central theme. O-antigen serotype was the dominant unique predictor of both the magnitude of strain susceptibility (Type III ANOVA partial η² = 0.243, p = 0.019) and the pattern of which specific phages infect each strain (phylogeny-controlled PERMANOVA R² = 0.218, p < 0.001). Defense systems, anti-defense systems, and prophage burden influence specific interactions rather than generally decreasing susceptibility, consistent with the sequence-specific recognition that defines CRISPR-Cas, restriction-modification, and abortive-infection systems.

Phylogenetic signal analyses further suggested that compatibility determinants vary substantially in their phylogenetic structure. Anti-defense systems and prophage burden showed essentially no phylogenetic signal, consistent with horizontal mobility of these elements, while defense-system content and overall susceptibility showed only moderate phylogenetic structure. Together with the modest phylogenetic contribution in the multivariate models, these results indicate that strain-specific genomic characterization is likely more informative for susceptibility prediction than phylogenetic placement alone.

Two methodological points qualify these conclusions. First, the high local multiplicity of infection in the clearance-based spot assay predominantly captures receptor-level compatibility and may underweight defense-system effects that manifest as titer reduction rather than binary blocking; quantitative liquid-culture or in vivo formats may shift the relative contributions. Second, the interaction matrix itself was significantly anti-nested at both global and within-genus scales (NODF = 60.5, Z = −4.0, p < 0.001; five of 10 well-sampled genera individually anti-nested after Benjamini-Hochberg correction), with no significant modularity (Q = 0.024, p = 0.14). Broad-range and narrow-range phages therefore target non-overlapping host subsets rather than a hierarchical generalist-includes-specialist architecture, indicating that phage redundancy within a cocktail is best measured by which strains a phage covers rather than how many.

### Curated and learned features capture overlapping biology

The comparison between models trained on manually curated genomic features (serotype, defense and anti-defense counts, prophage burden, genome size, gene count) and unsupervised GenoPHI pangenome features revealed no significant performance difference at the median (paired Wilcoxon p ≥ 0.24 for AUROC and AUPR at both per-strain and per-phage aggregation). The GenoPHI top features are predominantly LPS biosynthesis genes, the molecular determinants of the categorical serotype feature, indicating that the unsupervised pipeline largely recapitulates serotype at gene resolution rather than identifying fundamentally new predictive biology. On hard strains, GenoPHI did not statistically outperform non-ML baselines computed from the same interaction matrix, specifically a greedy set-cover and a promiscuity heuristic. We attribute this to the high positive rate of the dataset rather than to a limitation of the model. At 33.3% positive interactions, this panel is dense by phage-host standards. Among published matrices scored by spot or plaquing assays, *Pseudomonas* appears characteristically permissive, with this dataset (33.3%) and an independent *Pseudomonas* matrix (36.2%) both exceeding the largest comparably scored *E. coli* matrix (approximately 21%)^21,36,37^. When infections are this common, most strains are infectable by many phages, and a cocktail of broadly infective phages, which both the set-cover and promiscuity heuristics select, succeeds frequently on base rate alone, leaving little room for a model to improve on. The advantage of model-based prediction over such degree heuristics emerges when positive interactions are rare. There, generically promiscuous phages frequently miss, and correctly predicting which specific phage matches which specific strain becomes decisive. For *P. aeruginosa* specifically, the interaction matrix and the model are complementary deliverables. The matrix is a community resource characterizing this phage panel, and the model is what extends that resource to clinical isolates not represented in it.

### In vivo validation of ML-guided cocktail design

The head-to-head in vivo comparison showed that both ML and CG cocktails substantially reduced PAO1 wound burden relative to vehicle (CG vs SM q = 9.8 × 10□□, ∼9-fold; ML vs SM q = 1.2 × 10□□, ∼113-fold) and that the ML cocktail produced a greater median bacterial-burden reduction than CG by CFU enumeration (∼12-fold, q = 0.045), with 16S sequencing showing the same direction — significantly lower *P. aeruginosa* read counts in ML than CG (q = 1.8 × 10□³). PAO1, a standard reference strain widely used in *P. aeruginosa* infection models, was held out from feature selection, model training, and cocktail-design optimization across all cross-validation folds, so the model’s selection reflects generalization from the remaining 98 strains to a previously unseen isolate. This single-strain test cannot establish the general superiority of ML-guided design over expert-driven heuristics, but it demonstrates that genomics-based prediction trained on a panel of related strains can identify therapeutically active cocktails for an isolate whose interaction profile is unknown to the model, achieved with no prior receptor characterization or resistance profiling.

### Limitations

Several limitations frame these results. (i) The clearance-based spot assay used to generate the interaction dataset captures receptor-level compatibility well but does not resolve quantitative differences in burst size, adsorption kinetics, or infection efficiency, and may underweight defense-system effects that manifest as titer reduction rather than binary blocking. (ii) The *in vivo* comparison is a single-strain, single-time-point test in one animal model using a laboratory reference strain (PAO1) and cocktails of fixed three-phage composition; prospective validation across multiple unseen clinical isolates, dose regimens, and infection contexts will be required to establish the generalization of the framework. (iii) The 16S analysis on the in vivo samples was constrained by host dominance (>96% host reads in ML) and low residual microbial depth, and the *Pseudomonas*-fraction-of-microbial metric did not differ between ML and CG (q = 0.64) even though absolute and total-fraction measures did; we treat 16S as a check that the CFU result is consistent with the broader microbial readout, not as a statistically independent confirmation. (iv) Mouse-side data were collected exclusively in male BALB/c mice; sex-stratified validation will be needed before clinical translation.

## Conclusions

Despite these limitations, this work demonstrates that large-scale, systematically constructed phage-host interaction datasets can train machine learning models capable of predicting therapeutic candidates from genome sequences alone. In vivo validation showed that an ML-designed cocktail produced greater bacterial-burden reduction than an expert-designed cocktail against a held-out *P. aeruginosa* strain. This single-strain demonstration establishes that genomics-based prediction can identify therapeutically active cocktails for isolates whose interaction profiles are unknown to the model.

Our results support a receptor-dominant model of strain susceptibility in *P. aeruginosa*, in which surface receptor composition is the primary determinant of compatibility while intracellular defense and prophage systems shape phage-specific interaction outcomes within permissive receptor backgrounds. This architecture makes genome-based susceptibility prediction feasible because many receptor-associated loci are evolutionarily constrained and readily annotated. The principal practical contribution is the integrated workflow itself, encompassing a curated, genomically diverse phage library; the largest systematic *P. aeruginosa* interaction dataset to date; rigorous multivariate analyses identifying serotype and receptor architecture as the dominant strain-side compatibility determinants; and a trained predictive model that generalizes from a panel of characterized strains to identify therapeutically active cocktails for held-out isolates. Together, these establish a data-driven approach for matching phages to genetically diverse clinical isolates that does not require de novo empirical resistance profiling for each new strain.

The receptor-dominant architecture revealed here opens additional avenues for mechanistic investigation and precision applications. Systematic receptor characterization across this phage library, analogous to recent efforts mapping receptor usage in *E. coli* phage collections^60^, would directly connect phage genotype to bacterial phenotype at the molecular level. Integrating host range prediction models like those developed here with computational approaches for predicting receptor specificity from genome sequence alone would create a powerful combinatorial framework. Such integration would enable the identification of phages predicted to infect a target strain and simultaneously specify which bacterial surface receptor they engage, all from genomic data without prior empirical characterization. This dual prediction capability would transform strain-specific microbiome engineering, where phages with computationally defined receptor specificities could be deployed to selectively remove or suppress target bacterial populations while preserving commensal flora. The combination of predictive models identifying candidate phages and computationally inferred receptor profiles specifying their precise molecular targets would extend phage applications beyond infection treatment to deliberate ecological manipulation of complex microbial communities, with applications ranging from targeted pathogen clearance to precision microbiome remodeling in agricultural and clinical contexts.

## Materials and methods

### Bacterial strains and bacteriophages

*Pseudomonas aeruginosa* PAO1 was obtained from the Manoil Lab (University of Washington); *P. aeruginosa* PA14 was obtained from ATCC. The MRSN panel strains were obtained from the Walter Reed Army Institute of Research ^15^. We have removed strains MRSN1906 from our analysis because it was missing in our copy of collection and strains MRSN3705, MRSN8141, and MRSN6241 because these strains are from PA7 clade and are now recognized as P. *paraeruginosa* ^61^. Strains were cultured in LB (Lennox;10 g/L Tryptone, 5 g/L NaCl, 5 g/L yeast extract) or LB agar [LB(Lennox with 1.5% Bacto agar)] at 37°C.

Phages were shared by Dr. Daria Van Tyne (University of Pittsburgh), Dr. Joe Bondy-Denomy (UCSF), Dr. Catherine M. Mageeney (Sandia National Laboratories) and Dr. Christine Pourcel (Université Paris-Saclay) or isolated in this study. Clonal isolates of phages were also streak purified from Phago-Pyo (batch # M1-1301) obtained from Eliava Institute, Tbilisi, Georgia. Phage titers were determined by ten-fold serial dilution in SM buffer and spot plating 2 µL on 0.5% top agar (10 g/L Tryptone, 10 g/L NaCl, 5 g/L Bacto agar) lawns supplemented with 5 mM CaCl□ and 5 mM MgSO□

### Phage enrichment and amplification

Phage enrichment employed two parallel strategies designed to maximize the genomic diversity of isolated phages ^21^. In the pooled enrichment approach, 3ml each of 17 wastewater samples collected at multiple time points from four California facilities - Oro Loma Sanitary District, Watsonville Wastewater Plant, Santa Cruz Wastewater Treatment Facility and East Bay Municipal Utility District were combined (51 ml total). Similarly, 20 *P. aeruginosa* MRSN strains carrying susceptible AMR phenotypes were grown overnight in individual tubes. 100 ul of each culture was pooled into 50 ml of 2x LB supplemented with CaCl2 and MgSO4 (10 mM each). The pooled wastewater and pooled bacterial culture were combined in a single culture flask and incubated overnight at 37 C with shaking at 200 rpm. The resulting enrichment was clarified by centrifugation (10,000 x g, 10 min) and filter-sterilized. For phage isolation, 10 ul of clarified enrichment was spotted on the lawn of individual *P. aeruginosa* MRSN strains and spread with sterile paper. Plaques of distinct morphologies were picked with a sterile P200 pipet tip, transferred to a microcentrifuge tube with 500 ul SM buffer and subjected to two additional rounds of spotting and streaking to obtain a clonal isolates with uniform plaque morphology. Where morphological heterogeneity persisted after three rounds, plaques of each distinct type were picked and purified separately to confirm that each represented a stable, distinct phage.

For the standard single-host enrichment strategy, 2 mL of individual environmental samples were combined with an equal volume of 2× LB supplemented with 10 mM CaCl□ and MgSO□, and inoculated with a 1:50 dilution of overnight *P. aeruginosa* PAO1 or PA14 cultures in 24-well deep-well plates sealed with Breathe-Easy® membrane (Sigma-Aldrich). Plates were incubated at 37°C, with shaking at 200 rpm. Enrichments were processed and phage plaques isolated as described above.

For amplification, a 1:50 dilution of an overnight P. aeruginosa culture in 30 mL LB supplemented with 5 mM CaCl□ and MgSO□ was incubated at 37°C, 200 rpm for approximately 30 min to reach early exponential phase, then inoculated with 100 µL of purified phage stock. After 3 h of co-incubation, the culture was harvested, clarified by centrifugation (10,000 × g, 10 min), and filter-sterilized. Amplification was confirmed by titering the filtrate as described above.

### Phage DNA extraction and sequencing

Phage DNA was extracted from amplified phage lysates using a modified Wizard DNA purification kit (Promega) as described previously ^62^. NGS library preparation and Illumina Sequencing for phage genomes were performed by Neochromosome Inc. Prior to NGS library preparation, phage DNA was treated with RNase I. Phage DNA was then fragmented and partial Illumina adapters were added by tagmentation with Illumina TDE1 enzyme. Nextera compatible unique dual indices were added and sample libraries were amplified by PCR. Resulting NGS libraries were sequenced on the Illumina NextSeq2000 platform to produce paired-end 300bp reads at a target coverage depth of 10x per sample.

### Phage genome assembly and annotation

Adapters were first trimmed from raw reads using cutadapt v4.7^63^ within trim_galore v0.6.10 (default parameters)^64^, and reads or regions of lower quality were identified and trimmed with bbduk v39.06 (qtrim=r, trimq=20, maq=20) ^65^. The library was then normalized to 100x coverage with bbnorm v39.06 (target=100 min=2)^65^, before assembly with SPAdes v3.15.5 (--isolate -k 21,33,55,77,99,121)^66^. The resulting contigs of ≥2,000 bp were screened for phage genomes with geNomad v1.7.6 (--conservative)^67^, and their quality and completeness were evaluated with CheckV v1.0.3 (default parameters)^68^. For each library, a single phage genome estimated to be 100% complete was obtained, and selected for further characterization.

Phage genomes were annotated with Pharokka v 1.6.1^69^. Specifically, coding sequences (CDS) were predicted with PHANOTATE^70^, tRNAs were predicted with tRNAscan-SE 2.0 ^71^, tmRNAs were predicted with Aragorn^72^ and CRISPRs were predicted with CRT^73^. Functional annotation was generated by matching each CDS to the PHROGs^74^, VFDB^75^ and CARD^76^ databases using MMseqs2^77^ and PyHMMER^78^. Contigs were matched to their closest hit in the INPHARED database^79^ using mash^80^. Annotations of coding genes were improved with PHOLD v 0.2.0^81^.

### Genomic characterization of phages

Phage genome similarity was estimated using the proteomic equivalence quotient (PEQ) metric calculated by PhamClust^41^. Briefly, PhamClust uses gene families inferred by PhaMMSeqs^41,82^ to determine the proportion of genes shared by two genomes. For each pair of homologous proteins, PhamClust calculates the amino acid identity value. The PEQ score for a pair of genomes is calculated as a product of the proportion of shared genes and average amino acid identity, thus PEQ values are ranging from 0 to 1. A score of 0 indicates there is no similarity in the proteins encoded in the two phage genomes, whereas 1 indicates that all proteins encoded in these two phage genomes are identical^41^. Although for genome similarity analysis PEQ values are comparable to nucleotide based BLASTN identity, we preferred PEQ to compare two phage genomes because PhamClust is computationally efficient and is easier to use for large numbers of genomes^41^. For this work, we clustered the PEQ distance matrix with 0.95 threshold. The PEQ similarity matrix of 95 phages was converted into a distance matrix in PHYLIP format and a distance-based phylogeny tree was calculated ^83^.

### Bioinformatic analysis of *P. aeruginosa* genomes

Genome assemblies of *P. aeruginosa* were downloaded from NCBI Datasets^84^. Antibiotic resistance genes and phage defence systems were predicted by the GenomeDepot annotation pipeline; per-strain defense-system counts are provided in *Dataset S5*^85^. For strains with no serotype data available, O-types were inferred *in silico* using PAst; predicted serotypes are provided in *Dataset S6*^86^. Prophages were predicted with geNomad; per-prophage predictions are in *Dataset S7* and prophage counts by viral realm in *Dataset S8*^67^. The phylogenetic tree of *P. aeruginosa* strains was calculated by FastTree ^87^ using nucleotide multiple alignment of the core pangenome produced by PPanGGOLiN ^88^.

### Phage host range determination

Phage-host interaction assays were performed by spotting 2 µL of each phage onto *P. aeruginosa* lawns using a semi-automated Rainin MicroPro 20 system. The 95 phages were tested against 99 strains, generating 9,405 interaction datapoints. Applied phage input ranged from 1E3 to 2E9 PFU. Interactions were assessed based on host clearance, where clearance phenotypes can result from mechanical or enzymatic bacterial lysis, growth inhibition through toxicity, or phage replication and productive lysis. With individual phage titers, the mechanisms of clearance are challenging to distinguish, but all represent clinically-relevant bactericidal and bacteriostatic phenotypes. Interactions were therefore scored as 0 (no lysis), 1 (hazy/turbid lysis), or 2 (clear lysis), averaged across two biological replicates, and classified as positive when the mean score reached ≥1.

### Statistical analysis

#### Dataset and binarization

The phage–host interaction dataset comprised 99 *P. aeruginosa* strains tested against 95 phages, yielding an interaction matrix of 9,405 pairs. Interactions were originally scored on a three-point scale (0 = no infection, 1 = turbid/partial lysis, 2 = clear lysis). Because values 1 and 2 both represent lytic host range with no meaningful biological distinction between lysis intensities in this dataset, all non-zero values were collapsed to 1, producing a binary classification target. The resulting class balance was 66.7% negative (6,273 pairs) and 33.3% positive (3,132 pairs). The pair-level dataset was constructed by melting the interaction matrix and merging all phage and strain features, producing one row per phage-host combination.

#### Univariate phage-side analyses

Host range was defined as the number of strains infected by each phage (n = 95). Spearman rank correlations were computed between host range and each continuous phage feature (genome size, CDS count, tRNA count). To test for genus-level differences in host range, a Kruskal–Wallis H test was applied to the 14 genera containing ≥2 phages (88 phages total; 7 singleton genera excluded). Effect sizes were reported as both epsilon-squared (ε² = (H − k + 1) / (n − k)) and eta-squared (η² = (H − k + 1) / (N + 1)), where k is the number of groups and N is the total sample size. Post-hoc pairwise comparisons used Dunn’s test with Bonferroni correction.

#### Univariate strain-side analyses

Susceptibility was defined as the number of phages infecting each strain (n = 99). The effect of O-antigen serotype was tested by Kruskal–Wallis across 13 serotypes. Spearman rank correlations were computed between susceptibility and each strain-level feature (prophage count, defense gene count, antidefense gene count, total gene count).

#### Partial correlations and per-genus tests

To distinguish independent feature contributions from features that covary with phage genus, partial Spearman correlations between continuous phage features and host range were computed controlling for genus identity using pingouin.partial_corr (Python). Per-genus Spearman correlations were additionally computed within each of the 10 well-sampled genera (≥5 phages); p-values were adjusted using the Benjamini–Hochberg false discovery rate procedure.

#### Collinearity diagnostics

Multicollinearity among predictors was assessed via variance inflation factors (VIF) for the Type III ANOVA design matrix (statsmodels.stats.outliers_influence.variance_inflation_factor) and generalized variance inflation factors (GVIF) for the PERMANOVA design matrix (car::vif in R). VIF/GVIF values < 5 were taken to indicate moderate, interpretable collinearity.

#### Type III ANOVA on susceptibility magnitude

The magnitude of strain susceptibility was modeled by Type III ANOVA with O-antigen serotype, defense system count, anti-defense count, prophage burden, and total gene count as predictors, using statsmodels.formula.api.ols with typ=3. Partial η² and 95% confidence intervals are reported per predictor; overall model adequacy was assessed via the F-test and residual diagnostics.

#### Phylogeny-controlled PERMANOVA on susceptibility pattern

The pattern of strain susceptibility in which specific phages infect each strain, was analyzed by PERMANOVA on Jaccard distances of per-strain infection profiles. To control for strain phylogenetic structure, the first 10 principal coordinates of neighbor matrices (PCNM eigenvectors) derived from the marker-gene phylogeny were entered as conditioning variables. PERMANOVA was performed using vegan::adonis2 (R) with 9,999 permutations and by=“margin” to obtain Type III unique-variance partitioning. Partial R² values reflect variance uniquely attributable to that predictor after controlling for all other predictors and PCNM phylogenetic structure. The PERMANOVA was computed on the binarized interaction matrix, consistent with the binary target used throughout; an analysis on raw 0/1/2 scores gave near-identical results (all focal-predictor R² within 0.002).

#### Phylogeny-controlled dbRDA on phage infection profiles

Phage-side variance in infection profiles was partitioned by distance-based redundancy analysis (dbRDA) on Jaccard distances between phages, using vegan::capscale. The PEQ-based phage phylogeny was decomposed into PCNM eigenvectors; the first 10 were entered as conditioning variables, and phage genus was the focal predictor. Unique-variance contributions were estimated by comparing nested models with and without genus. Significance of each term was assessed via 9,999 permutations. Both phylogeny-controlled analyses are conditioned on the first 10 PCNM eigenvectors. To confirm the conclusions do not depend on this count, the dbRDA and PERMANOVA were re-run across PCNM conditioning sets of 5, 10, and 20 eigenvectors and with forward selection (vegan::ordiR2step; Table S15). Genus (phage-side) and O-antigen serotype (strain-side) remained significant across every variant; defense-system count was also significant throughout, while anti-defense count and prophage burden were significant at PCNM counts ≤ 10 and under forward selection but not when 20 axes were retained.

#### Phylogenetic signal

Phylogenetic signal for each strain-level feature (defense system count, anti-defense count, prophage burden, total susceptibility) was quantified using Pagel’s λ on the marker-gene phylogeny, computed via phytools::phylosig (method=“lambda”, test=TRUE) in R. Likelihood-ratio tests against λ = 0 were used to assess significance.

#### Network topology — nestedness and modularity

Phage–host interactions were represented as a binary bipartite matrix (hosts × phages). Nestedness was quantified using the NODF metric following the BiWeb framework^89–91^, adapted to Python 3. Prior to calculation, matrices were ordered by decreasing row and column degree, and nestedness was computed using strict decreasing fill with separate contributions from rows and columns. Significance was assessed using a degree-preserving swap null model, with 1,000 null matrices generated by iterative edge swaps preserving marginal totals; observed values were compared to the null distribution using Z-scores and empirical p-values. Both nestedness and anti-nestedness directions were tested. Per-genus anti-nestedness was assessed by computing NODF separately within each of the 10 well-sampled genera (≥5 phages) and comparing each observed NODF to a genus-specific degree-preserving null distribution (1,000 replicates); one-sided p-values were adjusted across genera using Benjamini–Hochberg correction. Modularity was estimated using the LP-BRIM algorithm (label propagation followed by bipartite recursive improvement) implemented via the adapted BiWeb code, with ≥50 random initializations retaining the partition with the highest modularity score. Modularity significance was evaluated using the same degree-preserving null model (1,000 replicates).

#### Jaccard distances and ordination

Jaccard distance matrices were computed from binary interaction scores using vegan::vegdist. Principal coordinate analyses (PCoA; k = 10) used stats::cmdscale. Phylogenetic and PEQ-based trees were converted to PCNM eigenvectors for use as conditioning variables in dbRDA and PERMANOVA models as described above. PCoA, dbRDA, and heatmap visualizations were generated with ggplot2.

### Machine learning modeling

#### Models

We trained three deployable CatBoost classifiers on the binary interaction matrix under 20-fold strain-based nested cross-validation, with 10% of strains entirely withheld from feature selection, training, and hyperparameter tuning in each fold:

1. Genus+Serotype: a minimal-feature baseline using only phage genus (one-hot encoded) and host O-antigen serotype (one-hot encoded).
2. Curated baseline: adds strain-level defense system count, anti-defense system count, prophage burden, and total gene count, and phage-level genome size, CDS count, and tRNA count.
3. GenoPHI: replaces the curated bioinformatic features with pangenome-derived MMSeqs2 protein-cluster features and applies the model-based feature selection procedure described previously^45^.

Within each fold, an ensemble of 50 CatBoost classifiers was trained on internal 80/20 splits of the training strains with early stopping on the internal held-out split. Hyperparameters for curated feature models were fixed at CatBoost defaults (1,000 iterations, learning rate 0.1, depth 6, balanced class weights). Predictions on the held-out validation strains were summarized as the median confidence across ensemble members.

#### Non-deployable degree-only reference

As a degree-only reference, we computed a marginal model that assigns each phage–strain pair the product of full-matrix phage breadth and strain susceptibility. Because it requires knowledge of the test strain’s interactions, this reference is not deployable for novel isolates; it characterizes the performance attainable from first-order degree structure alone, not a theoretical upper bound on phage–host prediction.

#### Performance evaluation

Performance was evaluated under three aggregation views: per-fold (predictions pooled within each of the 20 folds), per-strain (predictions pooled across all folds in which each strain appeared, one AUROC/AUPR per strain), and per-phage (predictions pooled across all folds in which each phage appeared, one AUROC/AUPR per phage). For each view we computed AUROC, AUPR, and Matthews correlation coefficient (MCC). Pairwise model comparisons used paired Wilcoxon signed-rank tests within each aggregation view and metric. Variance differences across models were assessed with the Brown–Forsythe test.

#### Difficulty-axis stratification

To localize where model performance differences are concentrated, per-entity AUROC and AUPR were binned along four difficulty axes: (1) phylogenetic novelty, defined as patristic distance to the nearest training-fold neighbor; (2) phenotypic divergence, defined as Jaccard distance from the entity’s five phylogenetic nearest neighbors in interaction profile; (3) marginal difficulty, defined as 1 minus the degree-only reference AUROC, capturing failures of degree-based prediction; (4) full-matrix degree, defined as total susceptibility (per-strain) or host range (per-phage) in the full interaction matrix. Each entity was assigned to a tercile (low/mid/high) per axis, and per-tercile performance was compared across models with Mann–Whitney U tests with BH correction.

#### Cocktail design and benchmarking

For each deployable model, candidate 3- and 5-phage cocktails were selected per held-out strain using model confidence scores under HDBSCAN-derived receptor-diversity constraints (clustered on GenoPHI’s selected pangenome features for the GenoPHI model, and on the training-fold interaction matrix for the curated and Genus+Serotype models). Cocktails were additionally compared against two non-ML baselines: (i) a greedy set-cover that iteratively selects the phage maximizing additional strain coverage on the training-fold interaction matrix; and (ii) a promiscuity baseline that hierarchically clusters phages on the training-fold interaction matrix and selects the most-infectious phage per cluster. Cocktail success was defined as ≥1 phage in the top-k recommendation that infects the held-out strain. Strains were stratified into quartiles by full-matrix susceptibility; top-k success was compared between models within each quartile using McNemar’s test.

#### Cocktail selected for in vivo testing

For the GenoPHI cocktail tested in vivo, all training and feature selection was performed on the 98 strains other than PAO1; the trained model was applied to predict PAO1 susceptibility against all 94 remaining phages, and the cocktail-design procedure above was applied to select three phages.

#### SHAP analysis

Shapley Additive exPlanations (SHAP) values were computed for the GenoPHI model using the TreeSHAP algorithm. The 25 features with largest mean absolute SHAP value were annotated with gene functions to compare against known mediators of phage-host interaction in *P. aeruginosa*.

### Generation of phage cocktails to test efficacy in murine model

Once the first two phages (phiKZ and phiKMV) were selected for the phage cocktail, a third phage potentially utilizing a different receptor for infection was desired. For this, we generated mutants of PAO1 that are not infected by phiKZ or phiKMV, independently. Lysates of phages phiKZ or phiKMV were diluted ten-fold to a magnitude of 10^-3^. 100 ul of each dilution was incubated with 100 ul of PAO1 ON culture for 10 min at RT. 100 ul of this phage-host mix was then spread into a LB agar plate and incubated overnight at 37C. An isolated colony was picked from the plate with the least diluted phage stock. This colony was streaked and a single colony that is potentially resistant to phiKZ infection (PAO1_phiKZ_^R^) or phiKMV infection(PAO1 ^R^) was picked. Phage phiKZ or phiKMV was spotted on the lawns of these two PAO1 mutants to confirm that the mutant was resistant to phiKZ or phiKMV.

To screen for the third candidate phage, the phage panel was spotted on the lawn of PAO1_phiKZ_^R^ or PAO1 ^R^ and the infection outcome was scored as described above. Only the phages that showed infection score of 2 on both PAO1phiKZR or PAO1phiKMVR were selected. These phages belong to genera *Nankokurvirus* (Ab03, Ab04, Ab06, Ab17, Ab11), *Pbunavirus* (Ab27, DP8, DP10, F8, Ab29, PB1), *Pakpunavirus* (PAO1_MA12), and *Bruynoghevirus* (DP9, DP24). Among these four genera, phages belonging to *Pakpunavirus* and *Pbunavirus* appear to have a broad host range similar to phages belonging to *Phikzvirus* and *Phikmvvirus*. Phage PAO1_MA12 belonging to *Pakpunavirus* genus has a larger genome (∼92kb) compared to phages from *Pbunavirus* genus (∼66kb); Hence, PAO1_MA12 was chosen as the third component phage of this cocktail.

### Murine wound infection model

To assess the therapeutic efficacy of phage cocktails in vivo, we used an established murine wound infection model^92–95^. All procedures were reviewed and approved by the Animal Welfare and Research Committee (AWRC), the Institutional Animal Care and Use Committee at Lawrence Berkeley National Laboratory, under protocol number 271034. Male BALB/c mice (10–16 weeks of age; Jackson Laboratory West) were single-housed in disposable ventilated cages provisioned with Alpha-dri and Enviro-dri bedding and a Shepherd Shack for environmental enrichment. Animals were weighed and monitored daily for health status throughout the experiment.

On the day prior to wounding (Day-1), mice were weighed and anesthetized using isoflurane inhalation. Analgesia was administered subcutaneously at the dorsal caudal region (Ethiqa-XR, 3.25 mg/kg). Approximately 30 minutes after analgesia administration, the dorsal fur was shaved with electric clippers, and the skin was cleaned twice with 70% ethanol, beginning at the intended wound site and moving caudally. Mice were then transferred to a sterile surgical field for wounding.In the surgical field, two full-thickness excisional wounds were generated under aseptic conditions. The dorsal midline was palpated, and the skin was gently elevated with sterile forceps. Two 6 mm circular wounds were created using a sterile disposable biopsy punch. Wounds were photographed alongside a ruler for size reference, then covered with pre-cut 25 mm Tegaderm dressings. Mice were monitored until recovery from anesthesia before being returned to their cages.

On Day 0, mice were anesthetized with isoflurane and weighed. *Pseudomonas aeruginosa* PAO1 was prepared at OD□□_ = 0.5 (target of ∼1.5 x 10□ CFU/mL) after three washes in sterile saline. The solution was diluted to 4 x 10^^6^ CFU/mL. A total of 50 µL PAO1 (2 x 10^^5^ CFU total) suspension was injected into each wound bed through the Tegaderm using an insulin syringe. Two hours post-infection, mice were re-anesthetized and mice received 100 µL of ML or CG bacteriophage cocktail solution (10□ PFU/mL in SM buffer) or SM buffer alone (control) into each wound bed via insulin syringe. Animals were monitored during recovery and returned to their cages.

On Days 1 and 2 post-infection, mice were weighed, and anesthetized with isoflurane. Wounds were treated with 100 µL of phage solution (ML or CG) or SM buffer was injected into each wound bed. Mice were monitored until full recovery from anesthesia before being returned to their cages. On Day 2, animals received a second dose of sustained-release analgesia (Ethiqa-XR, 3.25 mg/kg, SC) administered 72 hours after the initial dose, in accordance with the manufacturer’s guidelines.

On Day 3, mice were weighed and anesthetized with isoflurane. Digital images of the wound sites were captured for documentation and mice were subsequently euthanized by cervical dislocation. Wound tissues were excised with an 8 mm disposable biopsy punch, minced into 1–2 mm fragments, and placed in pre-weighed collection tubes with homogenization beads and 1mL of 1XPBS buffer.

Skin samples were weighed inside the pre-weighed tubes and homogenized in 1 mL sterile PBS (pH 7.4) using zirconia beads in a TissueLyser maximum speed (50Hz) for 5 minutes. Approximately 700-900uL of homogenate was transferred to a 96-well deep well plate. Homogenates were serially diluted in PBS to 10□□, and 100 µL of each dilution was plated in duplicate on Cetrimide agar plates. Plates were incubated overnight at 37 °C, and colony counts were used to determine bacterial burden (CFU per gram of tissue).

Genomic DNA of bacteria present in the skin samples was extracted using Qiagen DNeasy Blood & Tissue Kit. gDNA template was added to a PCR reaction to amplify the V4/V5 16S gene region using the 515F/ 926R primers based on the Earth Microbiome Project primers that have broad coverage for diverse bacteria and archaea^96,97^, but with in-line dual Illumina indexes^98,99^. The amplicons were sequenced on an Illumina MiSeq (Illumina, San Diego, CA, USA) with 2×300□bp Ilumina v3 reagents. Reads were processed with custom Perl scripts implementing PEAR for read merging^100^, USearch^101^ for filtering reads with more than one expected error, demultiplexing using inline indexes and Unoise^102^ for filtering rare reads and chimeras. 16S sequences in the relative abundance table were searched against the RDP database^103^ to assign taxonomy. We were able to assign a taxonomy to >97% of the reads in the 16S amplicon sequencing dataset

### CFU statistical analysis

Log □ □-transformed CFU per gram of tissue were compared across treatment groups (ML, CG, SM) by Kruskal–Wallis test with Dunn’s post-hoc pairwise comparisons and Holm correction. Values below the limit of detection (LOD = 100 CFU/g) were set to the LOD for statistical analysis; this affected 3/16 ML samples and no CG or SM samples, so reported ML-vs-CG and ML-vs-SM fold reductions are conservative lower bounds on the true ML effect. Effect sizes were reported as rank-biserial correlations.

### 16S amplicon analysis

Samples with fewer than 1,000 total reads were excluded, yielding 44 samples (ML, n = 15; CG, n = 13; SM, n = 16). Reads were classified as host or microbial based on Ribosomal Database Project (RDP)-assigned taxonomy; microbial reads were defined as non-host, non-chloroplast/mitochondrial reads. *Pseudomonas* reads were defined as those assigned to genus *Pseudomonas* at ≥80% RDP confidence. We computed per-sample summary metrics including total reads, host reads, microbial reads (host-removed), absolute *Pseudomonas* read counts, *Pseudomonas* as a fraction of total reads, *Pseudomonas* as a fraction of microbial reads, ZOTU richness, and host fraction. Group differences (ML vs CG vs SM) were assessed by Kruskal–Wallis tests with Dunn’s post-hoc multiple comparisons and Benjamini–Hochberg correction across all between-group contrasts; reported q-values reflect BH correction within each metric. Given the host-dominated read pool in treatment samples (>96% host reads in ML) and low residual microbial depth, per-sample diversity statistics (alpha and beta diversity) were not pursued; 16S was treated as a community-context check on the CFU result rather than as a statistically independent confirmation.

## Author contributions

D.P., Conceptualization, Methodology, Investigation, Validation, Formal Analysis, Data Curation, Visualization, Writing - Original Draft, Review & Editing

A. J.C. N., Conceptualization, Methodology, Investigation, Validation, Formal Analysis, Data Curation, Visualization, Writing - Original Draft, Review & Editing

H. S., Methodology, Investigation, Validation, Formal Analysis, Visualization, Data Curation, Writing - Original Draft, Review & Editing

M. A., Investigation, Validation,

S. K. V., Investigation, Validation, Visualization

F. M., Formal Analysis, Data Curation, Visualization

I. M., Investigation, Validation,

M. S.,Investigation, Validation,

B. B.,Investigation, Validation, M.H.,Investigation, Validation,

C. O.,Investigation, Validation,

A.K., Investigation, Software, Formal Analysis, Data Curation, Visualization - Review & Editing H.K.C.,Investigation, Validation, Review & Editing

Y.Y., Investigation, Validation, Methodology, Formal Analysis, Visualization E.S.,Investigation, Validation, Methodology

S.R., Investigation, Software, Formal Analysis, Data Curation, Visualization - Review & Editing A.M.D.,Investigation,Formal Analysis - Review & Editing

J.L.I., Methodology, Investigation, Validation, Formal Analysis, Visualization - Review & Editing A.P.A., Conceptualization, Methodology, Writing - Review & Editing, Supervision, Funding Acquisition, Project Administration

V.K.M., Conceptualization, Methodology, Writing - Review & Editing, Supervision, Funding Acquisition, Project Administration

## Data Availability

Assembled genome sequences for all phages characterized in this study have been deposited in NCBI GenBank under accession numbers [submitted]. Bacterial genome sequences are available under accession numbers [submitted]. Matrix data and analysis outputs are listed in Supplementary Datasets (DOI: 10.5281/zenodo.20276795) and Supplementary Tables described in the Supplementary Materials.

## Code Availability

The GenoPHI software package used for k-mer-based receptor prediction is available at https://github.com/Noonanav/GenoPHI.

## Materials & Correspondence

All other data supporting the findings of this study are available from the corresponding author Vivek K. Mutalik or Adam P. Arkin upon reasonable request.

## Acknowledgments

We thank members of the BRaVE Phage Foundry team, the Arkin lab and the Mutalik lab for discussions, and all of our phage friends who shared phages for this work. We also thank Morgan Price for data processing assistance and Jennifer Kuehl for support with 16S amplicon sequencing.

## References

1. Knight, G. M. et al. Antimicrobial resistance and COVID-19: Intersections and implications. Elife 10, (2021).

2. De Oliveira, D. M. P. et al. Antimicrobial resistance in ESKAPE pathogens. Clin. Microbiol. Rev. 33, (2020).

3. Health Organization, W. Who publishes list of bacteria for which new antibiotics are urgently needed. Saudi Med. J. 38, 444–445 (2017).

4. López-Calleja, A. I. et al. Antimicrobial activity of ceftolozane-tazobactam against multidrug-resistant and extensively drug-resistant Pseudomonas aeruginosa clinical isolates from a Spanish hospital. Rev. Esp. Quimioter. 32, 68–72 (2019).

5. Magill, S. S. et al. Multistate point-prevalence survey of health care-associated infections. N. Engl. J. Med. 370, 1198–1208 (2014).

6. McCarthy, K. L. & Paterson, D. L. Increased risk of death with recurrent Pseudomonas aeruginosa bacteremia. Diagn. Microbiol. Infect. Dis. 88, 152–157 (2017).

7. Letizia, M., Diggle, S. P. & Whiteley, M. Pseudomonas aeruginosa: ecology, evolution, pathogenesis and antimicrobial susceptibility. Nat. Rev. Microbiol. 23, 701–717 (2025).

8. Gordillo Altamirano, F. L. & Barr, J. J. Phage therapy in the postantibiotic era. Clin. Microbiol. Rev. 32, (2019).

9. Young, R. & Gill, J. J. Phage therapy redux—What is to be done? Science 350, 1163–1164 (2015).

10. MacNair, C. R., Rutherford, S. T. & Tan, M.-W. Alternative therapeutic strategies to treat antibiotic-resistant pathogens. Nat. Rev. Microbiol. 22, 262–275 (2024).

11. Koskella, B. & Meaden, S. Understanding bacteriophage specificity in natural microbial communities. Viruses 5, 806–823 (2013).

12. Cui, L. et al. A comprehensive review on phage therapy and phage-based drug development. Antibiotics (Basel) 13, 870 (2024).

13. Nobrega, F. L. et al. Targeting mechanisms of tailed bacteriophages. Nature Reviews Microbiology 16, 760–773 (2018).

14. Tran, V. N. & Burrows, L. L. Steering the course: targeting and exploiting surface receptors in phage therapy. J. Bacteriol. 208, e0037525 (2026).

15. Lebreton, F., et al. A panel of diverse clinical isolates for research and development. JAC Antimicrob Resist 3, dlab179 (2021).

16. Klockgether, J. et al. Genome diversity of Pseudomonas aeruginosa PAO1 laboratory strains. J. Bacteriol. 192, 1113–1121 (2010).

17. Mishra, P., Sahoo, D. & Sahu, M. C. Genetic diversity of Pseudomonas aeruginosa isolated from clinical samples with ISSR molecular marker in a tertiary care teaching hospital. Sci. Rep. 16, 5315 (2026).

18. Klockgether, J., Cramer, N., Wiehlmann, L., Davenport, C. F. & Tümmler, B. Pseudomonas aeruginosa Genomic Structure and Diversity. Front. Microbiol. 2, 150 (2011).

19. Harrington, N. E. et al. Global genomic diversity of Pseudomonas aeruginosa in bronchiectasis. J. Infect. 89, 106275 (2024).

20. Markwitz, P. et al. Genome-driven elucidation of phage-host interplay and impact of phage resistance evolution on bacterial fitness. ISME J. 16, 533–542 (2022).

21. Gaborieau, B. et al. Prediction of strain level phage-host interactions across the Escherichia genus using only genomic information. Nat. Microbiol. 9, 2847–2861 (2024).

22. Müller, D. M. et al. Bacterial Receptors but Not Anti-Phage Defense Mechanisms Determine Host Range for a Pair of Pseudomonas aeruginosa Lytic Phages. Microbiology (2024).

23. Hussain, F. A. et al. Rapid evolutionary turnover of mobile genetic elements drives bacterial resistance to phages. Science 374, 488–492 (2021).

24. Piel, D. et al. Phage-host coevolution in natural populations. Nat. Microbiol. 7, 1075–1086 (2022).

25. Egido, J. E., Costa, A. R., Aparicio-Maldonado, C., Haas, P.-J. & Brouns, S. J. J. Mechanisms and clinical importance of bacteriophage resistance. FEMS Microbiol. Rev. 46, fuab048 (2022).

26. Chan, B. K. et al. Personalized inhaled bacteriophage therapy for treatment of multidrug-resistant Pseudomonas aeruginosa in cystic fibrosis. Nat. Med. 31, 1494–1501 (2025).

27. Fujiki, J., et al. Phage cocktails containing a dual-receptor Phikzvirus suppress resistance evolution in Pseudomonas aeruginosa. Appl. Environ. Microbiol. 92, e0209525 (2026).

28. De Smet, J., Hendrix, H., Blasdel, B. G., Danis-Wlodarczyk, K. & Lavigne, R. Pseudomonas predators: understanding and exploiting phage-host interactions. Nat. Rev. Microbiol. 15, 517–530 (2017).

29. Chan, B. K., Abedon, S. T. & Loc-Carrillo, C. Phage cocktails and the future of phage therapy. Future Microbiol. 8, 769–783 (2013).

30. Strathdee, S. A., Hatfull, G. F., Mutalik, V. K. & Schooley, R. T. Phage therapy: From biological mechanisms to future directions. Cell 186, 17–31 (2023).

31. Kim, M. K. et al. A blueprint for broadly effective bacteriophage-antibiotic cocktails against bacterial infections. Nat. Commun. 15, 9987 (2024).

32. Nikolich, M. P. et al. Pseudomonas aeruginosa phage cocktails: Rational design and efficacy against mouse wound and systemic infection. Antibiotics (Basel) 15, 75 (2026).

33. Yu, X. et al. A review of phage therapy for drug-resistant Pseudomonas aeruginosa infections. Microbiol. Res. 305, 128417 (2026).

34. Essoh, C. et al. Investigation of a large collection of Pseudomonas aeruginosa bacteriophages collected from a single environmental source in Abidjan, Côte d’Ivoire. PLoS One 10, e0130548 (2015).

35. Nordstrom, H. R. et al. Genomic characterization of lytic bacteriophages targeting genetically diverse Pseudomonas aeruginosa clinical isolates. iScience 25, 104372 (2022).

36. Costa, A. R. et al. Accumulation of defense systems in phage-resistant strains of Pseudomonas aeruginosa. Sci. Adv. 10, eadj0341 (2024).

37. Vaitekenas, A. et al. Genomic determinants of phage activity against Pseudomonas aeruginosa: Roles of receptors, defence systems, and anti-defences. bioRxiv 2026.02.22.706922 (2026) doi:10.64898/2026.02.22.706922.

38. Wright, R. C. T., Friman, V.-P., Smith, M. C. M. & Brockhurst, M. A. Cross-resistance is modular in bacteria-phage interactions. PLoS Biol. 16, e2006057 (2018).

39. Noonan, A., et al. Phylogeny-agnostic strain-level prediction of phage-host interactions from genomes. bioRxiv (2025) doi:10.1101/2025.11.15.688630.

40. Malajczuk, C. J. et al. Towards accurate artificial intelligence models for strain-level phage-host prediction. Brief. Bioinform. 27, (2026).

41. Gauthier, C. H. & Hatfull, G. F. PhamClust: a phage genome clustering tool using proteomic equivalence. mSystems 8, e0044323 (2023).

42. Turner, D., Kropinski, A. M. & Adriaenssens, E. M. A roadmap for genome-based phage taxonomy. Viruses 13, 506 (2021).

43. Letunic, I. & Bork, P. Interactive Tree Of Life (iTOL) v5: an online tool for phylogenetic tree display and annotation. Nucleic Acids Res. 49, W293–W296 (2021).

44. Hyman, P. & Abedon, S. T. Bacteriophage host range and bacterial resistance. Adv. Appl. Microbiol. 70, 217–248 (2010).

45. Noonan, A. J. C. et al. Phylogeny-agnostic strain-level prediction of phage-host interactions from genomes. Microbiology (2025).

46. Flores, C. O., Meyer, J. R., Valverde, S., Farr, L. & Weitz, J. S. Statistical structure of host-phage interactions. Proc. Natl. Acad. Sci. U. S. A. 108, E288–97 (2011).

47. Dormann, C. F. et al. Collinearity: a review of methods to deal with it and a simulation study evaluating their performance. Ecography (Cop.) 36, 27–46 (2013).

48. Legendre, P. & Anderson, M. J. DISTANCE-BASED REDUNDANCY ANALYSIS: TESTING MULTISPECIES RESPONSES IN MULTIFACTORIAL ECOLOGICAL EXPERIMENTS. Ecological Monographs 69, 1–24 (1999).

49. Lood, C., Haas, P.-J., van Noort, V. & Lavigne, R. Shopping for phages? Unpacking design rules for therapeutic phage cocktails. Curr. Opin. Virol. 52, 236–243 (2022).

50. Gordillo Altamirano, F. L. & Barr, J. J. Unlocking the next generation of phage therapy: the key is in the receptors. Curr. Opin. Biotechnol. 68, 115–123 (2021).

51. Ranta, K., Skurnik, M. & Kiljunen, S. Isolation and characterization of fMGyn-Pae01, a phiKZ-like jumbo phage infecting Pseudomonas aeruginosa. Virol. J. 22, 55 (2025).

52. Li, Y., et al. A family of novel immune systems targets early infection of nucleus-forming jumbo phages. bioRxiv (2022) doi:10.1101/2022.09.17.508391.

53. Chibeu, A. et al. The adsorption ofPseudomonas aeruginosabacteriophage ÏKMV is dependent on expression regulation of type IV pili genes. FEMS Microbiol. Lett. 296, 210–218 (2009).

54. Peng, S. et al. Understanding phage Receptor-binding protein interaction with host surface receptor: the key for phage-Mediated detection and elimination of Pseudomonas aeruginosa. Eur. J. Clin. Microbiol. Infect. Dis. (2025) doi:10.1007/s10096-025-05262-x.

55. Li, L. et al. Characterization of Pseudomonas aeruginosa phage K5 genome and identification of its receptor related genes: Genome annotation and the receptor of phage K5. J. Basic Microbiol. 56, 1344–1353 (2016).

56. Brown, V. I. & Lowbury, E. J. Use of an improved cetrimide agar medium and other culture methods for Pseudomonas aeruginosa. J. Clin. Pathol. 18, 752–756 (1965).

57. Bailly-Bechet, M., Vergassola, M. & Rocha, E. Causes for the intriguing presence of tRNAs in phages. Genome Res. 17, 1486–1495 (2007).

58. Yoshikawa, G. et al. Xanthomonas citri jumbo phage XacN1 exhibits a wide host range and high complement of tRNA genes. Sci. Rep. 8, 4486 (2018).

59. Leprince, A., Somerville, V., Addablah, A. A., Morency, C. & Moineau, S. Phage host range: determinants, dynamics and applications. Nat. Rev. Microbiol. (2026) doi:10.1038/s41579-026-01306-x.

60. Moriniere, L. et al. Enabling the prediction of phage receptor specificity from genome data. bioRxiv 2026.04.02.716166 (2026) doi:10.64898/2026.04.02.716166.

61. Déraspe, M. et al. Comparative genomics of Pseudomonas paraeruginosa. J. Bacteriol. 207, e0014925 (2025).

62. Summer, E. J. Preparation of a phage DNA fragment library for whole genome shotgun sequencing. Methods Mol Biol 502, 27–46 (2009).

63. Kechin, A., Boyarskikh, U., Kel, A. & Filipenko, M. CutPrimers: A new tool for accurate cutting of primers from reads of targeted next generation sequencing. J. Comput. Biol. 24, 1138–1143 (2017).

64. Krueger, F., et al. FelixKrueger/TrimGalore: v0.6.10 - Add Default Decompression Path. (Zenodo, 2023). doi:10.5281/ZENODO.7598955.

65. BBMap. SourceForge https://sourceforge.net/projects/bbmap (2022).

66. Prjibelski, A., Antipov, D., Meleshko, D., Lapidus, A. & Korobeynikov, A. Using SPAdes DE Novo Assembler. Curr. Protoc. Bioinformatics 70, e102 (2020).

67. Camargo, A. P. et al. Identification of mobile genetic elements with geNomad. Nat. Biotechnol. 42, 1303–1312 (2024).

68. Nayfach, S. et al. CheckV assesses the quality and completeness of metagenome-assembled viral genomes. Nat. Biotechnol. 39, 578–585 (2021).

69. Bouras, G. et al. Pharokka: a fast scalable bacteriophage annotation tool. Bioinformatics 39, (2023).

70. McNair, K., Zhou, C., Dinsdale, E. A., Souza, B. & Edwards, R. A. PHANOTATE: a novel approach to gene identification in phage genomes. Bioinformatics 35, 4537–4542 (2019).

71. Chan, P. P., Lin, B. Y., Mak, A. J. & Lowe, T. M. tRNAscan-SE 2.0: improved detection and functional classification of transfer RNA genes. Nucleic Acids Res. 49, 9077–9096 (2021).

72. Laslett, D. & Canback, B. ARAGORN, a program to detect tRNA genes and tmRNA genes in nucleotide sequences. Nucleic Acids Res. 32, 11–16 (2004).

73. Bland, C. et al. CRISPR recognition tool (CRT): a tool for automatic detection of clustered regularly interspaced palindromic repeats. BMC Bioinformatics 8, 209 (2007).

74. Terzian, P., et al. PHROG: families of prokaryotic virus proteins clustered using remote homology. NAR Genom. Bioinform. 3, lqab067 (2021).

75. Chen, L. et al. VFDB: a reference database for bacterial virulence factors. Nucleic Acids Res. 33, D325–8 (2005).

76. Alcock, B. P. et al. CARD 2020: antibiotic resistome surveillance with the comprehensive antibiotic resistance database. Nucleic Acids Res. 48, D517–D525 (2020).

77. Steinegger, M. & Söding, J. MMseqs2 enables sensitive protein sequence searching for the analysis of massive data sets. Nat. Biotechnol. 35, 1026–1028 (2017).

78. Larralde, M. Pyrodigal: Python bindings and interface to Prodigal, an efficient method for gene prediction in prokaryotes. J. Open Source Softw. 7, 4296 (2022).

79. Cook, R. et al. INfrastructure for a PHAge REference Database: Identification of large-scale biases in the current collection of cultured phage genomes. Phage (New Rochelle) 2, 214–223 (2021).

80. Ondov, B. D. et al. Mash: fast genome and metagenome distance estimation using MinHash. Genome Biol. 17, 132 (2016).

81. Bouras, G., et al. Protein structure informed bacteriophage genome annotation with phold. bioRxiv (2025) doi:10.1101/2025.08.05.668817.

82. Gauthier, C. H., Cresawn, S. G. & Hatfull, G. F. PhaMMseqs: a new pipeline for constructing phage gene phamilies using MMseqs2. G3 (Bethesda) 12, (2022).

83. Lefort, V., Desper, R. & Gascuel, O. FastME 2.0: A comprehensive, accurate, and fast distance-based phylogeny inference program. Mol. Biol. Evol. 32, 2798–2800 (2015).

84. O’Leary, N. A. et al. Exploring and retrieving sequence and metadata for species across the tree of life with NCBI Datasets. Sci. Data 11, 732 (2024).

85. Kazakov, A. & Deutschbauer, A. M. GenomeDepot: data management system for microbial comparative genomics. Bioinform. Adv. 6, vbag027 (2026).

86. Thrane, S. W., Taylor, V. L., Lund, O., Lam, J. S. & Jelsbak, L. Application of whole-genome sequencing data for O-specific antigen analysis and in silico serotyping of Pseudomonas aeruginosa isolates. J. Clin. Microbiol. 54, 1782–1788 (2016).

87. Price, M. N., Dehal, P. S. & Arkin, A. P. FastTree 2--approximately maximum-likelihood trees for large alignments. PLoS One 5, e9490 (2010).

88. Gautreau, G. et al. PPanGGOLiN: Depicting microbial diversity via a partitioned pangenome graph. PLoS Comput. Biol. 16, e1007732 (2020).

89. Almeida-Neto, M., Guimarães, P., Guimarães, P. R., Jr, Loyola, R. D. & Ulrich, W. A consistent metric for nestedness analysis in ecological systems: reconciling concept and measurement. Oikos 117, 1227–1239 (2008).

90. Poisot, T. BiWeb. (Github).

91. Flores, C. O., Valverde, S. & Weitz, J. S. Multi-scale structure and geographic drivers of cross-infection within marine bacteria and phages. ISME J. 7, 520–532 (2013).

92. Bach, M. S. et al. Filamentous bacteriophage delays healing of Pseudomonas-infected wounds. Cell Rep. Med. 3, 100656 (2022).

93. Sweere, J. M. et al. The immune response to chronic Pseudomonas aeruginosa wound infection in immunocompetent mice. Adv. Wound Care (New Rochelle) 9, 35–47 (2020).

94. de Vries, C. R. et al. A delayed inoculation model of chronic Pseudomonas aeruginosa wound infection. J. Vis. Exp. (2020) doi:10.3791/60599.

95. Lin, Y.-H., et al. Dosing and delivery of bacteriophage therapy in a Murine wound infection model. bioRxivorg (2025) doi:10.1101/2024.05.07.593005.

96. Parada, A. E., Needham, D. M. & Fuhrman, J. A. Every base matters: assessing small subunit rRNA primers for marine microbiomes with mock communities, time series and global field samples: Primers for marine microbiome studies. Environ. Microbiol. 18, 1403–1414 (2016).

97. Quince, C., Lanzen, A., Davenport, R. J. & Turnbaugh, P. J. Removing noise from pyrosequenced amplicons. BMC Bioinformatics 12, 38 (2011).

98. Sharpless, W., Sander, K., Song, F., Kuehl, J. & Arkin, A. P. Towards environmental control of microbiomes. bioRxiv (2022) doi:10.1101/2022.11.04.515211.

99. Price, M. N. et al. Mutant phenotypes for thousands of bacterial genes of unknown function. Nature 557, 503–509 (2018).

100. Zhang, J., Kobert, K., Flouri, T. & Stamatakis, A. PEAR: a fast and accurate Illumina Paired-End reAd mergeR. Bioinformatics 30, 614–620 (2014).

101. Edgar, R. C. Search and clustering orders of magnitude faster than BLAST. Bioinformatics 26, 2460–2461 (2010).

102. Edgar, R. UNOISE2: improved error-correction for Illumina 16S and ITS amplicon sequencing. *bioRxiv* (2016) doi:10.1101/081257.

103. Cole, J. R. et al. Ribosomal Database Project: data and tools for high throughput rRNA analysis. Nucl. Acids Res. 42, D633–D642 (2014).

