## Supplementary material for "Comprehensive interaction profiling and machine learning prediction of bacteriophage infectivity across clinically diverse *Pseudomonas aeruginosa*": Suppl. material

**Supplementary Figures	2**

**Supplementary Tables	18**

### **Supplementary Figures**

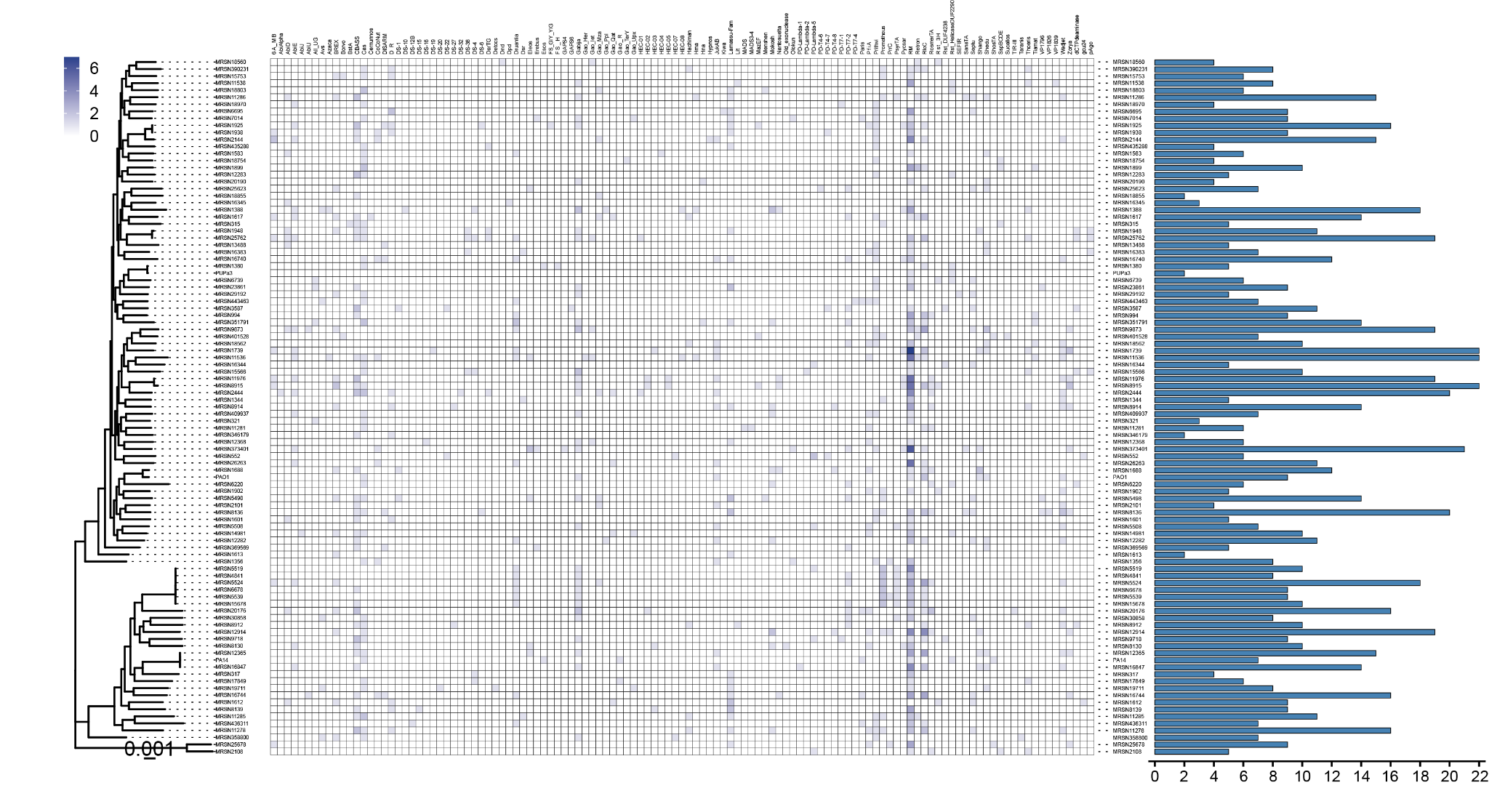

**Supplementary Figure 1. Defense systems identified in MRSN *P. aeruginosa* strain panel genomes**. The heatmap shows the identity and number of individual defense systems present in each strain. Counts are indicated by shades of blue. The bar chart shows the total number of defense systems identified in each strain.

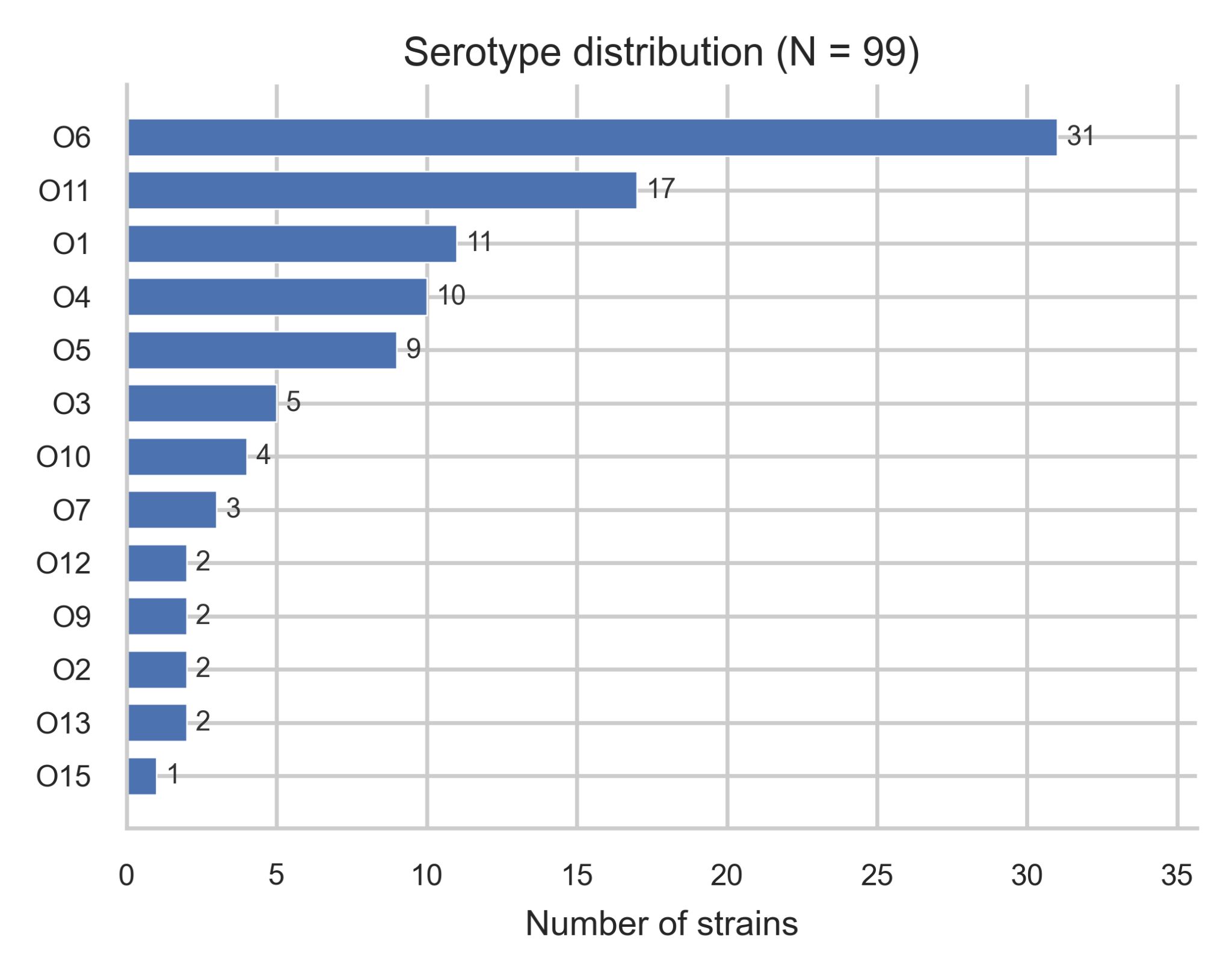

**Supplementary Figure 2.** Serotype distribution across 99 *P. aeruginosa* strains.

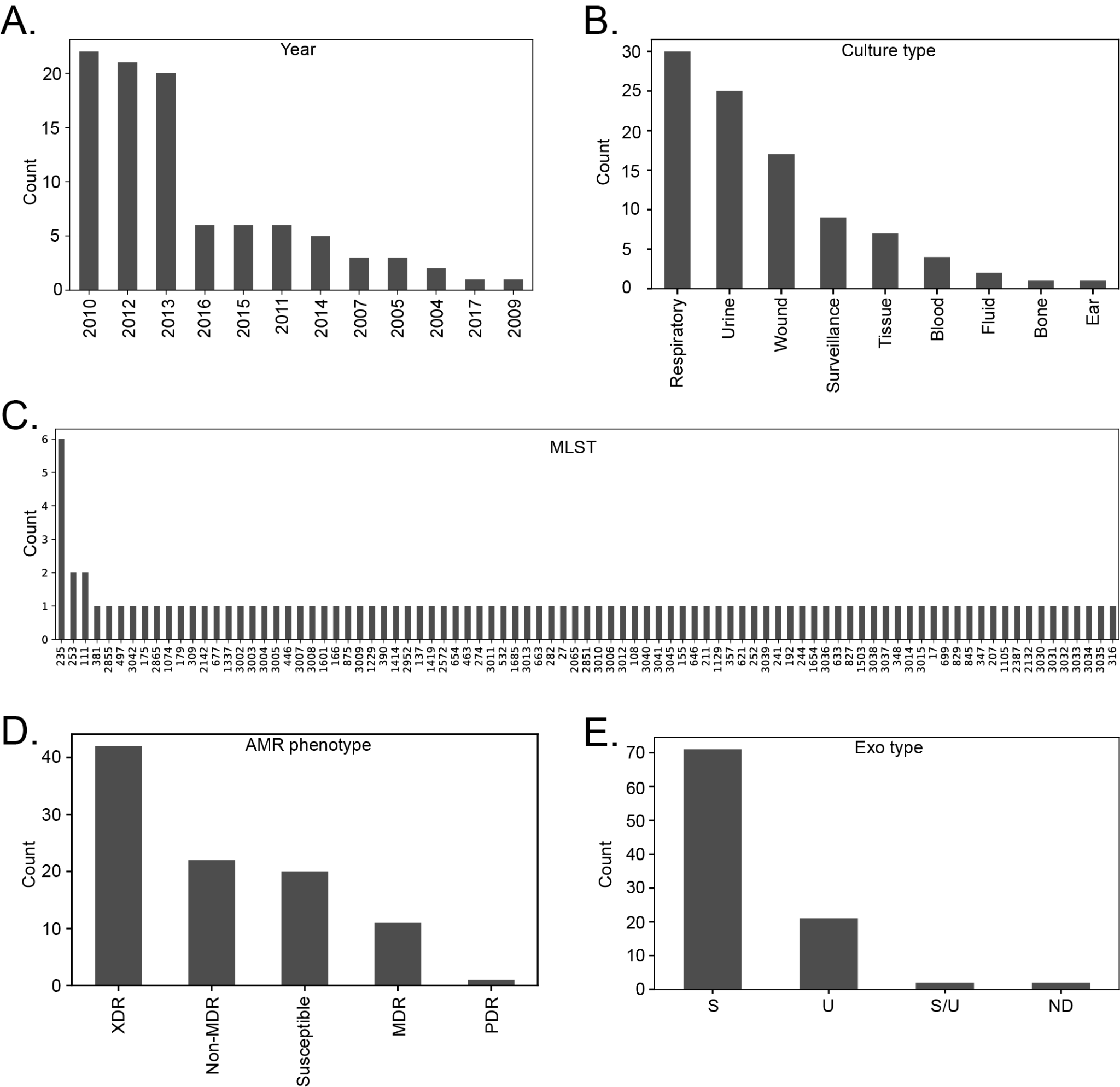

**Supplementary Figure 3. MRSN *Pseudomonas* Strain Panel.** The strains in the MRSN panel were isolated in different years (A) from different clinical specimens (B). (C) MLST profiles of the strains. (D) Antibiotics susceptibility profile. XDR- , MDR- , PDR-	(E) Classification of strains based on the presence of genes encoding the type III secretion system effector protein. S- *exoS*, U- *exoU*, S/U- both *exoS* and *exoU* , ND- not detected in the genome. The data visualized in panels B-F are obtained from earlier study ^1^ and do not include information about *P. aeruginosa* strains PAO1, PA14 and PUPa3.

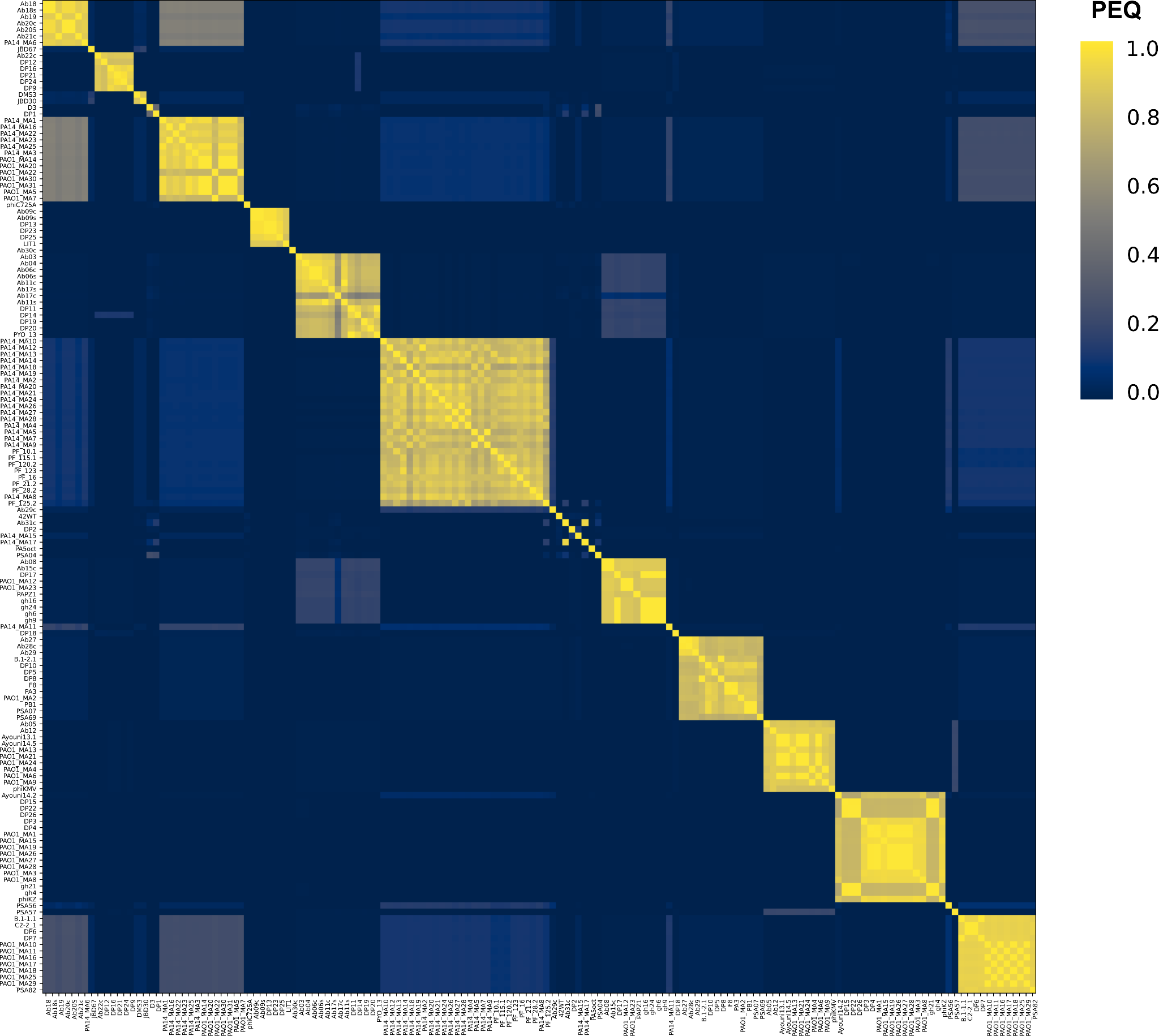

**Supplementary Figure 4. PhamClust comparison of *P. aeruginosa* phages.** All *P. aeruginosa* phages (total **153**) present in our collection were sequenced and analyzed via PhaMMseqs and PhamClust. The heatmap shows the proteomic equivalent quotient (PEQ) of these phages. Phage names are listed along the X- and Y-axes.

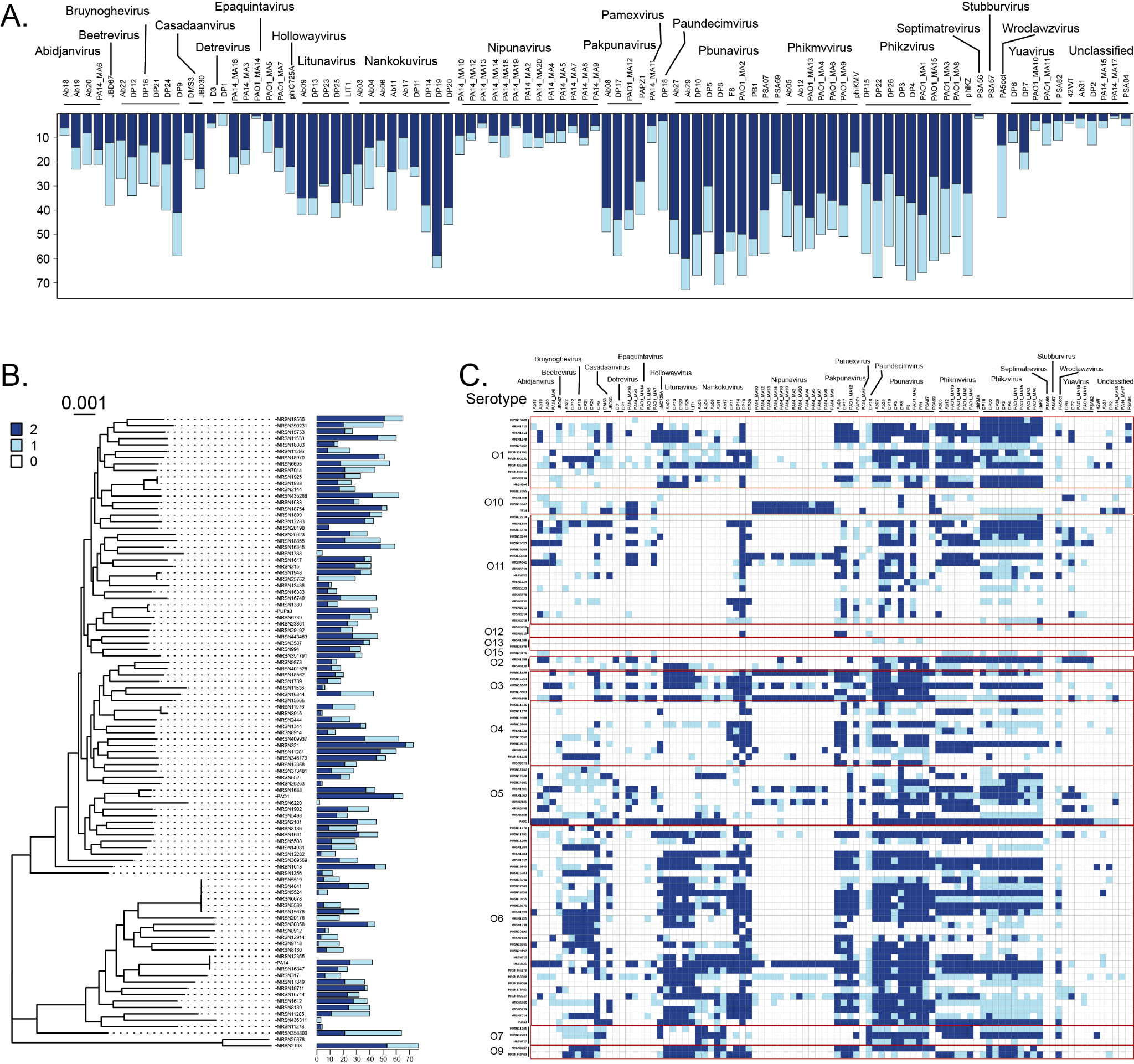

**Supplementary Figure 5. *Pseudomonas* interaction scoring overview.** (A) The number of positive interactions shown by phages were summed and represented as a barplot. Interaction outcomes designated as scores of “1” or “2” were grouped independently. The score of “1” indicates that a hazy clearing of the lawn was observed (light blue), whereas the score of “2” indicates that a complete clearing of lawn or individual plaques were observed (dark blue). The names of phages are listed on the top of the panel. The phages are grouped based on the genera and the names of phages belonging to the same genus are underlined. The genus name is indicated on the top of the panel. (B) The number of positive interactions seen in each strain were added independently and represented as a barplot. Interaction outcomes designated as scores of “1” or “2” were grouped independently. The genomic diversity of the strains are represented by the phylogenetic tree. (C) The strains were grouped together based on the serotype (enclosed in a red box) and the interaction outcome is represented as a heatmap.

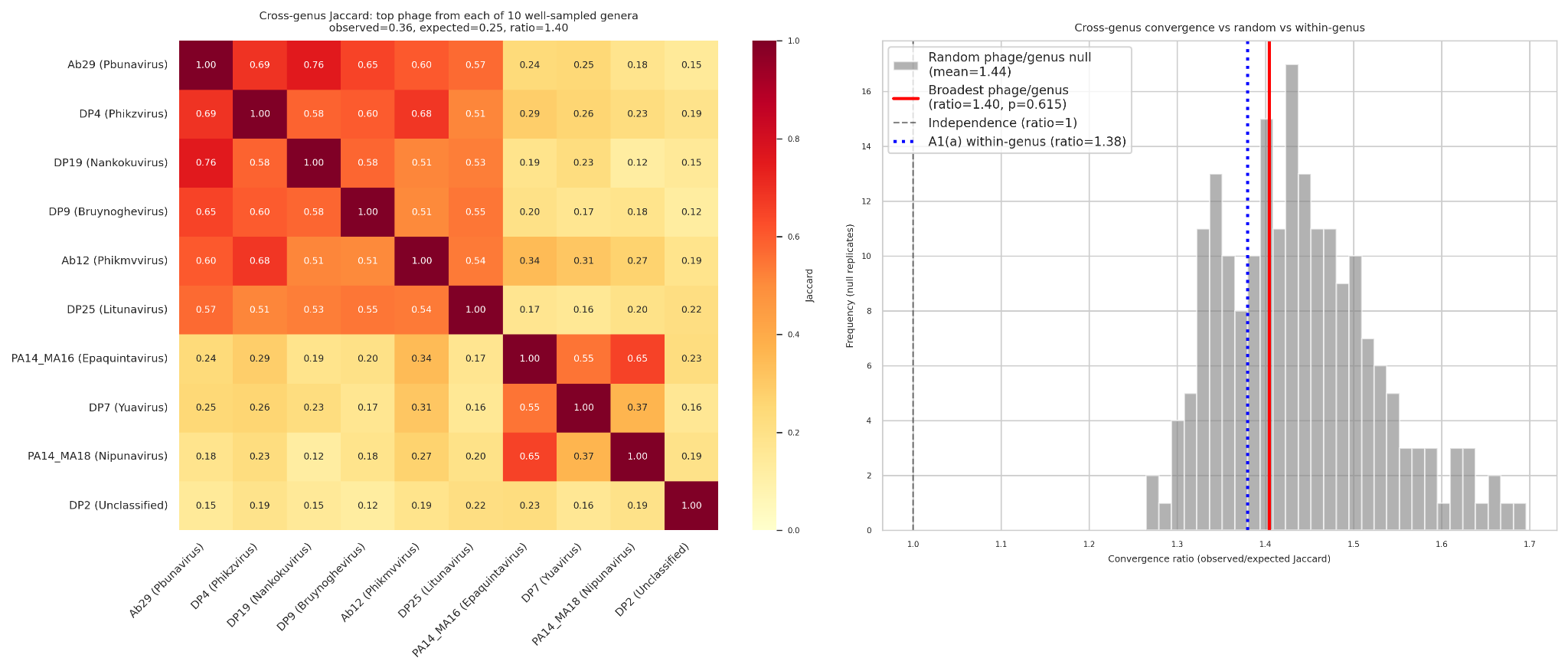

**Supplementary Figure 6. Cross-genus convergence among the broadest-range phages.** Pairwise Jaccard overlap matrix for the broadest-range phage from each of 10 well-sampled phage genera (≥5 phages). Cells show pairwise Jaccard similarity in host range; color scale is Jaccard distance. The mean observed Jaccard across these 45 inter-genus pairs is 0.36, ~40% above the marginal-rate expectation of 0.25 under independent host sampling (convergence ratio 1.40). This cross-genus convergence is comparable to the within-genus convergence among the top-10 broadest phages overall (ratio 1.38), indicating that apparent generalism is not primarily driven by within-genus shared receptor ancestry but reflects a property of broad host range itself.

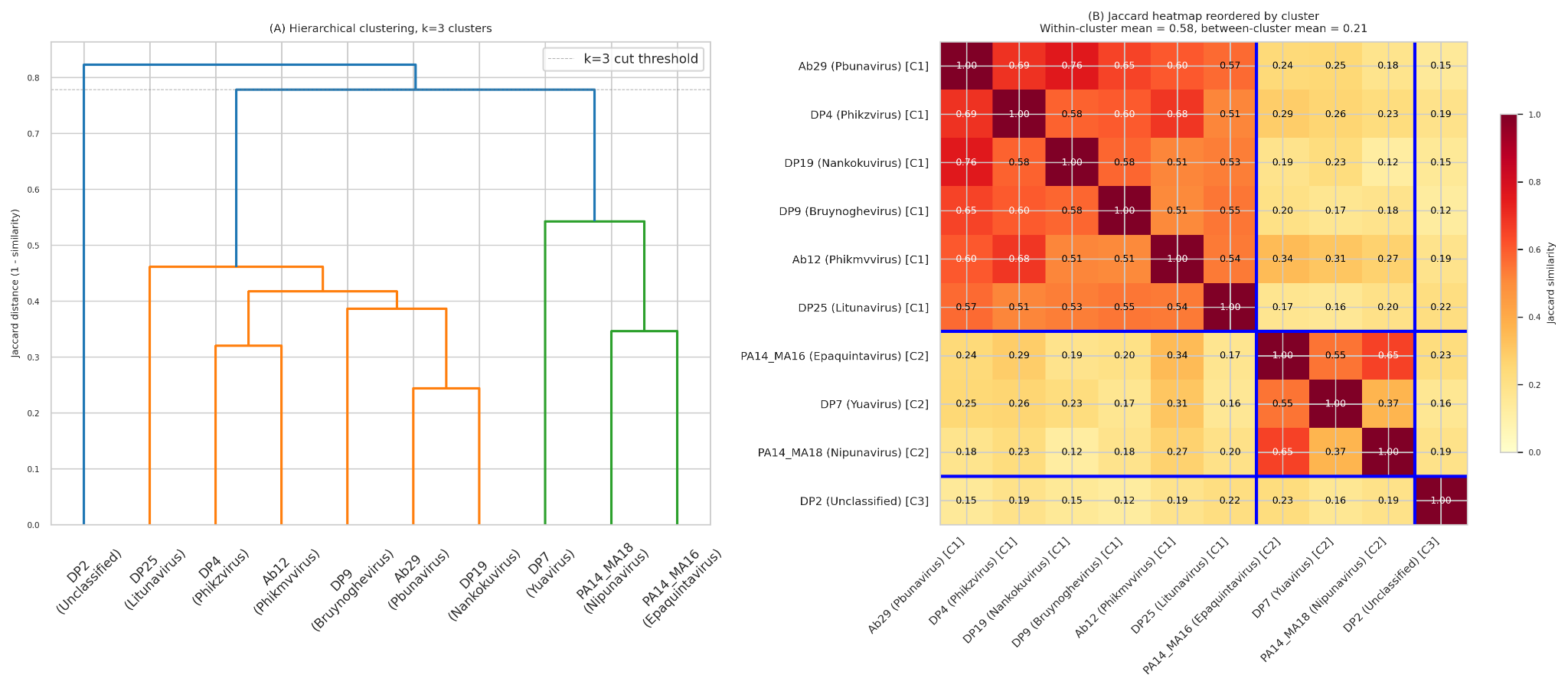

**Supplementary Figure 7. Hierarchical clustering of top-10 broadest phages by host-range overlap.** Dendrogram of the top-10 broadest-range phages clustered by pairwise Jaccard distance in host range. Hierarchical clustering identifies modest within-pool sub-structure but no qualitatively distinct broad-spectrum strategies, consistent with broad-range phages converging on largely overlapping host sets rather than partitioning into receptor-defined coverage tiers.

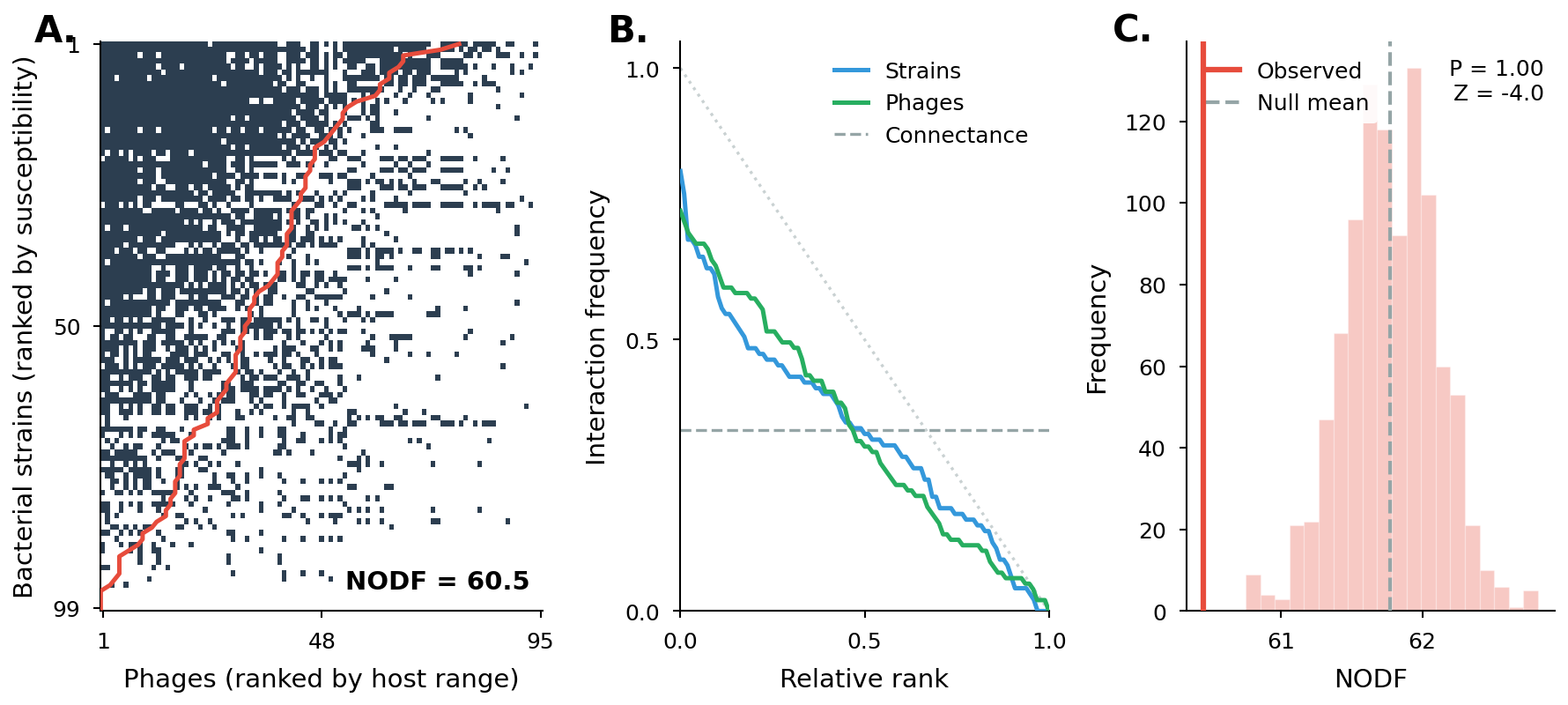

**Supplementary Figure 8**. **Nestedness statistics for the global phage-host interaction matrix.** (A) Binary interaction matrix sorted by decreasing phage host range and decreasing strain susceptibility; dark cells represent positive infections (score ≥1). The red isocline traces the expected boundary under perfect nestedness. The observed NODF of 60.5 indicates a heterogeneous interaction structure with substantial deviations from a hierarchical nested architecture. (B) Rank-frequency curves showing the proportion of interaction partners as a function of relative rank for strains (blue) and phages (green); both curves reflect heterogeneous degree distributions, with the dashed gray line indicating overall matrix connectance (~0.33). (C) Null-model significance test comparing the observed NODF (red line) against 1,000 null matrices generated under a degree-preserving swap null model (fixed row and column marginals); the observed NODF falls entirely below the null distribution (Z = −4.0, one-sided p < 0.001 for anti-nestedness), indicating that the network is significantly anti-nested — broad-range and narrow-range phages target non-overlapping host subsets rather than a hierarchical generalist-includes-specialist architecture. Per-genus NODF tests identified five of 10 well-sampled genera (Pbunavirus, Phikmvvirus, Bruynoghevirus, Epaquintavirus, Litunavirus) as significantly anti-nested after Benjamini-Hochberg correction.

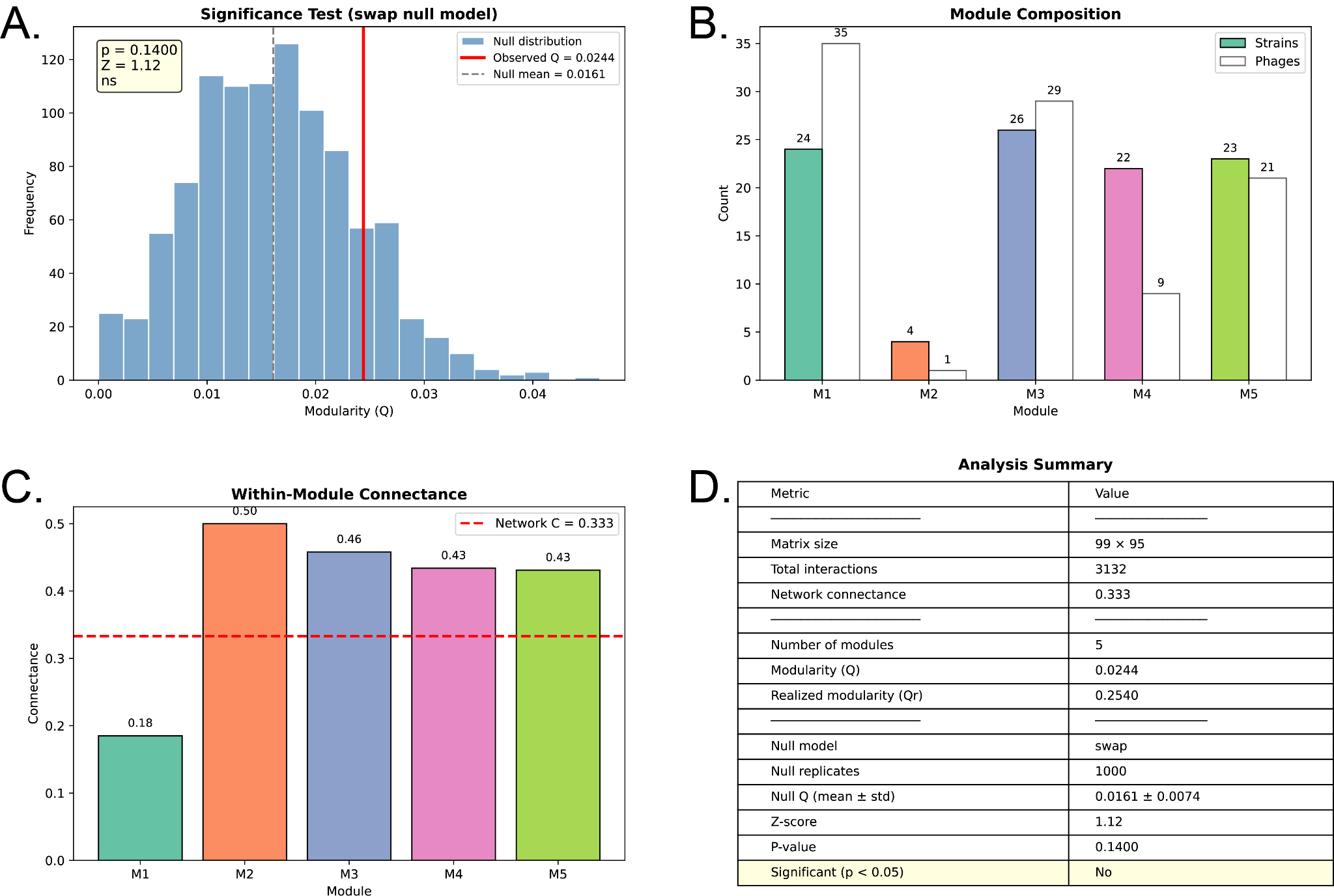

**Supplementary Figure 9. Modularity Statistics.** (A) Null model significance test showing the distribution of modularity (Q) values from 1000 swap-randomized matrices preserving row and column degree sequences; the observed Q of 0.0244 (red line) falls within the null distribution (mean = 0.0161 ± 0.0074), yielding a non-significant result (Z = 1.12, p = 0.14). (B) Module composition showing the number of strains (solid bars) and phages (hatched bars) assigned to each of the five modules, with relatively balanced sizes except for the small Module 2 (4 strains, 1 phage). (C) Within-module connectance for each module compared to the overall network connectance of 0.333 (red dashed line); modules 2–5 show moderately elevated connectance (0.43–0.50) while Module 1 is notably sparse (0.18). (D) Summary table of all key network metrics, modularity values, and null model results.

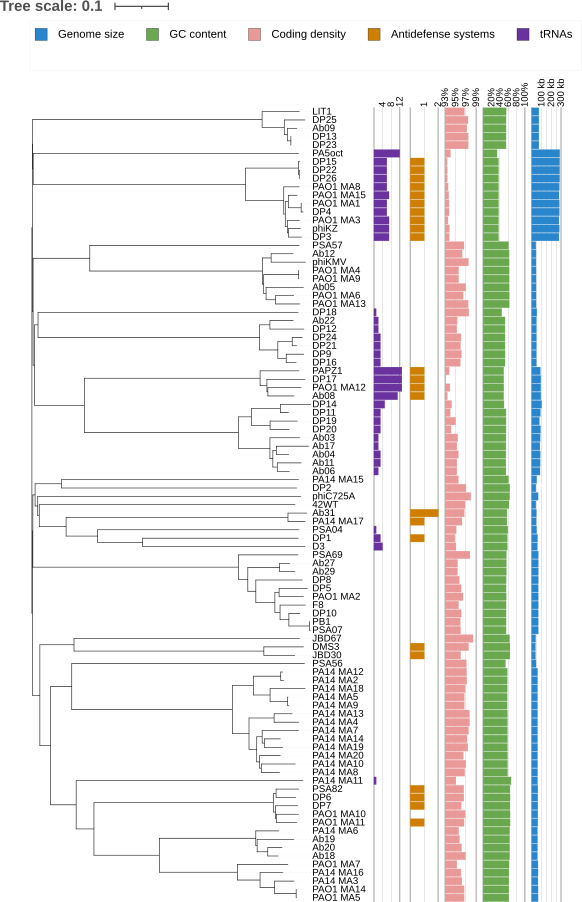

**Supplementary Figure 10. Phage genomic features.** Coding genes and tRNAs were identified in genomes of 153 *P. aeruginosa* phages by Pharokka. Anti-defense systems were identified by DefenseFinder. The dendrogram shows hierarchical clustering of phage genomes based on pairwise PEQ values calculated by PhamClust.

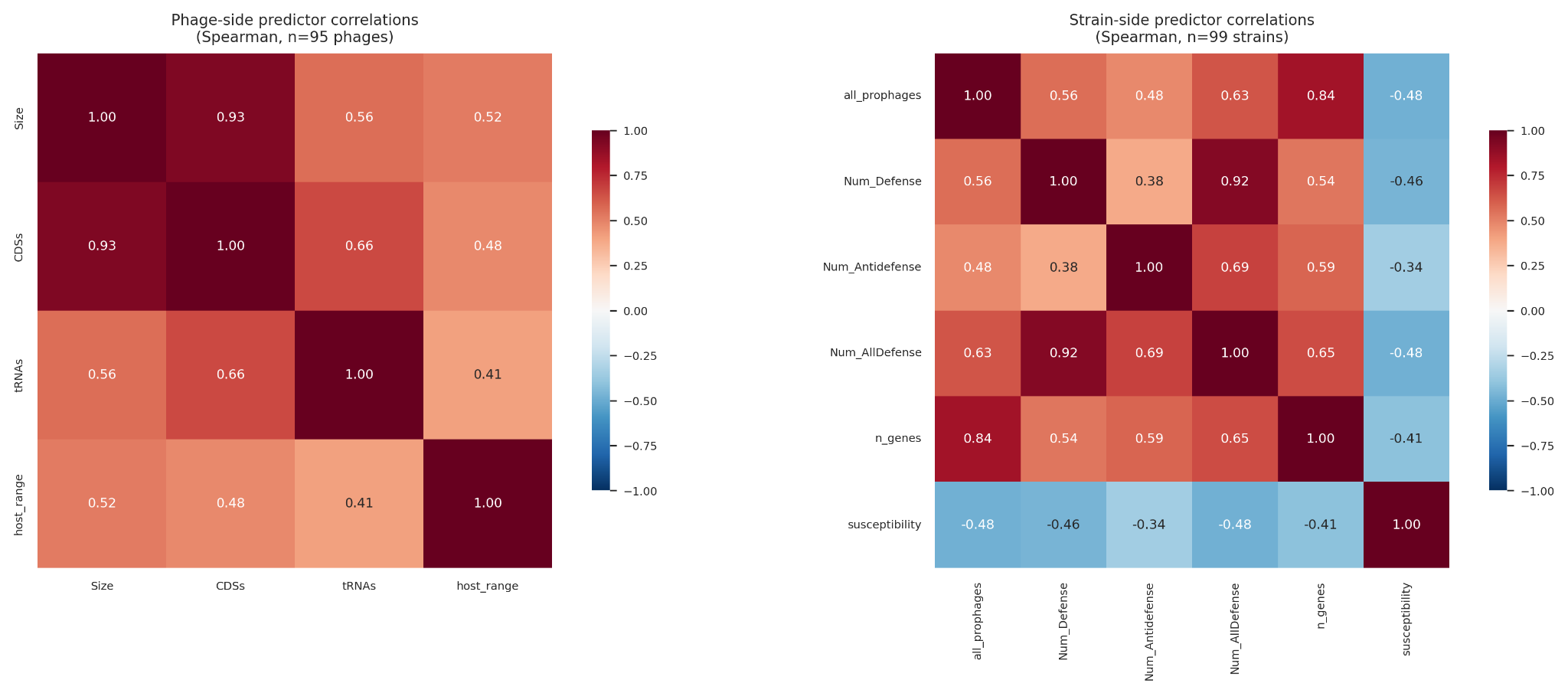

**Supplementary Figure 11. Predictor collinearity.** Spearman correlation matrices for (left) phage-side continuous predictors (genome size, CDS count, tRNA count) used in the Type III ANOVA on phage host range, and (right) strain-side continuous predictors (defense system count, anti-defense system count, prophage burden, total gene count) used in the Type III ANOVA on strain susceptibility. Cell color and number give Spearman ρ. Genome size and CDS count are nearly redundant on the phage side (ρ = 0.93); prophage burden correlates with anti-defense count (ρ = 0.48) and total gene count (ρ = 0.84) on the strain side. Variance inflation factors for all design matrices were below 5 (Table S-MV06, S-MV06b), confirming that residual multicollinearity is moderate and that unique-variance contributions can be interpreted at face value.

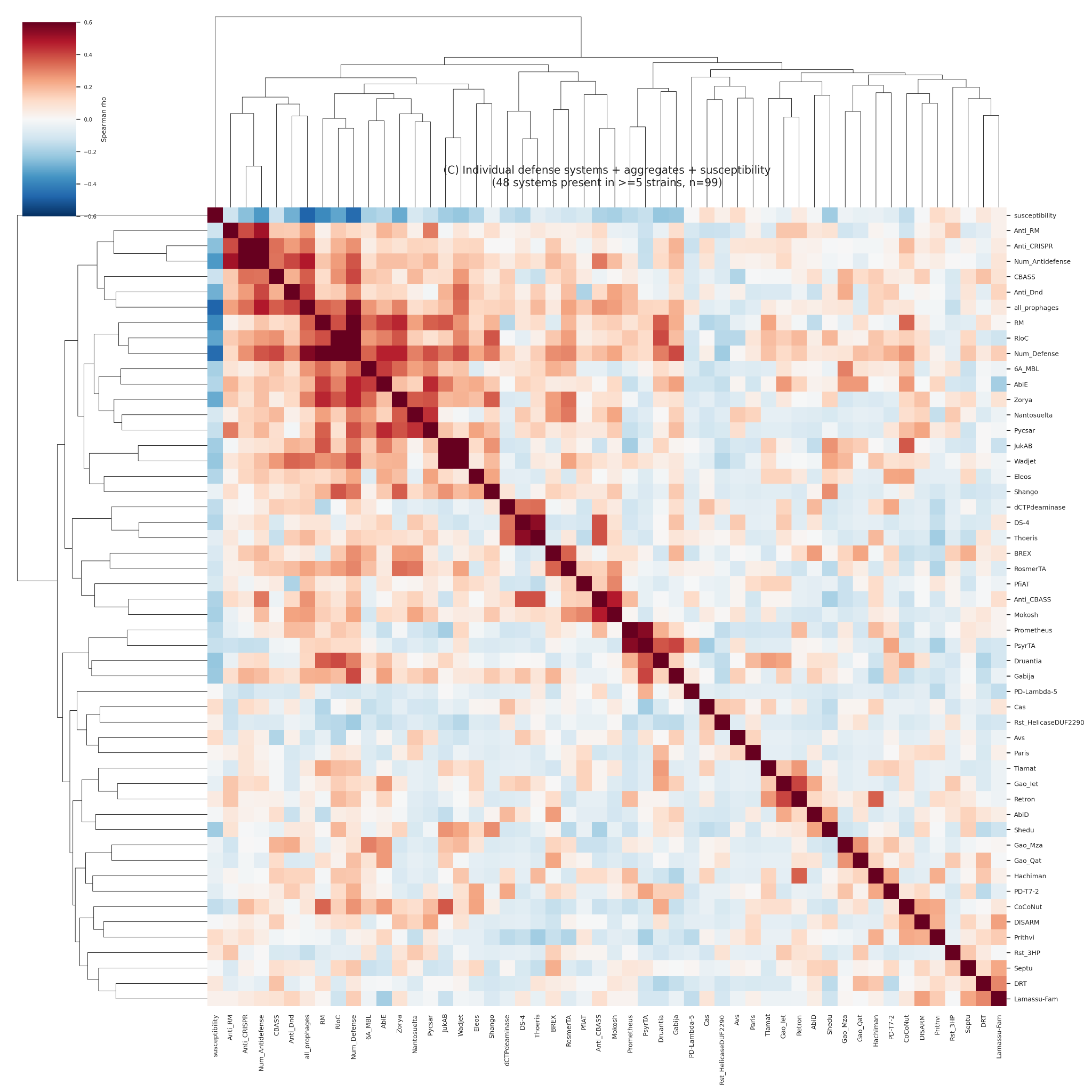

**Supplementary Figure 12. Pairwise correlation structure among individual defense systems and overall susceptibility.** Clustered heatmap of pairwise Spearman correlations among the 48 defense-system families present in ≥5 strains, with overall strain susceptibility, total defense-system count, total anti-defense count, and prophage count added as additional rows/columns. Cell values are per-strain copy counts of each defense-system family (range 0–13). Cells are colored by Spearman ρ (red = positive, blue = negative); hierarchical clustering on both axes groups defense systems with correlated abundance patterns across the 99-strain panel. No individual defense system shows a strong correlation with susceptibility

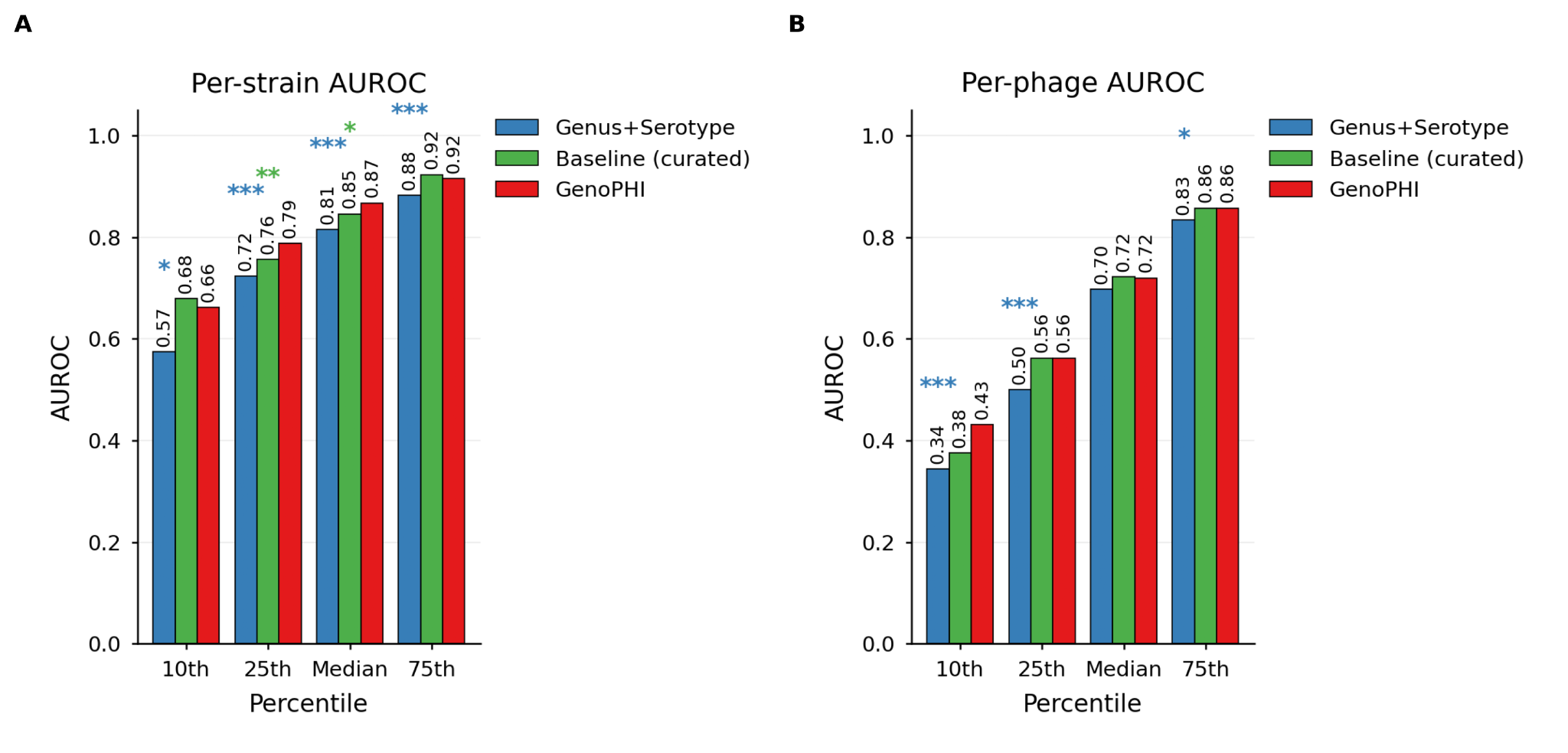

**Supplementary Figure 13. Performance across percentiles of per-entity AUROC.** Per-entity AUROC at the 10th, 25th, 50th (median), and 75th percentiles across held-out entities for the three deployable models. (A) Per-strain AUROC percentiles. (B) Per-phage AUROC percentiles. Asterisks indicate paired bootstrap significance vs GenoPHI at each percentile (*p < 0.05, **p < 0.01, ***p < 0.001; 2000 bootstrap resamples of paired entity-level AUROCs, two-sided). The figure highlights model behavior at the tail of difficulty: GenoPHI raises the 10th-percentile per-phage AUROC over Genus+Serotype (0.43 vs 0.34, p < 0.001) and matches the curated baseline at the 25th percentile. Per-strain floor performance follows the same direction but with smaller absolute differences across models.

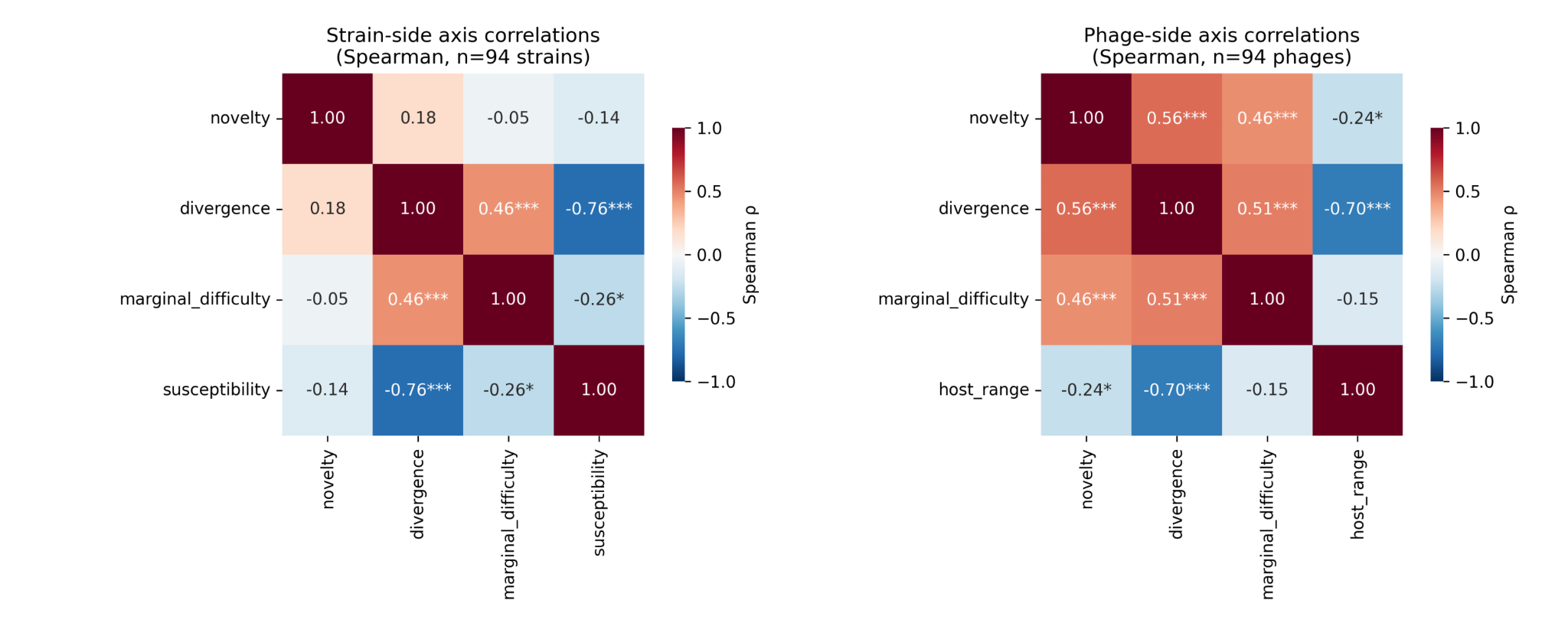

**Supplementary Figure 14. Difficulty-axis Spearman correlations.** Pairwise Spearman correlations between the four difficulty axes used for stratified performance analysis, computed across entities. (Left) Strain-side axis correlations across n=94 strains: phylogenetic novelty (patristic distance to nearest training strain), phenotypic divergence (Jaccard distance to five phylogenetic nearest neighbors in interaction profile), marginal difficulty (1 − marginal-model AUROC), and susceptibility (per-strain interaction count). (Right) Phage-side axis correlations across n=94 phages, with host range replacing susceptibility on the diagonal. Cells annotated with Spearman ρ and significance (*p < 0.05, **p < 0.01, ***p < 0.001). Strong correlations among divergence/marginal-difficulty and between divergence/susceptibility (or host-range) confirm that the axes capture partially overlapping signals but remain non-redundant enough to justify stratification along each individually; phylogenetic novelty is the most independent of the four on the strain side.

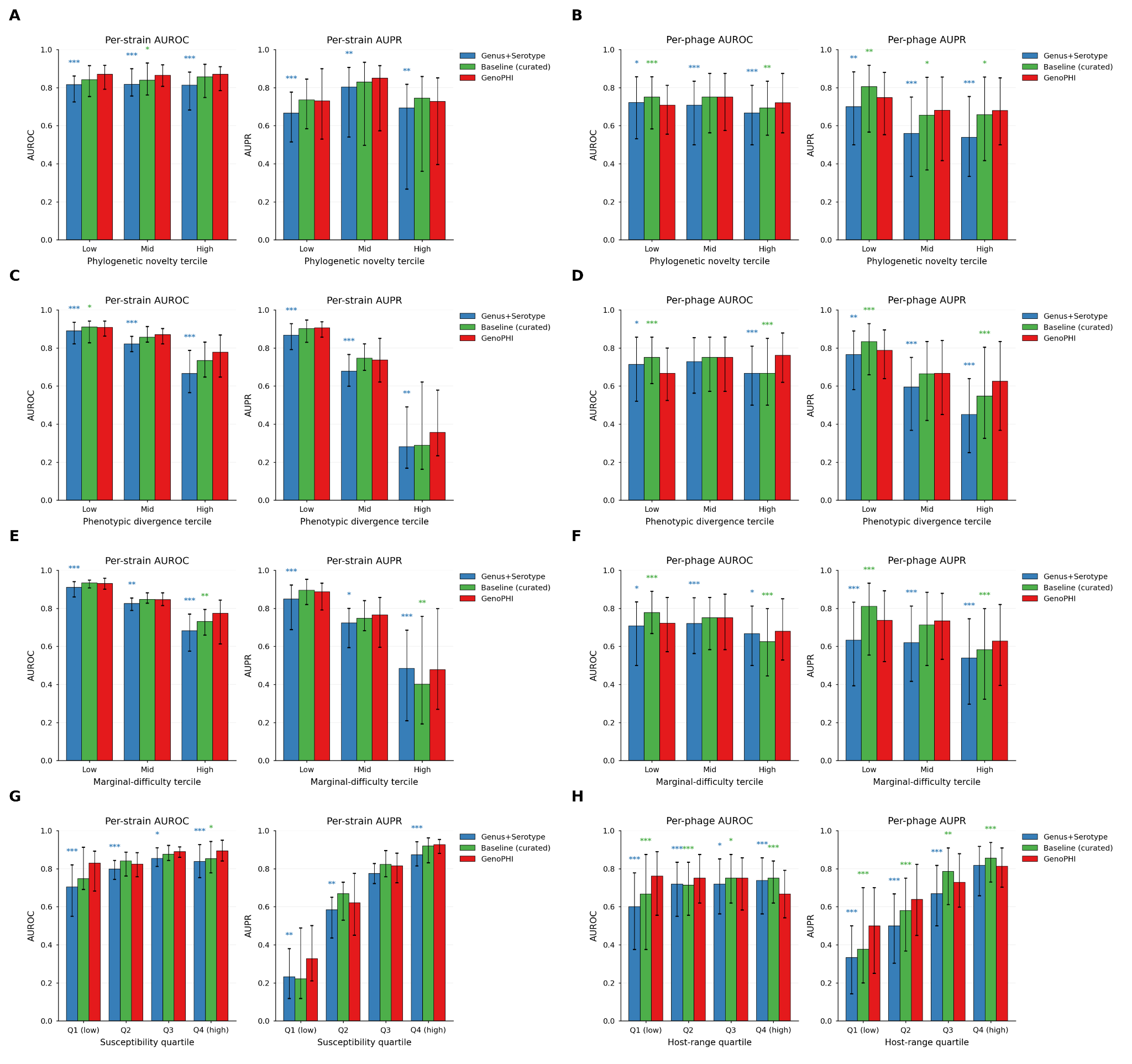

**Supplementary Figure 15. Per-entity model performance stratified by four difficulty axes.** Cross-validated per-strain (left panels of each row pair) and per-phage (right panels of each row pair) AUROC and AUPR for the three deployable CatBoost models — Genus+Serotype (blue), Baseline (curated, green), and GenoPHI (red) — across four difficulty axes. (A) Per-strain AUROC and AUPR by phylogenetic novelty tercile (patristic distance to nearest training strain). (B) Per-phage AUROC and AUPR by phylogenetic novelty tercile (patristic distance to nearest neighboring phage in PEQ-based tree). (C) Per-strain AUROC and AUPR by phenotypic divergence tercile (Jaccard distance to five phylogenetic nearest neighbors in interaction profile). (D) Per-phage AUROC and AUPR by phenotypic divergence tercile. (E) Per-strain AUROC and AUPR by marginal-difficulty tercile (1 − marginal-model AUROC; captures failures of degree-only prediction). (F) Per-phage AUROC and AUPR by marginal-difficulty tercile. (G) Per-strain AUROC and AUPR by susceptibility quartile (full-matrix per-strain interaction count). (H) Per-phage AUROC and AUPR by host-range quartile. Bars show median across entities within each bin; error bars span the interquartile range. Asterisks indicate paired Wilcoxon significance vs GenoPHI within each bin (*p < 0.05, **p < 0.01, ***p < 0.001). Both feature-rich models substantially outperformed Genus+Serotype on the hardest bins of each axis; GenoPHI specifically outperformed the curated baseline on per-phage AUROC and AUPR in the highest-difficulty terciles, while the curated baseline matched or modestly exceeded GenoPHI in the lowest-difficulty terciles.

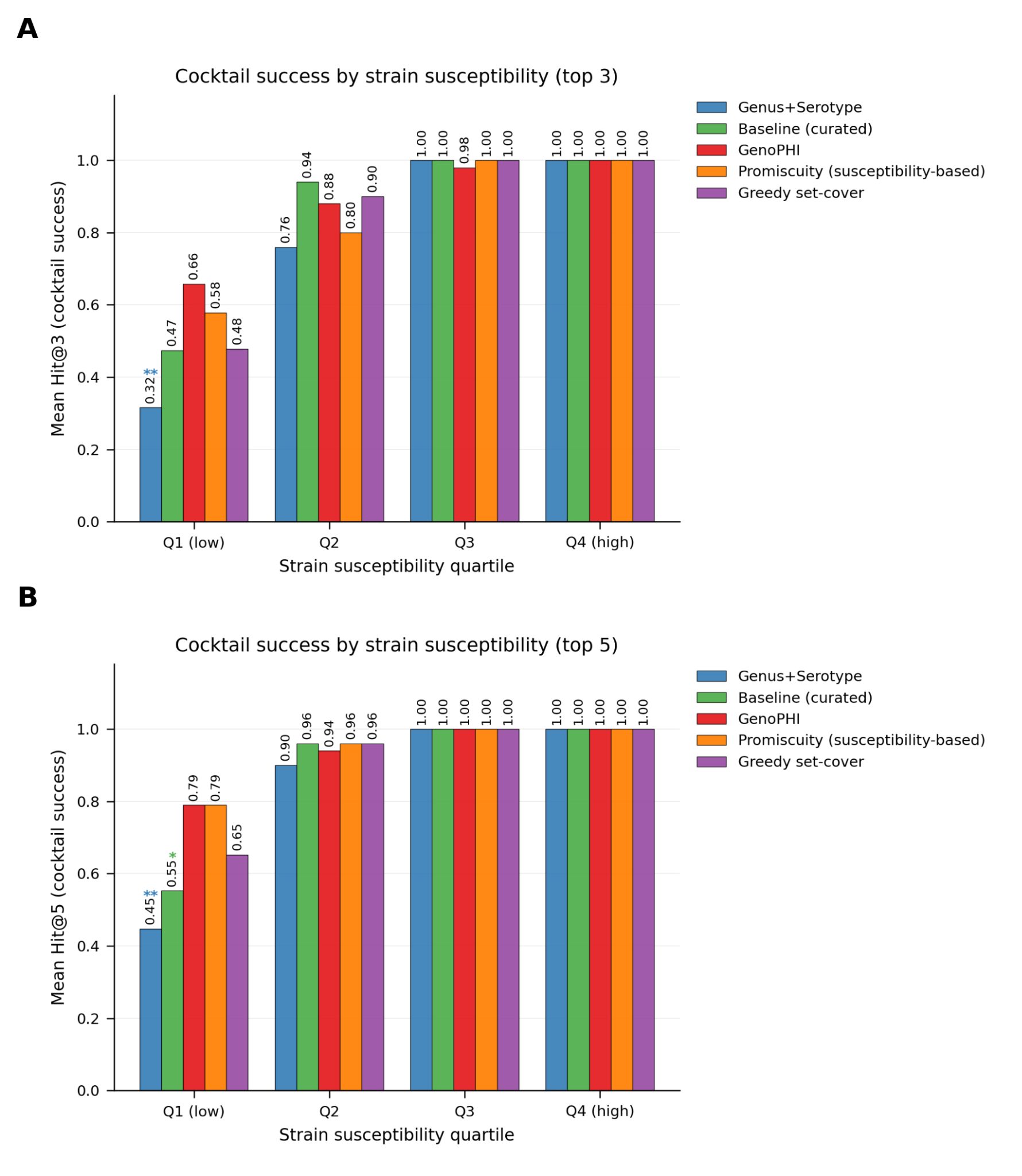

**Supplementary Figure 16. Cocktail-design performance by strain susceptibility quartile.** Cocktail success rate, defined as the fraction of held-out strains for which the recommended top-k phages contain at least one true infectious match, stratified by strain susceptibility quartile (Q1 lowest, Q4 highest). Five strategies are compared: three deployable CatBoost models: (1) Genus+Serotype (blue), (2) Baseline (curated genomic features, green), and (3) GenoPHI (pangenome-derived gene-cluster features, red); and two non-ML baselines: (1) a promiscuity-based method (orange) that hierarchically clusters phages on the training-fold interaction matrix and selects the most-infectious phage per cluster, and (2) a greedy set-cover that maximizes training-fold strain coverage (purple). For the ML models, top-k phages were selected per held-out strain by predicted P(infection), with HDBSCAN-derived phage clustering enforcing receptor diversity (GenoPHI clustered on its per-fold selected pangenome features; Genus+Serotype and the curated baseline clustered on the training-fold interaction matrix). (A) Top-3 cocktails. (B) Top-5 cocktails. Bars show the mean cocktail success rate across all held-out strains in each quartile. GenoPHI achieved the highest top-3 cocktail success on the lowest-susceptibility quartile (Q1: 65.8%) versus the susceptibility-based promiscuity baseline (57.9%), greedy set-cover (47.8%), the curated baseline (47.4%), and Genus+Serotype (31.6%; McNemar p = 0.002 vs GenoPHI). At top-5 GenoPHI tied the promiscuity baseline at 78.9% on Q1. On Q3 and Q4 strains all approaches converged to ≥98% top-3 cocktail success, indicating that cocktail design reduces to a covering problem on permissive hosts and that individual phage selection matters most for the lowest-susceptibility quartile.

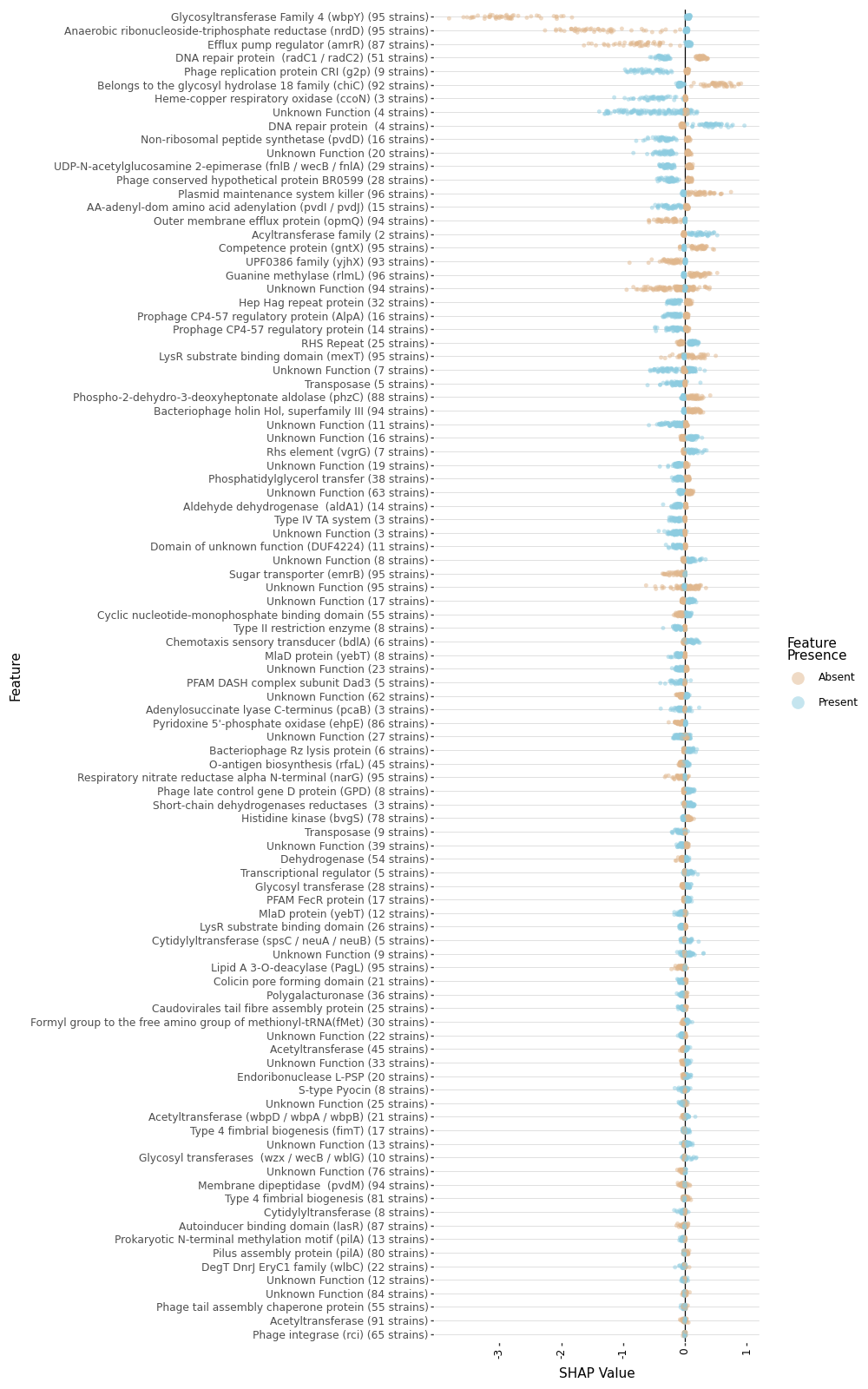

**Supplementary Figure 17. Predictive feature annotations and SHAP importances.** SHAP values associated with predictive features indicate whether presence of a feature (blue) or absence (orange) is associated with increased likelihood of infection (SHAP value > 0) or decreased likelihood of infection (SHAP value < 0).

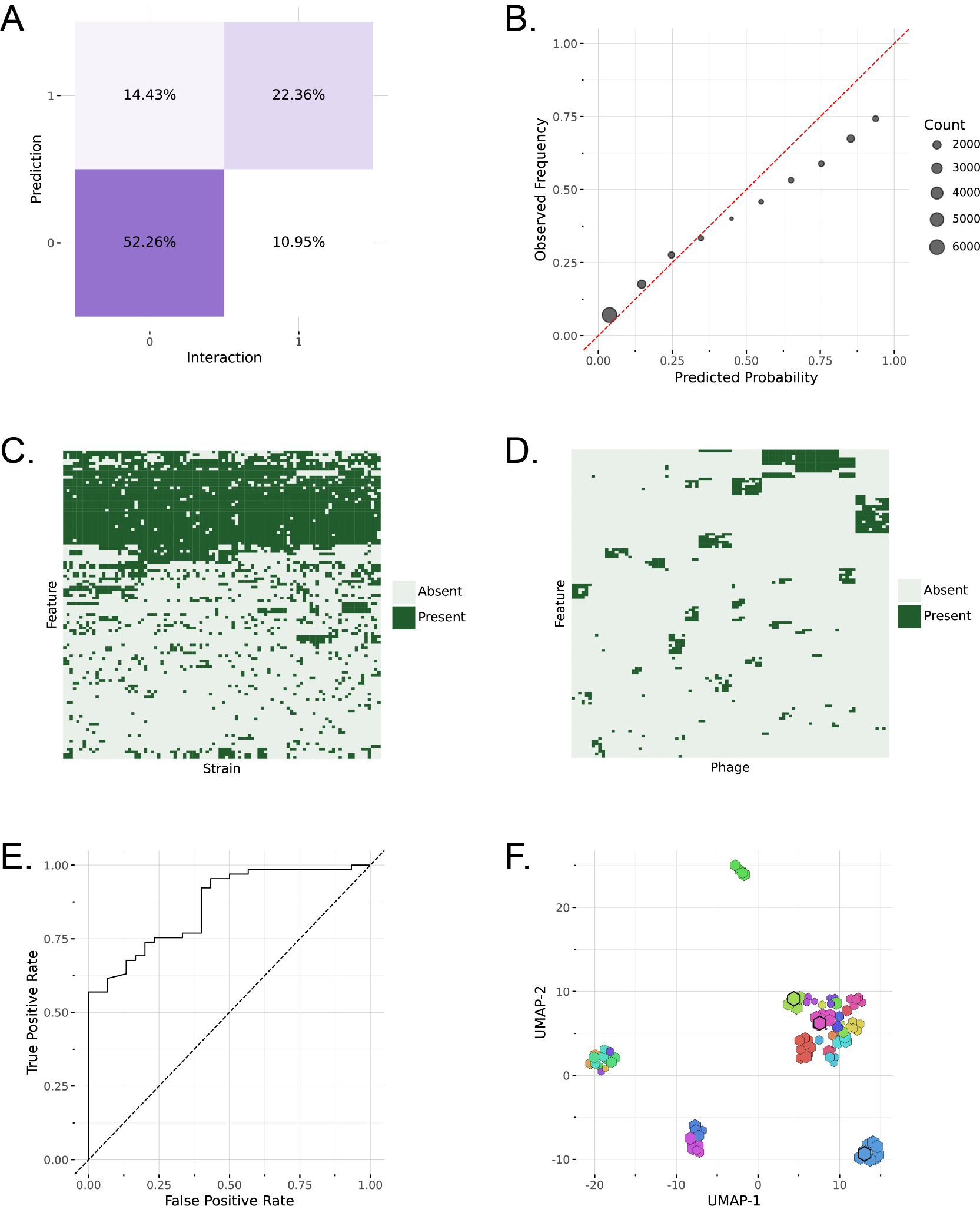

**Supplementary Figure 18. Overview of GenoPHI models.** (A) The confusion matrix shows the proportions of true-positives, true-negatives, false-positives, and false-negatives across 20-fold cross-validation experiments. (B) The calibration curve compares predicted vs. observed infection rates, showing a slight over prediction. (C) The distribution of features across strains shows limited clustering and a mixture of highly conserved and sparsely conserved features. (D) The distribution of features across phages shows tight clustering and limited sharing of features across phages. (E) The receiver operating characteristic (ROC) for model predictions on PAO1. (F) The UMAP projection shows the distribution of phages sized by their predicted likelihood of activity on PAO1. Colors represent HDBSCAN phage clusters and selected phages are highlighted in black.

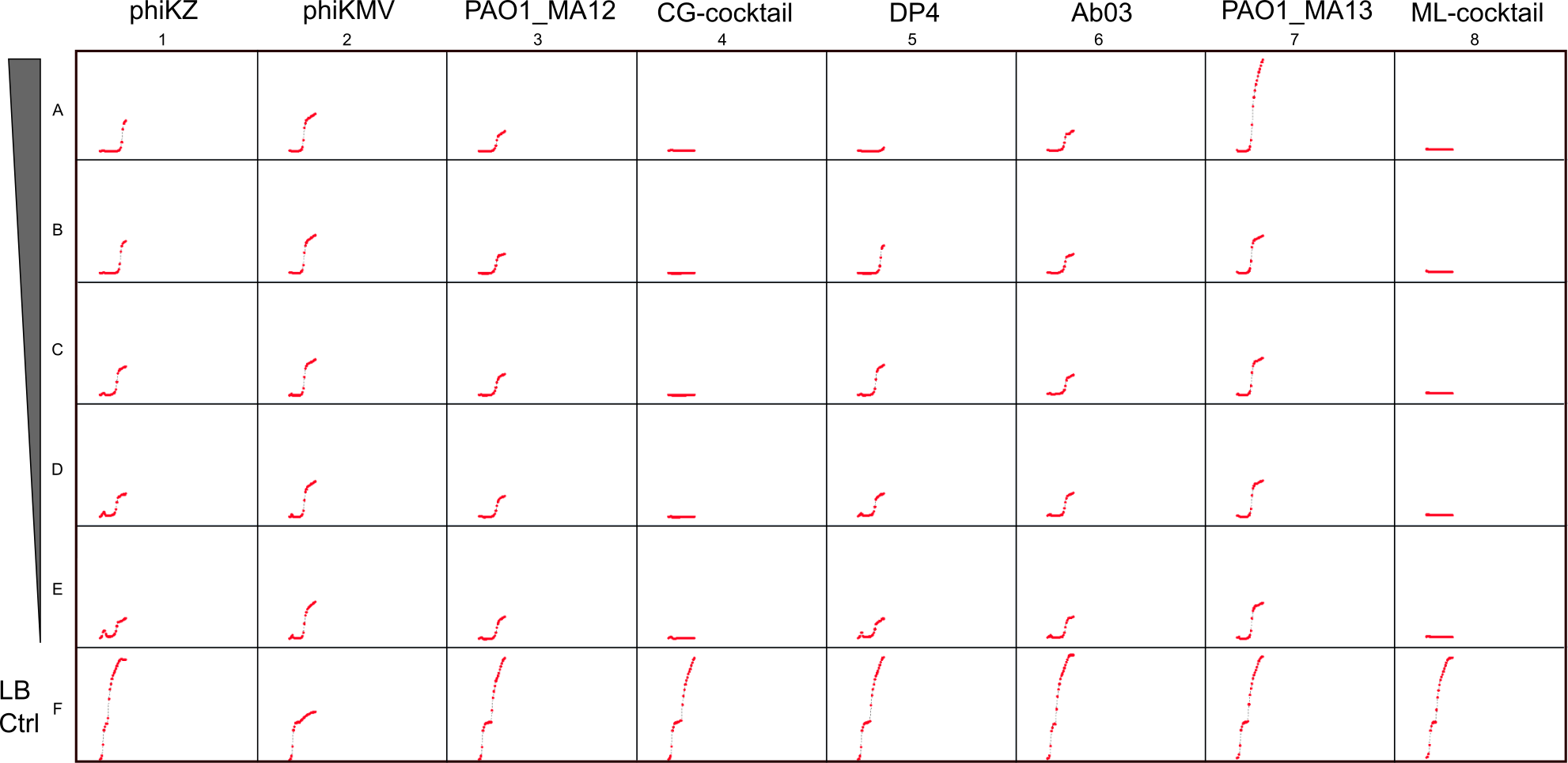

**Supplementary Figure 19. Inhibition of PAO1 growth by phage cocktails.** The titer of all individual phages were normalized to 1 x 10^9^ PFU/ml. To formulate the cocktails, individual phages were normalized to the titer of 3 x 10^9^ PFU/ml and equal volumes of phages were mixed together so that the titer of individual phages in the cocktail was 1 x 10^9^ PFU/ml. The individual phages and formulated cocktails were aliquoted into a 48w plate and were serially diluted ten-fold. The overnight culture of *P. aeruginosa* PAO1 was diluted 100-fold in fresh LB supplemented with 10 mM CaCl2 and MgSO4 and 350 ul was aliquoted into a 48w flat bottom plate. 350 ul of the stock and serially diluted phages were added to the culture. LB was added to all wells in row F. The plate was transferred into a plate reader at 37 ℃ and the killing activity of phages was monitored overnight. The data show that both CG- and ML-cocktails were equally effective in preventing bacterial growth *in vitro* at all dilutions. The plate was shaken linearly at ~570 cpm during the entire assay, except while monitoring the OD.

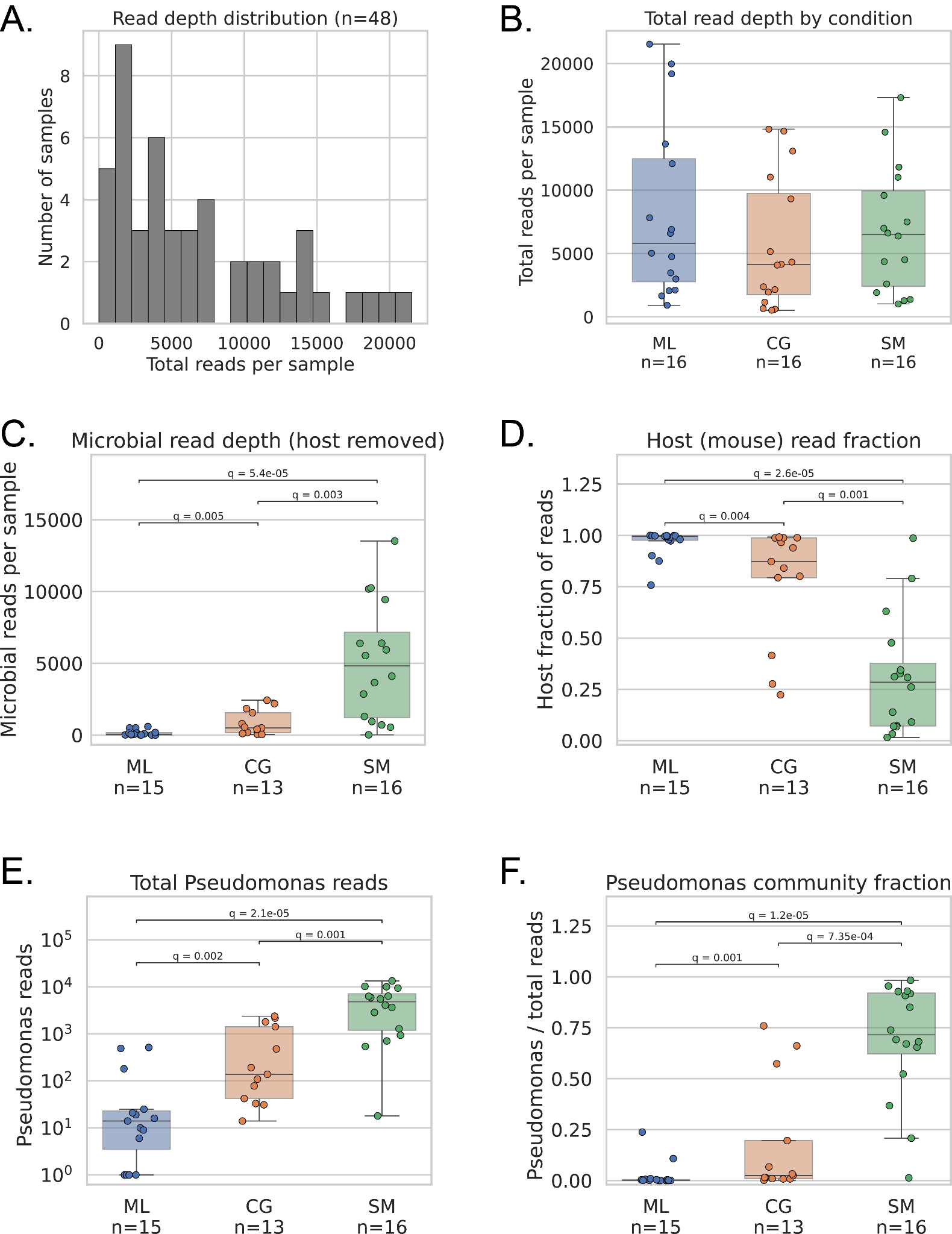

**Supplementary Figure 20. 16S rRNA amplicon sequencing of mouse wound samples across treatment conditions (ML = M-mix, CG = C-mix, SM = SM buffer control).** (A) Distribution of total read depth across all sequenced samples (n = 48 biological samples, sequencing replicates summed per sample). (B) Total read depth by condition. Read depth was comparable across conditions (Kruskal–Wallis P = 0.72). (C) Microbial read depth per sample, with reads assigned to the mouse host ASV removed. (D) Host (mouse) fraction of reads per sample. (E) Total *Pseudomonas* reads per sample (log scale). (F) Pseudomonas fraction of the microbial community, calculated as *Pseudomonas* reads divided by total reads per sample. Panels C–F show samples passing a minimum read-depth QC filter (≥1,000 total reads; n = 44: ML = 15, CG = 13, SM = 16). Boxes show the interquartile range with the median; whiskers extend to 1.5×IQR. Significance brackets indicate pairwise Mann–Whitney U tests with Benjamini–Hochberg FDR correction (q-values shown); non-significant comparisons (q ≥ 0.05) are not annotated.

### Supplementary Tables

| Rank | Genera | No. of phages | No. of positive interaction | Standard deviation |
| --- | --- | --- | --- | --- |
| 1 | Phikzvirus | 10 | 62 | 6 |
| 2 | Pbunavirus | 10 | 59 | 13 |
| 3 | Pakpunavirus | 4 | 50 | 7 |
| 4 | Phikmvvirus | 7 | 48 | 12 |
| 5 | Litunavirus | 5 | 39 | 5 |
| 6 | Nankokuvirus | 9 | 38 | 14 |
| 7 | Bruynoghevirus | 6 | 37 | 12 |
| 8 | Casadabanvirus | 2 | 25 | 8 |
| 9 | Abidjanvirus | 4 | 19 | 6 |
| 10 | Epaquintavirus | 5 | 18 | 9 |
| 11 | Yuavirus | 5 | 13 | 6 |
| 12 | Nipunavirus | 13 | 12 | 4 |
| 13 | Detrevirus | 2 | 6 | 1 |

**Table S1. Number of positive interactions per phage from each genus.** For genera that had more than one candidate phages in our collection, the number positive interactions were averaged. The genera are arranged based on the number of positive interactions. Phages from Phikzvirus genus show positive interaction with the highest number of hosts (62), whereas phages from Detrevirus show the lowest number of positive interactions (6).

| **Genus** | **N phages** | **N strains with any** | **NODF** | **null_mean** | **null_std** | **z_score** | **p_nested** | **p_antinested** | **direction** |
| --- | --- | --- | --- | --- | --- | --- | --- | --- | --- |
| Nipunavirus | 13 | 23 | 79.9 | 80 | 0.807 | -0.0595 | 0.57 | 0.43 | anti-nested |
| Pbunavirus | 10 | 83 | 77 | 79.3 | 0.298 | -7.62 | 1 | 0 | anti-nested |
| Phikzvirus | 10 | 81 | 67.7 | 67.6 | 0.217 | 0.249 | 0.455 | 0.545 | nested |
| Nankokuvirus | 9 | 83 | 68.4 | 68.4 | 0.576 | -0.0298 | 0.59 | 0.41 | anti-nested |
| Phikmvvirus | 7 | 65 | 73.2 | 75.7 | 0.298 | -8.6 | 1 | 0 | anti-nested |
| Unclassified | 6 | 25 | 24.7 | 23.9 | 1.4 | 0.558 | 0.335 | 0.665 | nested |
| Bruynoghevirus | 6 | 74 | 61.9 | 65.3 | 0.485 | -6.88 | 1 | 0 | anti-nested |
| Yuavirus | 5 | 29 | 55.5 | 57.5 | 1.21 | -1.63 | 0.935 | 0.065 | anti-nested |
| Epaquintavirus | 5 | 37 | 54.6 | 62.4 | 0.624 | -12.6 | 1 | 0 | anti-nested |
| Litunavirus | 5 | 46 | 58.6 | 59.3 | 0.23 | -2.87 | 0.965 | 0.035 | anti-nested |

**Table S2.** Per-genus anti-nestedness. NODF computed within each of the 10 well-sampled genera (≥5 phages), with degree-preserving null mean, Z-score, and one-sided p-values for nestedness and anti-nestedness. Five genera (Pbunavirus, Phikmvvirus, Bruynoghevirus, Epaquintavirus, Litunavirus) are significantly anti-nested after Benjamini-Hochberg correction.

| **Serotype** | **No of strains** | **No_1** | **No_2** | **Total** | **Average** | **SD** |
| --- | --- | --- | --- | --- | --- | --- |
| O3 | 5 | 14 | 46 | 60 | 49 | 26 |
| O6 | 31 | 1 | 3 | 4 | 40 | 17 |
| O2 | 2 | 7 | 37 | 44 | 37 | 10 |
| O1 | 11 | 2 | 9 | 11 | 36 | 17 |
| O5 | 9 | 11 | 3 | 14 | 36 | 15 |
| O7 | 3 | 29 | 11 | 40 | 34 | 14 |
| O11 | 17 | 13 | 3 | 16 | 22 | 13 |
| O4 | 10 | 2 | 4 | 6 | 21 | 13 |
| O9 | 2 | 5 | 35 | 40 | 43 | 4 |
| O10 | 4 | 0 | 0 | 0 | 19 | 18 |
| O15 | 1 | 17 | 0 | 17 | 17 | 0 |
| O12 | 2 | 2 | 0 | 2 | 3 | 1 |
| O13 | 3 | 4 | 0 | 4 | 2 | 3 |

**Table S3. Phage infection outcome grouped by host O-antigen serotype.** *P. aeruginosa* strains are grouped by serotype; for each serotype, score-1 (turbid) and score-2 (clear) interactions are summed across member strains, and the mean ± SD phages per strain is reported. O6 is most represented (31 strains); O15 has a single representative. Strains of serotype O3 are infected by the most phages on average, whereas O12 and O13 strains are infected by the fewest.

| **predictor** | **VIF** | **flag** |
| --- | --- | --- |
| n_genes | 3.48 | ok |
| all_prophages | 2.91 | ok |
| sero_O6 | 2.77 | ok |
| sero_O11 | 2.26 | ok |
| sero_O4 | 1.9 | ok |
| sero_O5 | 1.77 | ok |
| Num_Defense | 1.62 | ok |
| sero_O3 | 1.43 | ok |
| Num_Antidefense | 1.36 | ok |
| sero_O10 | 1.35 | ok |
| sero_O12 | 1.33 | ok |
| sero_O2 | 1.29 | ok |
| sero_O7 | 1.25 | ok |
| sero_O13 | 1.23 | ok |
| sero_O15 | 1.21 | ok |
| sero_O9 | 1.16 | ok |

**Table S4.** Variance inflation factors for the Type III ANOVA predictor matrix. All VIFs < 5 (max 3.5), indicating moderate, interpretable collinearity.

| **predictor** | **GVIF** | **Df** | **GVIF^(1/(2*Df))** |
| --- | --- | --- | --- |
| Defense | 1.59 | 1 | 1.26 |
| Antidefense | 1.26 | 1 | 1.12 |
| all_prophages | 1.47 | 1 | 1.21 |
| serotype | 1.79 | 12 | 1.02 |

**Table S5.** Generalized variance inflation factors for the PERMANOVA predictor matrix. All GVIFs < 5 (max 1.79).

| **comparison** | **controlling_for** | **rho_raw** | **p_raw** | **rho_partial** | **p_partial** | **shrinkage** |
| --- | --- | --- | --- | --- | --- | --- |
| Size vs. host_range | Genus | 0.519 | 7.2e-08 | 0.111 | 0.282 | 0.407 |
| CDSs vs. host_range | Genus | 0.482 | 7.4e-07 | 0.0862 | 0.406 | 0.396 |
| tRNAs vs. host_range | Genus | 0.41 | 3.6e-05 | 0.174 | 0.0922 | 0.237 |
| all_prophages vs. susceptibility | Num_Antidefense + Num_Defense | -0.48 | 5.0e-07 | -0.252 | 0.0128 | -0.228 |
| Num_Antidefense vs. susceptibility | all_prophages | -0.344 | 4.9e-04 | -0.148 | 0.147 | -0.196 |
| Num_Defense vs. susceptibility | all_prophages+Num_Antidefense | -0.462 | 1.5e-06 | -0.25 | 0.0135 | -0.212 |
| Num_Defense vs. susceptibility | serotype | -0.462 | 1.5e-06 | -0.343 | 5.2e-04 | -0.119 |

**Table S6.** Univariate versus controlled partial Spearman correlations. Top three rows: continuous phage features (genome size, CDS count, tRNA count) versus host range, controlled for genus. Remaining rows: strain-level features versus susceptibility, each controlled for the indicated covariates. Raw correlations attenuate substantially after partialling out the controlling variable.

| **Genus** | **feature** | **n_phages** | **rho** | **p_raw** | **reason_skipped** | **p BH per feature** | **P BH pooled** |
| --- | --- | --- | --- | --- | --- | --- | --- |
| Nipunavirus | Size | 13 | -0.111 | 0.719 |  | 0.985 | 0.985 |
| Nipunavirus | CDSs | 13 | -0.0648 | 0.833 |  | 0.926 | 0.985 |
| Nipunavirus | tRNAs | 13 |  |  | feature constant within genus | | |
| Pbunavirus | Size | 10 | 0.384 | 0.273 |  | 0.985 | 0.985 |
| Pbunavirus | CDSs | 10 | -0.54 | 0.107 |  | 0.371 | 0.668 |
| Pbunavirus | tRNAs | 10 |  |  | feature constant within genus | | |
| Phikzvirus | Size | 10 | -0.0426 | 0.907 |  | 0.985 | 0.985 |
| Phikzvirus | CDSs | 10 | 0.687 | 0.0281 |  | 0.281 | 0.479 |
| Phikzvirus | tRNAs | 10 | 0.0356 | 0.922 |  | 0.922 | 0.985 |
| Nankokuvirus | Size | 9 | 0.217 | 0.576 |  | 0.985 | 0.985 |
| Nankokuvirus | CDSs | 9 | 0.233 | 0.546 |  | 0.92 | 0.985 |
| Nankokuvirus | tRNAs | 9 | 0.689 | 0.0399 |  | 0.16 | 0.479 |
| Phikmvvirus | Size | 7 | 0.00909 | 0.985 |  | 0.985 | 0.985 |
| Phikmvvirus | CDSs | 7 | 0.138 | 0.769 |  | 0.926 | 0.985 |
| Phikmvvirus | tRNAs | 7 |  |  | feature constant within genus | | |
| Unclassified | Size | 6 | -0.029 | 0.957 |  | 0.985 | 0.985 |
| Unclassified | CDSs | 6 | -0.116 | 0.827 |  | 0.926 | 0.985 |
| Unclassified | tRNAs | 6 | -0.133 | 0.802 |  | 0.922 | 0.985 |
| Bruynoghevirus | Size | 6 | 0.0857 | 0.872 |  | 0.985 | 0.985 |
| Bruynoghevirus | CDSs | 6 | -0.309 | 0.552 |  | 0.92 | 0.985 |
| Bruynoghevirus | tRNAs | 6 | 0.414 | 0.414 |  | 0.829 | 0.985 |
| Yuavirus | Size | 5 | 0.1 | 0.873 |  | 0.985 | 0.985 |
| Yuavirus | CDSs | 5 | -0.791 | 0.111 |  | 0.371 | 0.668 |
| Yuavirus | tRNAs | 5 |  |  | feature constant within genus | | |
| Epaquintavirus | Size | 5 | -0.564 | 0.322 |  | 0.985 | 0.985 |
| Epaquintavirus | CDSs | 5 | 0.0513 | 0.935 |  | 0.935 | 0.985 |
| Epaquintavirus | tRNAs | 5 |  |  | feature constant within genus | | |
| Litunavirus | Size | 5 | -0.0789 | 0.9 |  | 0.985 | 0.985 |
| Litunavirus | CDSs | 5 | 0.703 | 0.185 |  | 0.464 | 0.89 |
| Litunavirus | tRNAs | 5 |  |  | feature constant within genus | | |

**Table S7.** Per-genus Spearman correlations between genome content (genome size, CDS count, tRNA count) and host range, computed within each of the 10 well-sampled genera. p-values are Benjamini-Hochberg adjusted; no test survives correction (24 valid tests).

A.

| **Term** | **Df** | **SumOfSqs** | **F** | **Pr(>F)** |
| --- | --- | --- | --- | --- |
| Model | 19 | 8.13 | 2.93 | 1.0e-04 |
| Residual | 64 | 9.34 |  |  |

B.

| **model** | **Df model** | **SumOfSqs** | **F statistic** | **P value** | **% variance explained** |
| --- | --- | --- | --- | --- | --- |
| Genus alone | 19 | 18.8 | 6.58 | 1.0e-04 | 64 |
| PEQ phylogeny alone | 10 | 12.4 | 5.91 | 1.0e-04 | 42.3 |
| Genus AFTER partialling out PEQ phylogeny | 19 | 8.13 | 2.93 | 1.0e-04 | 27.7 |

**Table S8.** Phage-side distance-based redundancy analysis. (a) Phage genus conditioned on the first 10 PCNM eigenvectors of the PEQ phylogeny. (b) Variance partitioning: genus alone, PCNM alone, and genus unique of PCNM.

A.

| **Term** | **sum_sq** | **df** | **F** | **PR(>F)** | **partial_eta_sq** |
| --- | --- | --- | --- | --- | --- |
| Intercept | 1.528e+04 | 1 | 71.9 | 7.8e-13 | 0.467 |
| C(serotype) | 5604 | 12 | 2.2 | 0.0191 | 0.243 |
| all_prophages | 825.6 | 1 | 3.88 | 0.0521 | 0.0452 |
| Num_Defense | 665.9 | 1 | 3.13 | 0.0804 | 0.0368 |
| Num_Antidefense | 501.8 | 1 | 2.36 | 0.128 | 0.028 |
| n_genes | 26.3 | 1 | 0.124 | 0.726 | 0.00151 |
| Residual | 1.743e+04 | 82 |  |  | 0.5 |

B.

| **metric** | **value** |
| --- | --- |
| R_squared | 0.469 |
| R_squared_adjusted | 0.365 |
| F_statistic | 4.52 |
| F_pvalue | 2.5e-06 |
| df_model | 16 |
| df_residual | 82 |
| n_observations | 99 |
| AIC | 826.8 |
| BIC | 871 |

**Table S9.** Type III ANOVA on per-strain susceptibility count. (a) Per-predictor terms (partial η², F, p). (b) Overall model fit (R², adjusted R², F, AIC, BIC).

|  | **Df** | **SumOfSqs** | **R2** | **F** | **Pr(>F)** |
| --- | --- | --- | --- | --- | --- |
| serotype | 12 | 7.83 | 0.283 | 2.91 | 1.0e-04 |
| Defense | 1 | 0.8 | 0.0289 | 3.56 | 1.0e-04 |
| Antidefense | 1 | 0.543 | 0.0196 | 2.42 | 0.0043 |
| all_prophages | 1 | 0.486 | 0.0175 | 2.16 | 0.0087 |
| PCNM1 | 1 | 0.314 | 0.0113 | 1.4 | 0.121 |
| PCNM2 | 1 | 0.211 | 0.00761 | 0.939 | 0.512 |
| PCNM3 | 1 | 0.277 | 0.00999 | 1.23 | 0.211 |
| PCNM4 | 1 | 0.227 | 0.00819 | 1.01 | 0.426 |
| PCNM5 | 1 | 0.408 | 0.0147 | 1.82 | 0.0275 |
| Residual | 74 | 16.6 | 0.6 | NA | NA |
| Total | 94 | 27.7 | 1 | NA | NA |

**Table S10.** Phylogeny-controlled PERMANOVA on Jaccard distances of per-strain infection profiles, full model (serotype, defense, anti-defense, prophage burden, PCNM1–5).

| **comparison** | **H** | **k** | **N** | **eta_sq** | **epsilon_sq** | **p_value** |
| --- | --- | --- | --- | --- | --- | --- |
| host_range_by_genus | 73.9 | 14 | 88 | 0.823 | 0.85 | 1.5e-10 |
| susceptibility_by_serotype | 31.6 | 13 | 99 | 0.228 | 0.322 | 0.0016 |

**Table S11.** Effect sizes (η² and ε²) for the two main Kruskal-Wallis tests: host range by phage genus, and strain susceptibility by O-antigen serotype.

| **Model** | **Term** | **Df** | **SumOfSqs** | **R2** | **F** | **Pr(>F)** |
| --- | --- | --- | --- | --- | --- | --- |
| Serotype-only model | serotype | 12 | 7.83 | 0.283 | 2.74 | 1.0e-04 |
| Serotype-only model | PCNM1 | 1 | 0.312 | 0.0113 | 1.31 | 0.172 |
| Serotype-only model | PCNM2 | 1 | 0.263 | 0.00948 | 1.1 | 0.317 |
| Serotype-only model | PCNM3 | 1 | 0.25 | 0.00902 | 1.05 | 0.381 |
| Serotype-only model | PCNM4 | 1 | 0.252 | 0.00908 | 1.06 | 0.38 |
| Serotype-only model | PCNM5 | 1 | 0.465 | 0.0168 | 1.95 | 0.0136 |
| Serotype-only model | Residual | 77 | 18.3 | 0.662 |  |  |
| Serotype-only model | Total | 94 | 27.7 | 1 |  |  |
| Defense-only model | Defense | 1 | 1.4 | 0.0504 | 5.08 | 1.0e-04 |
| Defense-only model | PCNM1 | 1 | 0.585 | 0.0211 | 2.13 | 0.01 |
| Defense-only model | PCNM2 | 1 | 0.294 | 0.0106 | 1.07 | 0.354 |
| Defense-only model | PCNM3 | 1 | 0.477 | 0.0172 | 1.73 | 0.0353 |
| Defense-only model | PCNM4 | 1 | 0.283 | 0.0102 | 1.03 | 0.369 |
| Defense-only model | PCNM5 | 1 | 0.447 | 0.0161 | 1.63 | 0.064 |
| Defense-only model | Residual | 88 | 24.2 | 0.874 |  |  |
| Defense-only model | Total | 94 | 27.7 | 1 |  |  |
| Anti-defense-only model | Antidefense | 1 | 0.751 | 0.0271 | 2.67 | 0.002 |
| Anti-defense-only model | PCNM1 | 1 | 0.582 | 0.021 | 2.07 | 0.0105 |
| Anti-defense-only model | PCNM2 | 1 | 0.299 | 0.0108 | 1.06 | 0.375 |
| Anti-defense-only model | PCNM3 | 1 | 0.497 | 0.0179 | 1.77 | 0.0323 |
| Anti-defense-only model | PCNM4 | 1 | 0.293 | 0.0106 | 1.04 | 0.364 |
| Anti-defense-only model | PCNM5 | 1 | 0.512 | 0.0185 | 1.82 | 0.0271 |
| Anti-defense-only model | Residual | 88 | 24.8 | 0.894 |  |  |
| Anti-defense-only model | Total | 94 | 27.7 | 1 |  |  |
| Prophage-only model | all_prophages | 1 | 1.14 | 0.041 | 4.1 | 1.0e-04 |
| Prophage-only model | PCNM1 | 1 | 0.603 | 0.0218 | 2.18 | 0.0064 |
| Prophage-only model | PCNM2 | 1 | 0.286 | 0.0103 | 1.03 | 0.395 |
| Prophage-only model | PCNM3 | 1 | 0.463 | 0.0167 | 1.67 | 0.046 |
| Prophage-only model | PCNM4 | 1 | 0.287 | 0.0103 | 1.03 | 0.383 |
| Prophage-only model | PCNM5 | 1 | 0.529 | 0.0191 | 1.91 | 0.0178 |
| Prophage-only model | Residual | 88 | 24.4 | 0.881 |  |  |
| Prophage-only model | Total | 94 | 27.7 | 1 |  |  |

**Table S12.** Per-feature phylogeny-controlled PERMANOVA (serotype, defense, anti-defense, prophage each modeled alone with PCNM1–5)

| **trait** | **lambda** | **logL** | **logL0** | **LR_p_value** | **n** | **significant** |
| --- | --- | --- | --- | --- | --- | --- |
| susceptibility_count | 0.478 | -461.8 | -463 | 0.119 | 95 | FALSE |
| defense_count | 0.451 | -292.4 | -292.8 | 0.362 | 95 | FALSE |
| antidefense_count | 7.3e-05 | -225.3 | -225.3 | 1 | 95 | FALSE |
| prophage_count | 7.3e-05 | -307.4 | -307.4 | 1 | 95 | FALSE |

**Table S13.** Pagel's λ tests for phylogenetic signal of strain-level traits on the marker-gene phylogeny (n = 95 strains). Susceptibility (λ = 0.48) and defense count (λ = 0.45) show moderate but non-significant phylogenetic structure; anti-defense count and prophage burden show no signal (λ ≈ 0). No trait reached LR-test significance.

| **method** | **Top k** | **Susceptibility quartile** | **Cocktail success %** | **N strain iterations** |
| --- | --- | --- | --- | --- |
| GenoPHI | 3 | Q1 (low) | 65.8 | 38 |
| Promiscuity baseline | 3 | Q1 (low) | 57.9 | 38 |
| Greedy set-cover | 3 | Q1 (low) | 47.8 | 46 |
| Curated baseline | 3 | Q1 (low) | 47.4 | 38 |
| Genus+Serotype | 3 | Q1 (low) | 31.6 | 38 |
| Curated baseline | 3 | Q2 | 94 | 50 |
| Greedy set-cover | 3 | Q2 | 90 | 50 |
| GenoPHI | 3 | Q2 | 88 | 50 |
| Promiscuity baseline | 3 | Q2 | 80 | 50 |
| Genus+Serotype | 3 | Q2 | 76 | 50 |
| Curated baseline | 3 | Q3 | 100 | 48 |
| Promiscuity baseline | 3 | Q3 | 100 | 50 |
| Genus+Serotype | 3 | Q3 | 100 | 50 |
| Greedy set-cover | 3 | Q3 | 100 | 50 |
| GenoPHI | 3 | Q3 | 98 | 50 |
| Curated baseline | 3 | Q4 (high) | 100 | 52 |
| Promiscuity baseline | 3 | Q4 (high) | 100 | 52 |
| GenoPHI | 3 | Q4 (high) | 100 | 52 |
| Genus+Serotype | 3 | Q4 (high) | 100 | 52 |
| Greedy set-cover | 3 | Q4 (high) | 100 | 52 |
| Promiscuity baseline | 5 | Q1 (low) | 78.9 | 38 |
| GenoPHI | 5 | Q1 (low) | 78.9 | 38 |
| Greedy set-cover | 5 | Q1 (low) | 65.2 | 46 |
| Curated baseline | 5 | Q1 (low) | 55.3 | 38 |
| Genus+Serotype | 5 | Q1 (low) | 44.7 | 38 |
| Curated baseline | 5 | Q2 | 96 | 50 |
| Promiscuity baseline | 5 | Q2 | 96 | 50 |
| Greedy set-cover | 5 | Q2 | 96 | 50 |
| GenoPHI | 5 | Q2 | 94 | 50 |
| Genus+Serotype | 5 | Q2 | 90 | 50 |
| Curated baseline | 5 | Q3 | 100 | 48 |
| Promiscuity baseline | 5 | Q3 | 100 | 50 |
| GenoPHI | 5 | Q3 | 100 | 50 |
| Genus+Serotype | 5 | Q3 | 100 | 50 |
| Greedy set-cover | 5 | Q3 | 100 | 50 |
| Curated baseline | 5 | Q4 (high) | 100 | 52 |
| Promiscuity baseline | 5 | Q4 (high) | 100 | 52 |
| GenoPHI | 5 | Q4 (high) | 100 | 52 |
| Genus+Serotype | 5 | Q4 (high) | 100 | 52 |
| Greedy set-cover | 5 | Q4 (high) | 100 | 52 |

**Table S14.** Cocktail success rate by susceptibility quartile. Top-3 and top-5 cocktail success for the three deployable models and two non-ML baselines, stratified by full-matrix susceptibility quartile (Q1 low – Q4 high). n = held-out strain evaluations across cross-validation folds.

| **Analysis** | **PCNM_variant** | **n_PCNM** | **Focal predictor** | **Partial R2** | **F** | **p_value** |
| --- | --- | --- | --- | --- | --- | --- |
| Phage dbRDA | PCNM 1-5 | 5 | Genus | 0.387 | 3.72 | 1.0e-04 |
| Phage dbRDA | PCNM 1-10 | 10 | Genus | 0.27 | 2.64 | 1.0e-04 |
| Phage dbRDA | PCNM 1-20 | 20 | Genus | 0.227 | 2.33 | 1.0e-04 |
| Phage dbRDA | forward selection (12) | 12 | Genus | 0.245 | 2.41 | 1.0e-04 |
| Strain PERMANOVA | PCNM 1-5 | 5 | serotype | 0.24 | 2.49 | 1.0e-04 |
| Strain PERMANOVA | PCNM 1-5 | 5 | Defense | 0.0203 | 2.53 | 0.0033 |
| Strain PERMANOVA | PCNM 1-5 | 5 | Antidefense | 0.0176 | 2.2 | 0.0145 |
| Strain PERMANOVA | PCNM 1-5 | 5 | all_prophages | 0.0198 | 2.47 | 0.004 |
| Strain PERMANOVA | PCNM 1-10 | 10 | serotype | 0.23 | 2.35 | 1.0e-04 |
| Strain PERMANOVA | PCNM 1-10 | 10 | Defense | 0.0203 | 2.5 | 0.0046 |
| Strain PERMANOVA | PCNM 1-10 | 10 | Antidefense | 0.0167 | 2.05 | 0.0187 |
| Strain PERMANOVA | PCNM 1-10 | 10 | all_prophages | 0.0171 | 2.11 | 0.0152 |
| Strain PERMANOVA | PCNM 1-20 | 20 | serotype | 0.187 | 2 | 1.0e-04 |
| Strain PERMANOVA | PCNM 1-20 | 20 | Defense | 0.0202 | 2.59 | 0.0026 |
| Strain PERMANOVA | PCNM 1-20 | 20 | Antidefense | 0.00963 | 1.24 | 0.235 |
| Strain PERMANOVA | PCNM 1-20 | 20 | all_prophages | 0.0118 | 1.51 | 0.101 |
| Strain PERMANOVA | forward selection (7) | 7 | serotype | 0.212 | 2.28 | 1.0e-04 |
| Strain PERMANOVA | forward selection (7) | 7 | Defense | 0.0222 | 2.88 | 0.0011 |
| Strain PERMANOVA | forward selection (7) | 7 | Antidefense | 0.0192 | 2.49 | 0.0071 |
| Strain PERMANOVA | forward selection (7) | 7 | all_prophages | 0.0124 | 1.61 | 0.076 |

**Table S15. PCNM eigenvector-count sensitivity analysis.** The primary phylogeny-controlled analyses (phage-side dbRDA and strain-side PERMANOVA) both condition on the first 10 PCNM eigenvectors. To confirm the conclusions do not depend on this count, both analyses were re-run across PCNM conditioning sets of 5, 10, and 20 eigenvectors and with forward selection (vegan::ordiR2step; 12 axes retained phage-side, 7 strain-side). Genus (phage dbRDA) and O-antigen serotype (strain PERMANOVA) are significant across every variant. Defense-system count is significant across all variants; anti-defense count and prophage burden are significant at PCNM counts ≤ 10 and under forward selection but not when 20 axes are retained. Columns are partial R², pseudo-F, and permutation p-value (9,999 permutations).

| **model** | **reference** | **Top n** | **N paired** | **Mean model** | **Mean reference** | **Median model** | **Median reference** | **Wilcoxon stat** | **Wilcoxon pval** |
| --- | --- | --- | --- | --- | --- | --- | --- | --- | --- |
| Genus+Serotype | GenoPHI | 1 | 20 | 0.643 | 0.725 | 0.667 | 0.778 | 30 | 0.088 |
| Baseline_Ungrouped | GenoPHI | 1 | 20 | 0.723 | 0.725 | 0.778 | 0.778 | 53 | 0.691 |
| SetCover_Greedy | GenoPHI | 1 | 20 | 0.708 | 0.725 | 0.7 | 0.778 | 65.5 | 0.602 |
| CG_Promiscuity | GenoPHI | 1 | 20 | 0.735 | 0.725 | 0.775 | 0.778 | 65 | 0.876 |
| Genus+Serotype | GenoPHI | 3 | 20 | 0.802 | 0.887 | 0.8 | 0.9 | 7.5 | 0.00769 |
| Baseline_Ungrouped | GenoPHI | 3 | 20 | 0.879 | 0.887 | 0.9 | 0.9 | 14.5 | 0.622 |
| SetCover_Greedy | GenoPHI | 3 | 20 | 0.854 | 0.887 | 0.9 | 0.9 | 17.5 | 0.161 |
| CG_Promiscuity | GenoPHI | 3 | 20 | 0.926 | 0.887 | 0.95 | 0.9 | 14 | 0.162 |
| Genus+Serotype | GenoPHI | 5 | 20 | 0.863 | 0.943 | 0.9 | 0.95 | 7 | 0.0117 |
| Baseline_Ungrouped | GenoPHI | 5 | 20 | 0.9 | 0.943 | 0.9 | 0.95 | 7.5 | 0.073 |
| SetCover_Greedy | GenoPHI | 5 | 20 | 0.909 | 0.943 | 0.9 | 0.95 | 14 | 0.0854 |
| CG_Promiscuity | GenoPHI | 5 | 20 | 0.948 | 0.943 | 1 | 0.95 | 15 | 0.673 |

**Table S16.** Paired cocktail-success comparisons against GenoPHI. Top-1/3/5 success rates and paired Wilcoxon tests for Genus+Serotype, curated baseline, greedy set-cover, and the promiscuity baseline.

| **Cocktail generation approach** | **Genera** | **Phages** | **No. of strains infected with score** | | **Sum** | **Infection score in PAO1** | **Genome size (kb)** | **Blastn comparison** |
| --- | --- | --- | --- | --- | --- | --- | --- | --- |
|  |  |  | **1** | **2** |  |  |  |  |
| Classical | Phikzvirus | phiKZ | 34 | 33 | 67 | 2 | 281428 |  |
|  | Phikmvvirus | phiKMV | 6 | 16 | 22 | 2 | 42551 |  |
|  | Pakpunavirus | PAO1_MA12 | 8 | 40 | 48 | 2 | 91870 |  |
| Machine learning | Phikzvirus | DP4 | 32 | 37 | 69 | 2 | 282257 | 97% query coverage, 98.6% identity to phiKZ |
|  | Nankokuvirus | Ab03 | 17 | 21 | 38 | 2 | 85475 |  |
|  | Phikmvvirus | PAO1_MA13 | 13 | 43 | 56 | 2 | 43121 | 91% query coverage, 93.24 % identity to phiKMV |

**Table S17. Phages used to test cocktail efficacy in the murine model.** Two three-phage cocktails (CG, ML); for each constituent phage, the number of panel strains infected at score 1 and score 2, their sum, PAO1 infection score, genome size, and — for the ML phages — BLASTn comparison to the genomically nearest CG phage. All six phages infect PAO1 at score 2.

| **Experiment** | **SM (%)** | **CG (%)** | **ML (%)** |
| --- | --- | --- | --- |
| Day 0 | 0 ± 0 (n=16) | 0 ± 0 (n=16) | 0 ± 0 (n=16) |
| Day 1 | 1.3 ± 8 (n=16) | 9.6 ± 4.1 (n=16) | 4.3 ± 6.8 (n=16) |
| Day 3 | 23.1 ± 6.1 (n=5) | 22 ± 4.6 (n=14) | 18.1 ± 5.2 (n=16) |

**Table S18.** Percent wound closure over time. Wound area was measured at the indicated time points and expressed as percent closure relative to baseline (Day 0). Data are presented as mean ± SEM. Sample size (n) represents the number of wounds that could be reliably measured at each time point. On Day 3, only 5 of 16 wounds were measurable because the remaining wounds were obscured by opaque yellowish wound secretion, preventing accurate wound-area quantification. Therefore, the Day 3 SM value is provided for descriptive purposes only and should not be interpreted as representative of the full group.

| **metric** | **metric_label** | **condition** | **n** | **mean** | **sd** | **min** | **q25** | **median** | **q75** | **max** |
| --- | --- | --- | --- | --- | --- | --- | --- | --- | --- | --- |
| total_reads | Total reads (all, incl. host) | ML | 15 | 8654 | 6927 | 1657 | 3230 | 6589 | 1.287e+04 | 21529 |
| total_reads | Total reads (all, incl. host) | CG | 13 | 6787 | 5077 | 1155 | 2373 | 4322 | 11028 | 14815 |
| total_reads | Total reads (all, incl. host) | SM | 16 | 6802 | 4957 | 1020 | 2424 | 6492 | 9944 | 17307 |
| microbial_reads | Microbial reads (host removed) | ML | 15 | 152.3 | 203.7 | 0 | 13.5 | 46 | 159 | 592 |
| microbial_reads | Microbial reads (host removed) | CG | 13 | 831.5 | 863.4 | 41 | 175 | 497 | 1556 | 2427 |
| microbial_reads | Microbial reads (host removed) | SM | 16 | 5110 | 4099 | 17 | 1203 | 4820 | 7158 | 13519 |
| host_reads | Host reads | ML | 15 | 8502 | 6986 | 1562 | 3178 | 6454 | 12850 | 21484 |
| host_reads | Host reads | CG | 13 | 5955 | 5253 | 530 | 1550 | 4040 | 9209 | 14640 |
| host_reads | Host reads | SM | 16 | 1692 | 2604 | 151 | 422.5 | 916 | 1721 | 10908 |
| host_frac | Host fraction of reads | ML | 15 | 0.963 | 0.068 | 0.758 | 0.977 | 0.996 | 0.998 | 1 |
| host_frac | Host fraction of reads | CG | 13 | 0.776 | 0.281 | 0.223 | 0.794 | 0.873 | 0.988 | 0.992 |
| host_frac | Host fraction of reads | SM | 16 | 0.308 | 0.287 | 0.0158 | 0.0721 | 0.285 | 0.377 | 0.987 |
| pseudo_reads | Pseudomonas reads | ML | 15 | 86.3 | 174.4 | 0 | 2.5 | 13 | 22 | 513 |
| pseudo_reads | Pseudomonas reads | CG | 13 | 682.7 | 906.7 | 13 | 41 | 137 | 1422 | 2380 |
| pseudo_reads | Pseudomonas reads | SM | 16 | 5083 | 4069 | 17 | 1196 | 4817 | 7126 | 13383 |
| pseudo_frac_total | Pseudomonas / total reads | ML | 15 | 0.0251 | 0.0648 | 0 | 3.2e-04 | 0.00159 | 0.00506 | 0.238 |
| pseudo_frac_total | Pseudomonas / total reads | CG | 13 | 0.182 | 0.282 | 0.0014 | 0.00925 | 0.025 | 0.196 | 0.759 |
| pseudo_frac_total | Pseudomonas / total reads | SM | 16 | 0.689 | 0.285 | 0.0133 | 0.622 | 0.715 | 0.92 | 0.983 |
| pseudo_frac_microbial | Pseudomonas / microbial reads | ML | 14 | 0.499 | 0.445 | 0 | 0.0565 | 0.363 | 0.995 | 1 |
| pseudo_frac_microbial | Pseudomonas / microbial reads | CG | 13 | 0.643 | 0.373 | 0.0746 | 0.241 | 0.783 | 0.978 | 1 |
| pseudo_frac_microbial | Pseudomonas / microbial reads | SM | 16 | 0.996 | 0.00466 | 0.988 | 0.991 | 0.997 | 1 | 1 |
| richness_all | Zotu richness (incl. host) | ML | 15 | 4.13 | 2 | 1 | 3 | 4 | 5.5 | 8 |
| richness_all | Zotu richness (incl. host) | CG | 13 | 4.85 | 1.72 | 3 | 4 | 4 | 5 | 10 |
| richness_all | Zotu richness (incl. host) | SM | 16 | 4.38 | 1.59 | 3 | 3 | 4 | 4.25 | 8 |
| richness_microbial | Zotu richness (microbial only) | ML | 15 | 3.13 | 2 | 0 | 2 | 3 | 4.5 | 7 |
| richness_microbial | Zotu richness (microbial only) | CG | 13 | 3.85 | 1.72 | 2 | 3 | 3 | 4 | 9 |
| richness_microbial | Zotu richness (microbial only) | SM | 16 | 3.38 | 1.59 | 2 | 2 | 3 | 3.25 | 7 |

**Table S19.** Per-group summary statistics for nine 16S metrics (ML, CG, SM): total reads, host reads, microbial reads, Pseudomonas reads, Pseudomonas fractions, ZOTU richness, host fraction.

| **metric** | **metric_label** | **n_total** | **n_groups** | **H** | **df** | **p_value** | **q_value_fdr** |
| --- | --- | --- | --- | --- | --- | --- | --- |
| total_reads | Total reads (all, incl. host) | 44 | 3 | 0.648 | 2 | 0.723 | 0.723 |
| microbial_reads | Microbial reads (host removed) | 44 | 3 | 24.3 | 2 | 5.4e-06 | 1.2e-05 |
| host_reads | Host reads | 44 | 3 | 17 | 2 | 2.0e-04 | 3.6e-04 |
| host_frac | Host fraction of reads | 44 | 3 | 26.2 | 2 | 2.0e-06 | 6.1e-06 |
| pseudo_reads | Pseudomonas reads | 44 | 3 | 27.1 | 2 | 1.3e-06 | 5.8e-06 |
| pseudo_frac_total | Pseudomonas / total reads | 44 | 3 | 28.8 | 2 | 5.5e-07 | 5.0e-06 |
| pseudo_frac_microbial | Pseudomonas / microbial reads | 43 | 3 | 14.2 | 2 | 8.3e-04 | 0.00125 |
| richness_all | Zotu richness (incl. host) | 44 | 3 | 1.91 | 2 | 0.385 | 0.433 |
| richness_microbial | Zotu richness (microbial only) | 44 | 3 | 1.91 | 2 | 0.385 | 0.433 |

**Table S20**. Kruskal-Wallis tests across ML/CG/SM for each 16S metric, with Benjamini-Hochberg-corrected q-values.

### Supplementary Dataset

**Dataset S1.** Per-strain prophage counts stratified by CheckV completeness threshold (99 strains).

**Dataset S2.** Full 95-phage × 99-strain interaction matrix (9,405 scored interactions).

**Dataset S3.** Per-fold, per-strain, and per-phage AUROC/AUPR/MCC for all models, with paired-test statistics.

**Dataset S4.** Per-tercile model performance across four difficulty axes (phylogenetic novelty, phenotypic divergence, marginal difficulty, full-matrix degree).

**Dataset S5.** Full quartile × variant × top-k cocktail-success table (144 rows; backs Table S14).

**Dataset S6.** Per-strain counts of 118 defense-system families.

**Dataset S7.** Predicted O-antigen serotype and OSA coverage per strain.

**Dataset S8.** Per-prophage geNomad predictions (coordinates, completeness, taxonomy).

**Dataset S9.** Prophage counts by viral realm and per-strain gene counts.

**References**

1.	Lebreton, F. *et al.* A panel of diverse clinical isolates for research and development. *JAC Antimicrob Resist* **3**, dlab179 (2021).
